# Insulin regulates lymphatic endothelial function via palmitoylation

**DOI:** 10.1101/2024.05.06.592341

**Authors:** Silvia Gonzalez-Nieves, Xiaochao Wei, Jay McQuillan, Qiang Zhang, Jinsong Zhang, Reagan M. McGuffee, David A. Ford, Latisha Love-Gregory, Nada A. Abumrad, Andrew E Gelman, Clay F. Semenkovich, Vincenza Cifarelli

**Affiliations:** Department of Pharmacology and Physiology, Saint Louis University School of Medicine, St. Louis, MO, USA; Division of Endocrinology Metabolism and Lipid Research, Department of Medicine, Washington University, St. Louis, MO, USA; Department of Biochemistry and Molecular Biology, Saint Louis University School of Medicine, St. Louis, MO, USA; Department of Medicine, Washington University, St. Louis, MO, USA; Department of Cell Biology and Physiology, Washington University, St. Louis, MO, USA; Department of Surgery Washington University, St. Louis, MO, USA

**Keywords:** insulin, LECs, palmitoylation, CD36, proteomics, claudin 5, p120-catenin

## Abstract

Lipid metabolism plays a critical role in lymphatic endothelial cell (LEC) development and maintenance. Altered lipid metabolism is associated with loss of lymphatic vessel integrity, which compromises organ function, protective immunity, and metabolic health. However, the role of lipid metabolism in LEC function is not well understood.

Insulin is a key regulator of lipid metabolism and protein palmitoylation, the reversible post-translational protein modification by palmitate that affects protein stability, trafficking, protein-protein, and protein-membrane interactions. Human LECs are highly sensitive to insulin and can develop insulin resistance *in vitro*, but whether insulin regulates LEC protein palmitoylation and function is unknown.

To examine the role of palmitoylation in LEC function, we generated the first palmitoylation proteomics profile in human LECs, validated insulin regulated targets and profiled differences in palmitoylation between lymphatic and blood endothelial cells. Palmitoylation occurred primarily in proteins involved in LEC vesicular or membrane trafficking, translation initiation, and in those found in membrane rafts. Insulin enriched palmitoylation of LEC proteins involved in GTPase signaling, ubiquitination, and junctional anchoring. We also determined that the long-chain fatty acid receptor CD36 mediates optimal lymphatic palmitoylation. CD36 silencing in LECs doubled palmitoylation targets involving proteins related to inflammation and neutrophil degranulation contributing to an *activated* inflamed endothelium. These results suggest that the coordination of the process of palmitoylation is critical for normal lymphatic endothelial function.

## INTRODUCTION

Lymphatic endothelial cells (LECs) preferentially utilize fatty acid β-oxidation (FAO) to grow and maintain lymphatic vessel identity through epigenetic regulation of the essential transcription factor prospero homeobox 1, PROX1 (1, 2). Accordingly, FAO is driven by robust expression in lymphatic vessels of genes involved in lipid transport and metabolism (3, 4). However, the role of lipids in lymphatic function is poorly understood despite reports that an abnormal lipid milieu exists in diseases with impaired lymphatic function including acquired forms of lymphedema (5), obesity (6, 7), hypercholesterolemia (8, 9), congestive heart failure (10, 11), and peripheral arterial/venous disease (12).

Lipid can affect cellular function via posttranslational modifications of proteins namely myristoylation, prenylation, and acylation (13). Protein lipidation increases the hydrophobicity of proteins, resulting in changes in protein conformation, stability, membrane association, trafficking, and binding to co-factors (13). Among the different types of lipidation, palmitoylation is the only reversible post-translational lipid modification and it usually links the long chain FA palmitate to a cysteine amino acid. Reversible palmitoylation confers spatiotemporal control of protein function by modulating protein stability, trafficking, and activity, as well as protein–protein and membrane–protein associations (14, 15). Palmitoylation is catalyzed by ∼20 aspartate-histidine-histidine-cysteine acyltransferases (DHHC) (16), many expressed in endothelial cells (17). Depalmitoylation is performed by a family of enzymes that includes acyl-protein thioesterase 1 and 2 (APT-1, 2) (18) and α/β-Hydrolase Domain proteins (ABHD) 17-A, B and C (19). Altered palmitoylation has been reported in pathological vascular remodeling such as peripheral artery disease (20–22) but the role of palmitoylation in lymphatics is not known.

Insulin is a strong regulator of lipid metabolism (23) and of protein palmitoylation (22). Human LECs are highly sensitive to insulin stimulation *in vitro* initiating a PI3K/AKT signaling cascade at lower insulin doses (e.g., 2.5 nM) than in blood endothelial cells (24) (BECs). Hyperglycemia and hyperinsulinemia-induced insulin resistance in cultured LECs blunt activation of PI3K/AKT and are associated with disruption of the integrity of the LEC monolayer and inflammation (25). *In vitro* findings are in line with studies in preclinical models of obesity, metabolic syndrome, and diabetes (11, 26, 27) reporting impaired lymphatic function and lost vessel integrity. However, targets of insulin action in lymphatics are largely unknown and protein palmitoylation is unstudied. Our study documents the first quantitative proteomics palmitoylation profile in human LECs; validates insulin regulated targets in LECs and interrogates role of the long-chain FA transporter CD36 in optimal lymphatic protein palmitoylation. Finally, we compare insulin induced palmitoylation profiles between BECs and LECs.

## METHODS

### Cell culture and treatments

Primary human dermal lymphatic endothelial cells (hLECs) were obtained from PromoCell and grown in EGM-2MV (microvascular endothelial cell growth medium-2) medium from Lonza Bioscience as previously described (3). hLECs were used between passages 2 and 8 for experiments. Cells were serum starved (0.5% FBS) for 1 h and treated with or without insulin (100 nM; Sigma, St. Louis) for 6 h at 37°C for proteomics experiments. Lipopolysaccharide (20ng/mL) treatment was utilized as a positive control to disrupt barrier integrity. For siRNA treatment, the LECs were grown to 60–80% confluence and transfected for 48 h with either control or CD36 Silencer® pre-designed siRNA as per the manufacturer’s instructions (ThermoFisher Scientific, Waltham, MA, USA). Cells were then serum starved for 1 h before insulin (100 nM) or media or 2-BP (20 μM) treatment.

### Acyl-RAC assays for protein palmitoylation

S-palmitoylated proteins were purified using a modified acyl-RAC (resin assisted capture) assay as previously described (20). Cells were homogenized in lysis buffer (150 mM NaCl, 50 mM Tris, 5 mM EDTA, 2% Triton-X-100, 0.2 mM HDSF, pH 7.4, and protease inhibitor cocktail from Roche). Following sonication on ice, protein concentrations were determined by BCA protein assay kits (Thermo), and the same amounts of protein lysates were diluted to 2.5% SDS with 0.1% MMTS (S-Methyl methanethiosulfonate), and incubated at 37°C for 20 min. Proteins were then precipitated with ice-cold acetone (1:3 v/v protein solution: acetone) at -20°C for 30 min. Protein precipitates were obtained by centrifuging at 12000 rpm for 10 min, washed 3 times with cold 70% acetone, and air dried. Pellets were resuspended with binding buffer (100 mM HEPES, 1 mM EDTA, 1% SDS, pH 7.4, and protease inhibitor cocktail from Roche) by vortexing for 20 min at room temperature. Solubilized fractions were obtained by centrifuging at 14000rpm for 5 min, and supernatants were then aliquoted into 2 tubes containing thiopropyl sepharose beads (GE Healthcare Life Sciences), with or without freshly-prepared HA (hydroxylamine 0.8 M, pH 7.4), and rotated at room temperature for 2 h. Beads were washed 5 times with binding buffer, and enriched proteins were eluted with binding buffer containing 50 mM DTT (20 min incubation at room temperature). Palmitoylated proteins were analyzed by Western blotting using Licor infrared fluorescence imaging.

### Proteomics

#### Peptide preparation

Peptides were prepared from eluates as previously described (29). Samples were reduced with 100 mM dithiothreitol at 95 °C for 15 min, mixed with 200 µl 100 mM Tris-HCL buffer, pH 8.5 containing 8 M urea (all from Sigma), transferred to the top of a 30,000 molecular weight cut-off filter (Millipore, part# MRCF0R030) and spun at 10,000 rcf for 10 minutes. An additional 300 µl of UA buffer was added and the filter was spun at 10,000 rcf for 10 minutes. The flow through was discarded and the proteins were alkylated using 100 µl of 50 mM Iodoacetamide (IAM) in UA buffer. IAM in UA buffer was added to the top chamber of the filtration unit. The samples were gyrated at 550 rpm using a Thermomixer (Eppendorf) at room temperature for 30 minutes in the dark. The filter was spun at 10,000 rcf for 10 minutes and the flow through discarded. Unreacted IAM was washed through the filter with two additions of UA buffer, and centrifugation at 10,000 rcf for 10 minutes after each buffer addition. The UA buffer was exchanged with digestion buffer (DB), (50 mM ammonium bicarbonate buffer). Two sequential additions of DB (200 µl) with centrifugation after each addition to the top chamber was performed. The filters were transferred to a new collection tube and samples digested with a combination of LysC (1 mAU per filter) and trypsin (1:50 wt/wt) in DB buffer on top of the filter for two hours and overnight at 37 °C. The filters were spun at 14,000 rcf for 15 minutes to collect the peptides in the flow through. The filter was washed with 50 µl 100mM ammonium bicarbonate buffer and the wash collected with the peptides. Residual detergent was removed by ethyl acetate extraction (30). In preparation for desalting, peptides were acidified to pH=2 with 1% trifluoroacetic acid (TFA) final concentration. Peptides were desalted using stage tips (C18) (31), eluted with 60 µl of 60% (vol/vol) of MeCN in 0.1% (vol/vol) FA, and dried in a Speed-Vac (Thermo Scientific, Model No. Savant DNA 120 concentrator). The peptides were dissolved in 100 µl of 1% MeCN in water. An aliquot (10%) was removed for quantification using the Pierce Quantitative Fluorometric Peptide Assay kit. The remaining peptides were transferred to autosampler vials, dried and stored at -80°C for LC-MS analysis.

#### Ultra-high performance liquid chromatography mass spectrometry – timsTOF

The unlabeled peptides were analyzed using trapped ion mobility time-of-flight mass spectrometry(32). Peptides were separated using a *nano-ELUTE* ® chromatograph (Bruker Daltonics. Bremen, Germany) interfaced to a timsTOF Pro mass spectrometer (Bruker Daltonics) with a modified nano-electrospray source (CaptiveSpray, Bruker Daltonics). The mass spectrometer was operated in PASEF mode(32). The samples in 2 µl of 1% (vol/vol) FA were injected onto a 75 µm i.d. × 25 cm Aurora Series column with a CSI emitter (Ionopticks). The column temperature was set to 50 °C. The column was equilibrated using constant pressure (800 bar) with 8 column volumes of solvent A (0.1% (vol/vol) FA). Sample loading was performed at constant pressure (800 bar) at a volume of 1 sample pick-up volume plus 2 µl. The peptides were eluted using one column separation mode with a flow rate of 300 nL/min and using solvents A (0.1% (vol/vol) FA) and B (0.1% (vol/vol) FA/MeCN): solvent A containing 2%B increased to 17% B over 60 min, to 25% B over 30 min, to 37% B over 10 min, to 80% B over 10 min and constant 80% B for 10 min. The MS1 and MS2 spectra were recorded from *m/z* 100 to 1700. The collision energy was ramped stepwise as a function of increasing ion mobility: 52 eV for 0–19% of the ramp time; 47 eV from 19–38%; 42 eV from 38–57%; 37 eV from 57–76%; and 32 eV for the remainder. The TIMS elution voltage was calibrated linearly using the Agilent ESI-L Tuning Mix (*m/z* 622, 922, 1222).

#### MS data analysis

For timsTOF files, data from the mass spectrometer were converted to peak lists using DataAnalysis (version 5.2, Bruker Daltonics). MS2 spectra with parent ion charge states of +2, +3 and +4 were analyzed using Mascot software(33) (Matrix Science, London, UK; version 2.8.0.1) against a concatenated UniProt (ver Feburary 2022) database of human (20,512 entries) and common contaminant proteins (cRAP, version 1.0 Jan. 1st, 2012; 116 entries). Trypsin/P enzyme specificity with a maximum of 4 missed cleavages allowed was used. Label-free “single-shot” LC-MS data from the timsTOF mass spectrometer were searched with a fragment ion mass tolerance of 20 ppm and a parent ion tolerance of 20 ppm. Carbamidomethylation of cysteine was specified in Mascot as a fixed modification. Deamidation of asparagine, formation of pyro-glutamic acid from N-terminal glutamine, acetylation of protein N-terminus, and oxidation of methionine were specified as variable modifications. PSMs were filtered at 1% false-discovery rate (FDR) by searching against a reversed database. A minimum of two peptides with unique sequences, not resulting from missed cleavages, was required for identification of a protein. The processing, quality assurance and analysis of LC-MS data were performed with proteoQ (version 1.7.5.1), part of the Mzion suite(34) software developed with the *tidyverse* approach with open-source software for statistical computing and graphics, R and R Studio. The precursor intensities were converted to logarithmic ratios (base 2), relative to the average precursor intensity across all samples. Within each sample, Dixon’s outlier removals were carried out recursively for peptides with greater than two identifying PSMs. The median of the ratios of PSM that could be assigned to the same peptide was first taken to represent the ratios of the incumbent peptide. The median of the ratios of peptides was then taken to represent the ratios of the inferred protein. To align protein ratios across samples, likelihood functions were first estimated for the log-ratios of proteins using finite mixture modeling, assuming two-component Gaussian mixtures (35). The ratio distributions were then aligned so that the maximum likelihood of log-ratios was centered at zero for each sample. Scaling normalization was performed to standardize the log-ratios of proteins across all samples. To reduce the influence of outliers from either log-ratios or reporter-ion intensities, the values between the 5^th^ and 95^th^ percentile of log-ratios and 5^th^ and 95^th^ percentile of intensity were used in the calculations of standard deviations.

#### Bioinformatic and statistical analysis

Metric multidimensional scaling (MDS) and Principal component analysis (PCA) of protein log2-ratios were performed with the base R function stats:cmdscale and stats:prcomp, respectively. Protein log2-ratios wasvisualized by volcano-plot. Linear modeling was performed using the contrast fit approach in limma to assess the statistical significance of protein abundance differences between indicated groups of contrasts. Adjustments of p-values for multiple comparisons were used with Benjamini-Hochberg (BH) correction.

#### Pathway over-representation analysis

WebGestalt analysis tool kit (36) was used to identify enriched molecular pathways based on protein palmitoylation enrichment in each condition. Gene sets from Gene Ontology biological process and cellular component were utilized to determine affected pathways.

### Cell immunofluorescence imaging

Following treatments as indicated, 100% confluent hLECs were fixed (4% formaldehyde, 10 min), permeabilized (0.1% Triton X-100/PBS, 2 min) and blocked (1% BSA/PBS, 60 min). Primary (overnight at 4°C) and fluorescent-labeled secondary antibody (1 h at room temperature) was added (**Supplemental Table S1**), each followed by PBS washes at room temperature. Cells were then mounted on a coverslip using ProLong™ Gold Anti-fade mounting medium with DAPI (ThermoFisher Scientific) for microscope visualization. To determine rearrangement of the cytoskeleton, phalloidin staining (1:400, AlexaFluor647-Phalloidin, ThermoFisher Scientific) was performed concomitantly with the junction staining following manufacturer’s instructions. Quantification of junctional thickness was assessed in a blinded manner to prevent experimenter bias. ImageJ was used to manually draw lines perpendicularly to VE-cadherin between adjacent cells in each image, 10 fields per replicate. The width of each junction was recorded; the majority (>75%) of junctions were between 7-10 µm in width in control monolayers. Junctions thicker than 7 µm classified as “thick” and those less than 7 µm in width classified as “thin”, and percentages of thin and thick junctions derived from these measurements. Stacks of the images were analyzed in Fiji by JACoP plugin for colocalization analysis. Images were converted to 8-bit and background subtracted using rolling-ball algorithm (r=50). The Mander’s coefficients of VE-cadherin and phalloidin channels were calculated based on the same threshold for the experiment, and data reported as the fraction of VE-cadherin-stained pixels overlapping with phalloidin stain. Two-color intensity profiles were generated in Fiji by drawing straight lines in the overlay images, and then plotted in Prism.

### Transendothelial electrical resistance (TEER)

Integrity of the monolayer was assessed by measuring changes in TEER using an electrical cell-substrate impedance sensing ECIS^91^ (Applied Biophysics, Troy, NY, USA) which was housed in a standard tissue culture incubator maintained at 37°C and 5% CO_2_. hLECs (5x10^4^) were seeded using complete EGM-2MV media into plates with gold-coated microelectrodes (Applied Biophysics catalog #8W10E+) and inspected under the phase contrast microscope to verify that the monolayer was complete and overtly normal. The day of the experiments, cells were serum starved (0.5% FBS) before placing the chamber in the TEER instrument. Impedance was monitored for 180 min until stabilized (plateau) before adding the same volume of media or insulin (100 nM), or insulin and 2-Bromopalmitate (2-BP, 20 μM). Lipopolysaccharide (LPS, 50 ng/mL, which impairs the integrity of the monolayer (37), was used as a positive control. The electric current passing through the endothelial monolayers was measured independently in each well, and continuously in real-time for 18 h. TEER data are presented as resistance (Ohm) measured in each well over time.

### Western blotting

Cells were lysed (30 min, new protocol) in cold Cell Signaling 1X Cell Lysis Buffer (20 mM Tris-HCl (pH 7.5), 150 mM NaCl, 1 mM Na2EDTA, 1 mM EGTA, 1% Triton, 2.5 mM sodium pyrophosphate, 1 mM beta-glycerophosphate, 1 mM Na3VO4, 1 µg/ml leupeptin) containing cOmplete Mini proteinase inhibitors and PhosphoStop. Lysates were cleared (12,500 × g, 15 min) and assayed for protein concentration (ThermoFisher Scientific). Proteins were separated on 4–12% gradient gels and transferred to nitrocellulose membranes (Bio-Rad), blocked (Li-COR Biosciences, NE, USA) 1 h at room temperature before incubation with primary antibodies (**Supplementary Table 1**) overnight at 4 °C. Infrared dye–labeled secondary antibodies (**Supplementary Table 1**) were added for 1 h at room temperature. Protein signals were detected using the Li-Cor Odyssey Infrared (Li-COR Biosciences) and quantified using Image Studio Lite v5.2 Software.

### Proximity Ligation Assay (PLA)

PLA protocol was performed following the manufacturer’s instructions (Sigma). Cells (∼13x10^5^) were seeded into poly-lysine-coated coverslips. After 48 hours of CD36 silencing by siRNA, cells were serum starved for 1 hour, and then treated with media alone or supplemented with insulin (100mM, 6h). Cells were then fixed (4% formaldehyde, 10 mi), permeabilized (0.5% Triton X-100/PBS, 10 min), blocked and incubated with primary antibodies rabbit anti-human CD63 and mouse anti-human integrin β1 (**Supplemental Table S1**) overnight at 4°C. Anti-mouse and anti-rabbit probes were added the next day, followed by ligation and amplification steps. Coverslips were mounted on slides using ProLong™ Gold Anti-fade mounting medium with DAPI (ThermoFisher Scientific) and PLA signal was detected as Texas Red® signal using a Leica CTR6 LED microscope. Signal was quantified as a mean intensity of fluorescence/cell/treatment, and background was subtracted as the mean intensity of fluorescence of five different empty areas for each field using ImageJ software.

### Human subjects, blood collection and genotyping

Recruitment of unrelated African American study participants was conducted at Washington University. All clinical investigation was conducted according to the Declaration of Helsinki principles. All human studies were approved by the Human Research Protection Office Institutional Review Board at Washington University (HRPO, # 201202101). Written informed consent was obtained from participants prior to inclusion in the study. Homozygosity for the G allele (allele frequency ∼9%) of coding SNP rs3111938 (G/G) introduces a stop codon and results in a truncated CD36 protein that is degraded, resulting in complete CD36 deficiency(38). Study participants with type 2 diabetes were excluded as well as individuals reporting statin use. Study subjects were admitted in the morning at the Washington University Clinical Research Unit for vitals. Fasting (overnight) plasma glucose concentration was determined by using the glucose oxidase method (YSI, Inc., Yellow Springs, Ohio). Plasma HbA1c was measured in the Washington University Core Laboratory for Clinical Studies. Genomic DNA was extracted from peripheral blood with a salting-out precipitation procedure (Gentra Puregene Blood kit) and *CD36* SNP rs3211938 detected using a predesigned TaqMan SNP Genotyping Assay (Applied Biosystems) on a 7900HT instrument. CD36 expression was determined on freshly isolated monocytes and/or platelets as previously described (38).

### Flow cytometry

Whole peripheral blood was collected in heparin coated tubes, diluted with an equal volume with Dulbecco’s PBS (ThermoFisher Scientific) with 2% with Fetal Bovine serum (ThermoFisher Scientific), layered carefully over 10 mls of Lymophoprep^TM^ density centrifugation medium (StemCell Technologies, Vancouver, Canada) and centrifuged at 800 x g for 20 minutes at room temperature with the brake off. Granulocyte-erythrocyte pellet was resuspended in ammonia chloride red blood lysis solution (StemCell) on ice for 10 minutes and then at 500 x g for 10 minutes. The resulting granulocyte pellet were resuspended in Dulbecco’s PBS (ThermoFisher Scientific) with 2% with Fetal Bovine serum with F_c_ block (Biolegend, San Diego, California, USA) and stained with monoclonal antibodies specific for CD66b (clone GM2H6), CD11b (clone VIM12), CD15 (clone MEM-158), CD14 (clone Tuk4) and CD62L (clone Dreg-56) (All antibodies obtained from ThermoFisher Scientific). Cells were finally stained with the Zombie vital dye (Biolegend) and data on live neutrophils was collected on a modified FACS-Scan with 488, 633 and 405 nm lasers. Data were analyzed with FlowJo software Version 8.8.7.

### Analysis of published palmitoylation and RNA sequencing data

Palmitoylation profiles from human umbilical vein endothelial cells (HUVEC) were obtained from Wei et colleagues (22). Previously published RNA sequencing data (39) from HUVEC and HDELCs were retrieved from the National Center of Biotechnology Information (NCBI) Gene Expression Omnibus database GSE209855 and GSE212743.

### Statistical Analysis

GraphPad Prism software (version 8.4.3) was used for all statistical analyses and to plot quantitative data. All data shown are means ± standard error mean (SEM). Statistical significance was evaluated using two-sided Student’s t test with a p value of ≤0.05 indicating significant differences. One-way ANOVA with Sidak’s post hoc test was used for multiple comparisons.

## RESULTS

### Global Identification of Palmitoylated Proteins in LECs and Effect of Insulin

Since targets of lymphatic palmitoylation are unknown, we performed an unbiased quantitative proteomic profile of palmitoylated proteins in human dermal lymphatic endothelial cells (hLECs) under unstimulated conditions (baseline) and ± insulin for 6 h to identify putative targets of insulin-regulated palmitoylation. The median half-life of mammalian proteins is estimated at ∼46h (40), thus it is unlikely that steady-state protein levels would be affected within 6 hours. Following treatments, cells were first treated with N-ethylmaleimide (NEM) to modify free thiols, to prevent their subsequent biotinylation, and then with hydroxylamine (HA^+^), which cleaves the thioester bond between palmitate and cysteines, leaving cysteines susceptible to biotin labeling and subsequent pulldown. As a control for the HA step, we generated a minus HA condition, so each sample was HA+ or HA-. Unlabeled peptides were analyzed using trapped ion mobility time-of-flight mass spectrometry (**Supplemental Figure S1**) as described in Methods. Three matched groups of media or insulin treated cell preps were analyzed, and normalized fold changes with p values were generated for individual proteins. Of more than 3,300 identified proteins, 380 proteins were significantly palmitoylated at baseline (p values ≤0.05 and ≥2 peptides) (**Figure 1a**). These included proteins involved in vesicular or membrane trafficking, translation initiation, and membrane rafts, and are grouped based on functional characteristics (**Figure 1c-d**; full list in **Supplemental Table S2**). Insulin enriched palmitoylation of 176 proteins (fold ≥1.2 and ≥2 identifying peptides), 75 of these were shared with unstimulated conditions and included proteins involved in GTPase signaling, ubiquitination, and junction anchoring (**Figure 1e-f** and **Supplemental Table S3**).

**Figure 1.**
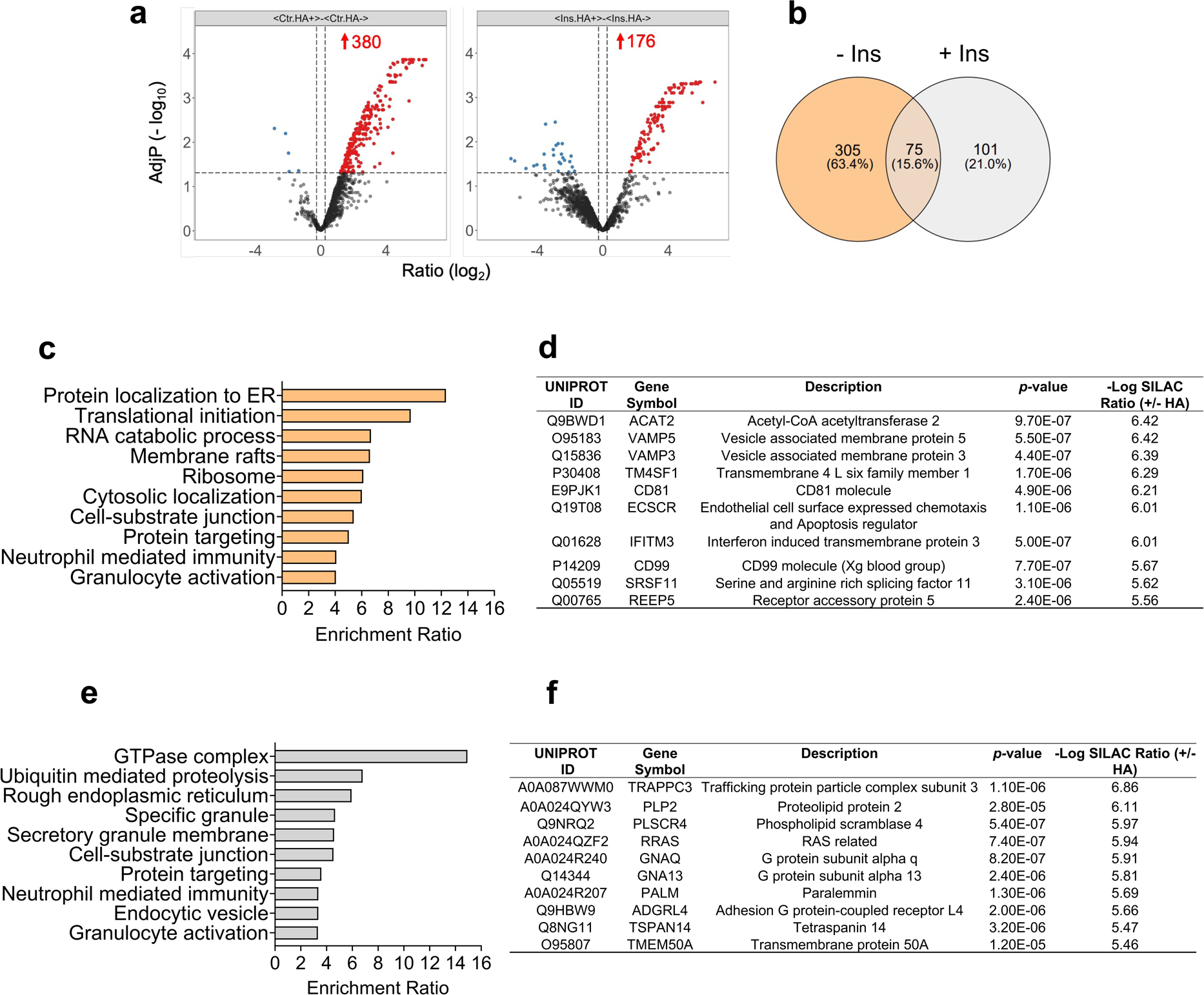
Palmitoylation proteomics in hLECs. (**a**) Volcano plot and (**b**) Venn diagram of palmitoylated targets in media (Ctrl, left panel) and insulin (right panel) treated hLECs. Three matched groups of control and insulin treated cell preps were analyzed and normalized fold changes with P values were generated for individual proteins. (**c-d**) Pathway analysis and list of the of the 10 most upregulated target in hLECs at baseline or (**e-f**) following 6h insulin treatment. Adjustments of p-values for multiple comparisons were used with Benjamini-Hochberg (BH) correction.

We next focused on insulin-mediated regulation of palmitoylation of proteins important for LEC junctions, since insulin resistance associates with impaired lymphatic vessel integrity (25) and leakage (41). Using AcylRac and western blotting, we validated insulin induced palmitoylation of several candidates (42) namely anchoring junction p120-catenin and tight junction claudin 5 (**Figure 2**), both co-localized at the plasma membrane with vascular endothelial (VE)-cadherin (**Supplemental Figure S2a-b**), a gatekeeper of lymphatic integrity (42). Immunofluorescence analysis showed that insulin induced junctional remodeling of p120-catenin at the plasma membrane, which appeared more linearized as compared to unstimulated hLECs (**Figure 2b**). Similarly, claudin 5 staining revealed more homogenous distribution of the tight junction at the plasma membrane in addition to punctate staining in the cytoplasm (**Figure 2d**). Additional targets of insulin-induced palmitoylation are proteins enriched in membrane rafts reported to regulate lymphatic integrity (3, 43, 44), including long-chain FA transporter CD36. Localization of CD36 in plasma membrane rafts was enhanced by insulin treatment (**Figure 2e**) and this was supported by more CD36 localization in the plasma membrane compared to unstimulated hLECs (**Figure 2f**).

**Figure 2.**
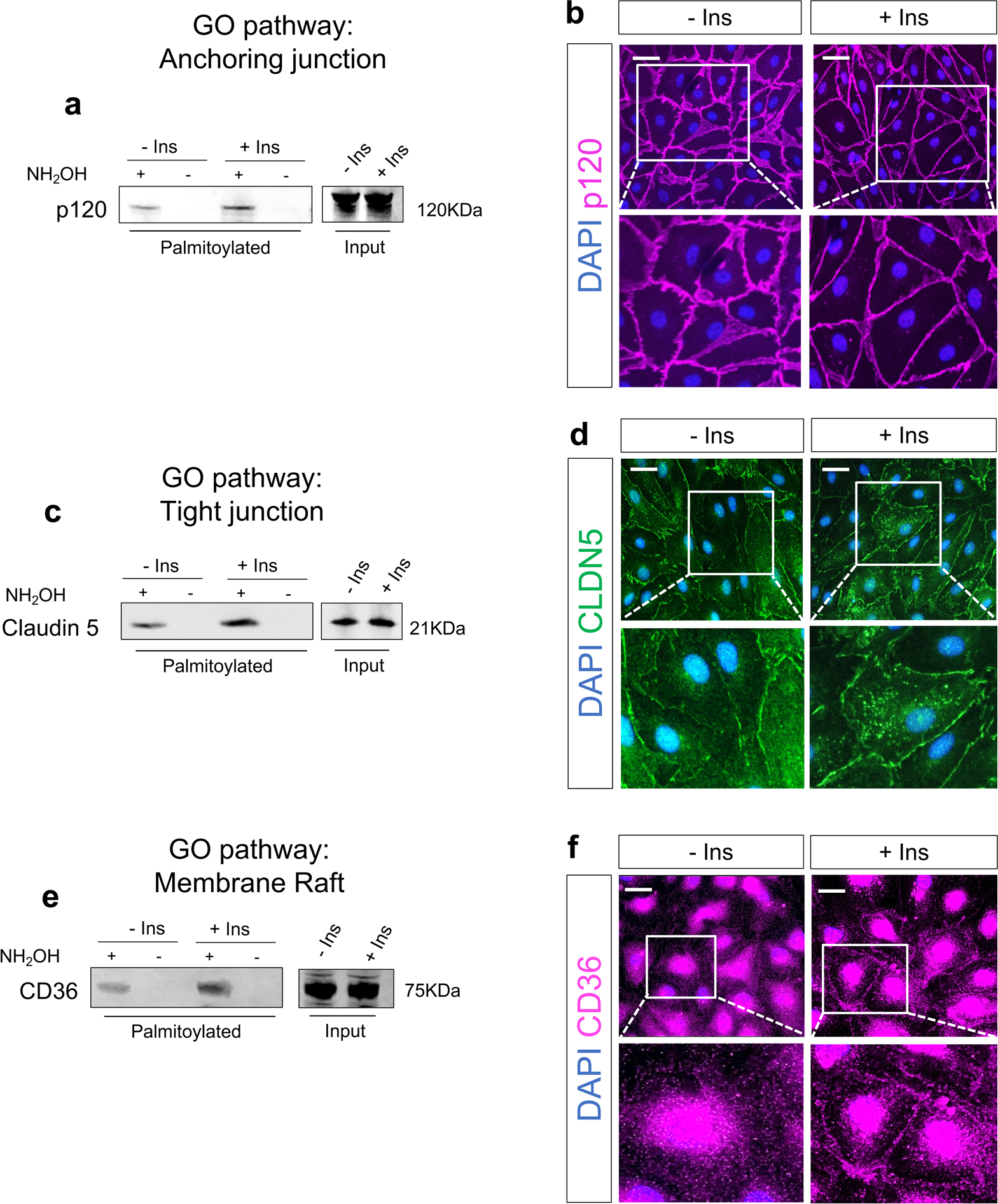
Insulin induces palmitoylation of anchoring junction p120-catenin, tight junction claudin 5 and CD36. **(a, c, e)** Western blot analysis of palmitoylated p120-catenin, claudin 5 and CD36 at baseline and following insulin treatment by acyl-RAC assays. (**b, d, f**) Representative images of immunofluorescence depicting cellular localization of p-120 catenin, claudin 5 and CD36. DAPI is a nuclear staining. Data are from three independent experiments.

### Role of Protein Palmitoylation in LECs

Since several of the above palmitoylated proteins are involved in LEC integrity, we next addressed the potential role of global palmitoylation in lymphatic VE-cadherin junctional remodeling. Insulin treatment induced straighter VE-cadherin– lined junctions and increased actin filament (phalloidin) anchoring to VE-cadherin perpendicularly with colocalization evident from the yellow merge between green VE-cadherin and red phalloidin (**Figure 3a**). Quantification shows that insulin treatment associates with significant i) reduction in junctional width (p<0.01), ii) increased percentage of thin VE-cadherin junctions (p<0.01) (**Figure 3b**) and iii) higher VE-cadherin-actin colocalization (**Figure 3c**) with a 1.5 increase in Mander’s overlap coefficient (**Figure 3d**). Inhibiting palmitoylation with 2-bromopalmitate(45) (2-BP, 20 μmol/L) blunts insulin induced linearization of VE-cadherin, increases junctional width and thickness (p<0.05) (**Figures 3b-c**), and reduces VE-cadherin-actin co-localization (Mander’s coefficient, p<0.05) (**Figure 3d**). Similarly, TEER measurements conducted in a fully confluent hLEC monolayer show a tighter monolayer in the insulin-treated condition as compared to media only (p<0.05 from time point 09:68h up to 15:00h) and blunting of the insulin effect with 2-BP (p<0.05 from time point 9:21h up to 18:00h) (**Figure 3e**). As expected, lipopolysaccharide (LPS) fully disrupts integrity of the LEC monolayer, causing a rapid decline in TEER (**Figure 3e-f**).

**Figure 3.**
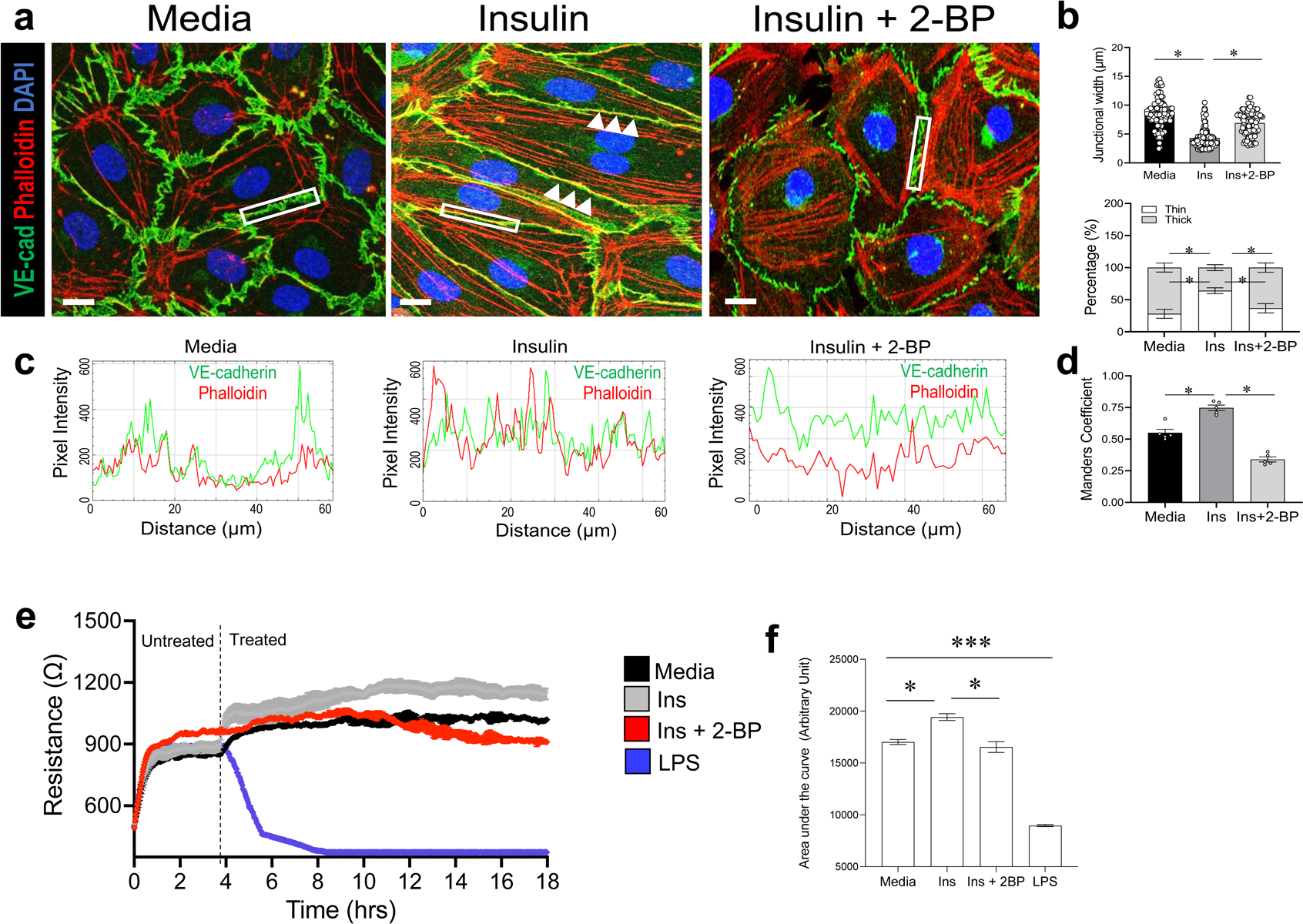
Insulin induces straighter VE-cadherin in LECs. (**a**) Confocal microscopy images showing that insulin (100nM) induces straighter (white arrows) VE-cadherin in hLECs as compared to media only. 2-bromo palmitate (2-BP, 20µM), concomitant to insulin treatment, reverses insulin-mediated VE-cadherin linearization. Scale bar: 20 μm. (**b**) Quantification VE-cadherin linear junctions. (**c**) Stacks of the images were analyzed in Fiji by JACoP plugin for colocalization analysis (white box). Data (n=10 cells in 10 images/group) are means ± SE. One-way ANOVA with Sidak’s post hoc test for multiple comparisons (*p<0.05). (**d**) VE-cadherin co-localization with actin (Mander’s coefficient. (**e**) TEER measurements in a fully confluent LEC monolayer treated with media only, insulin (100 nM) or insulin (100nM) and 2-BP (20μm), or LPS (50 ng/mL). (**f**) Area under the curve for each treatment. Data (n=6/group) are means ± SEM. * P<0.05, ***P<0.001 by 2-tailed Student *t* test.

### Role of CD36 in Protein Palmitoylation in LECs

Palmitoylation can be fueled by uptake of palmitate, imported across the plasma membrane via the long-chain FA transporter CD36 (46). Alternatively, palmitate can be synthesized endogenously from non-FA precursors by FA synthase (FASN) (47). To determine whether CD36 is a rate-limiting substrate for palmitoylation cycling, we silenced CD36 expression in hLECs using small interfering RNA (siCD36) before subjecting cells to insulin treatment (100 nM, 6h) (**Figure 4a**) and proteomic analysis. CD36 silencing, confirmed by western blotting, did not affect FASN expression (**Figure 4b**). Surprisingly, CD36 silencing associates with a 1.8-fold increase in palmitoylated targets at baseline (749 vs 380 targets in siCtrl LECs), and with an almost 3-fold increase in targets in insulin treated hLECs (574 vs 176 targets in WT LECs) (**Figure 4c-d, Supplemental Figure S3,** and **Supplemental Tables S3** and **S4**). As compared to siCtrl and insulin stimulation, CD36 silencing and insulin treatment increases palmitoylation of proteins involved in mitochondrial respiration (p<2.2e-16, FDR < 2.2e-16), protein localization to the endoplasmic reticulum (p<3.34E-12; FDR<7.11E-10), and neutrophil mediated immunity (p<2.2e-16, FDR < 2.2e-16) after insulin stimulation (**Figure 4e**, and **Supplemental Table S5**) while palmitoylation of 90 proteins, among these R-Ras and claudin-11 was lost (**Supplemental Table S6**). Insulin induced phosphorylation of AKT at S473 was blunted in siCD36 LECs as compared to control cells (**Figure 4f**) as reported in other cell types (48) suggesting that in the absence of CD36, insulin signaling might regulate palmitoylation via pathways other than PI3kinase/AKT.

**Figure 4.**
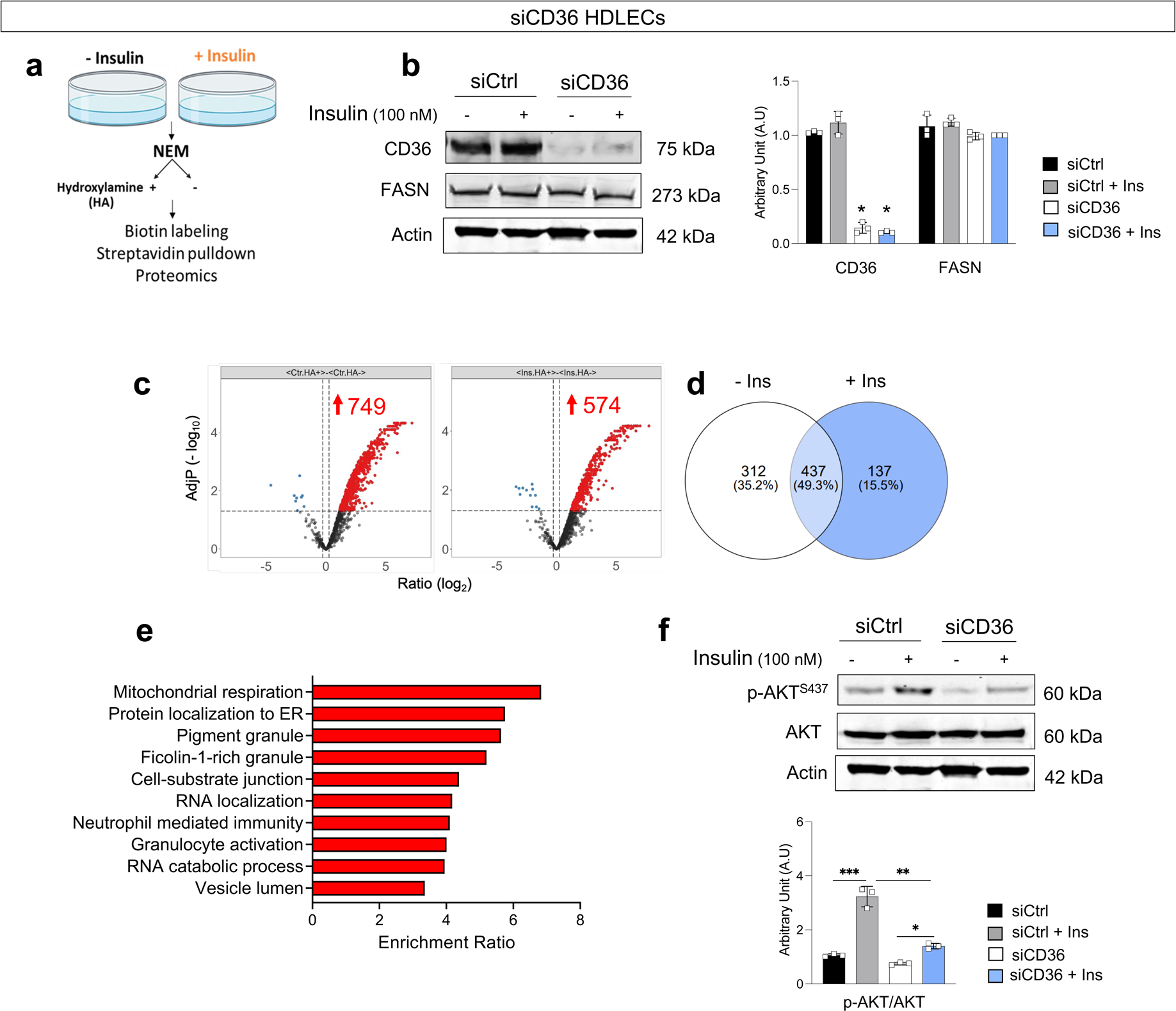
CD36 regulates LEC palmitoylation proteomics profile in hLECs. (**a**) Strategy for isolating palmitoylated proteins. (**b**) CD36 and FASN expression by western blotting and quantification following siRNA strategy. Actin is loading control. (**c**) Volcano plot and (**d**) Venn diagram of palmitoylated targets in media (Ctrl, left panel) or insulin (right panel) treated siCD36 LECs. Three matched groups of control and insulin treated cell preps were analyzed and normalized fold changes with P values were generated for individual proteins. (**e**) Pathway analysis of enriched palmitoylated proteins in siCD36 versus siCtrl LECs after insulin treatment. (**f**) Expression of AKT and pAKT^S473^ by western blotting, and quantification in siCtrl and siCD36 before and after 30 minutes insulin treatment (100 nM). Actin is loading control. Data (n=3/group) are means ± SEM. *P<0.05, **P<0.01, ***P<0.001 by 2-tailed Student *t* test.

### Insulin Induces Palmitoylation of Tetraspanin CD63 and its Interaction with Integrin β1 in LECs Devoid of CD36

CD36 silencing increased palmitoylation of tetraspanin CD63 at baseline and after insulin signaling (**Supplemental Tables S4 and S5**) which was validated by AcylRAc pulldown and western blotting (**Figure 5a-b**). CD63 is a member of the transmembrane-4 glycoprotein superfamily that clusters with other tetraspanin family members (i.e., CD53) and with integrins (49) (i.e., integrin β1) to regulate membrane protein trafficking, leukocyte recruitment, and adhesion processes (50). In addition, there is evidence that signaling pathways initiated by growth factors synergize functionally with those triggered by integrins (51) to regulate interaction between cell surface with the extracellular matrix. To determine whether insulin increases interaction between CD63 and integrin β1, we performed a proximity ligation (PLA) assay in siCD36 hLECs in the presence or absence of insulin (100 nM, 6h). We found that insulin significantly increases the PLA signal as compared to untreated siCD36 cells (**Figure 5c**) supporting increased interaction between CD63 and Integrin β1, despite no effect on expression of either protein (data not shown). These data show that in CD36 deficient LECs, insulin upregulates palmitoylation of CD63 enhancing engagement with integrin β1.

**Figure 5.**
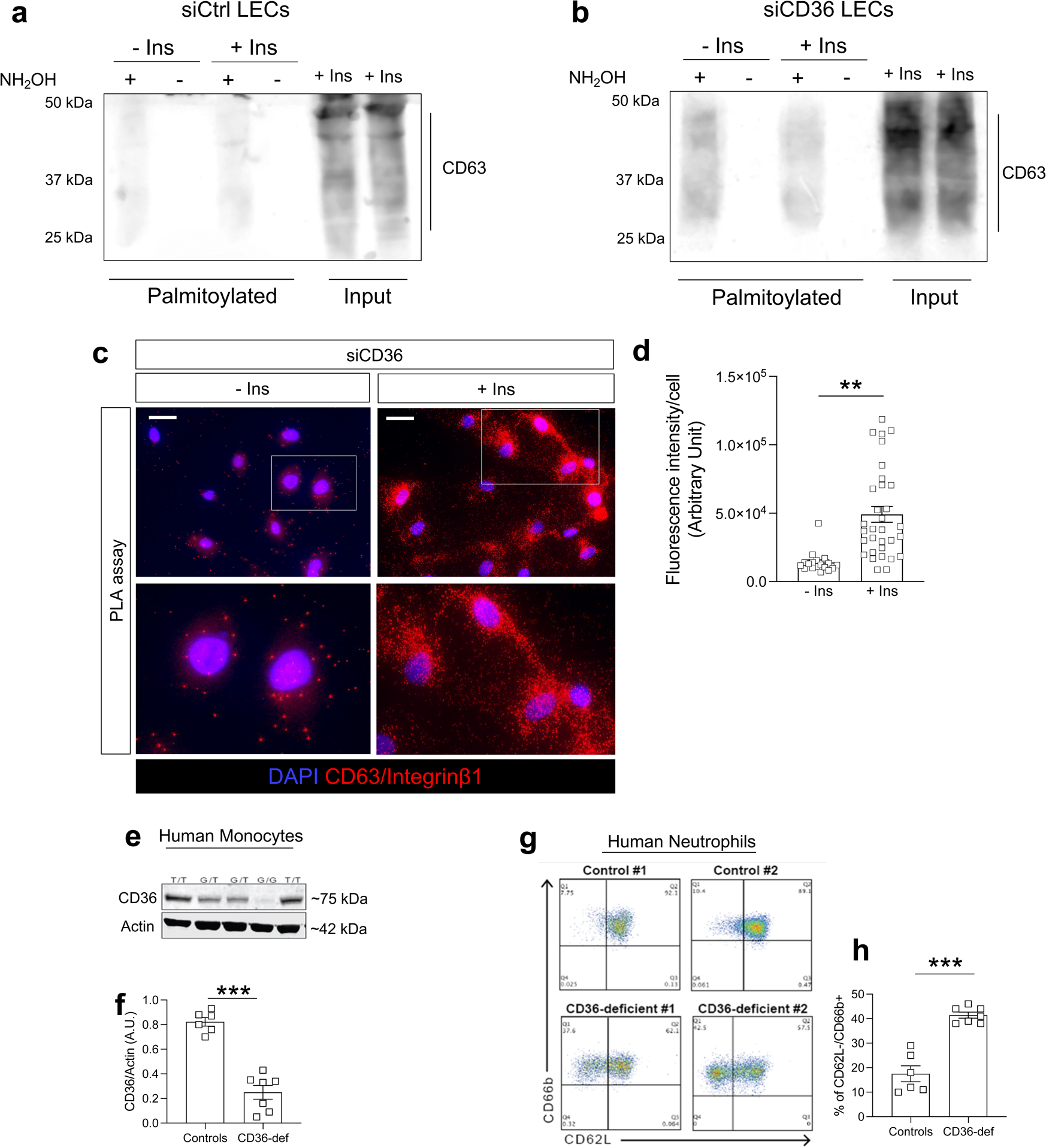
Loss of CD36 increases tetraspanin CD63 palmitoylation and interaction with integrin β1. (**a-b**) Western blot analysis of pulldown palmitoylated by acyl-RAC assays CD63 in siCtrl and siCD36 hLEC at baseline and following insulin treatment. Lysates are from three independent experiments that were pulled together. (**c**) Representative images of the proximity ligation assay (PLA) depicting interaction between CD63 and integrin β1 (red fluorescence) in siCD36 HDLECs in presence or absence of insulin (100nM, 6h) and (**d**) quantification. Signal (n=3/10 cells/treatment) was quantified as a mean intensity of fluorescence and background was subtracted as the mean intensity of fluorescence of five different empty areas for each field using ImageJ software. Data (n=3/group) are means ± SEM. (**e-f**) CD36 expression by western blotting and quantification in human monocytes isolated from pre-diabetic obese body mass index (BMI)-matched unrelated individuals, with or without a coding single nucleotide polymorphism (SNP) in the *CD36* gene, rs3211938 (T>G). (**g-h**) Human peripheral blood neutrophils were identified by flow cytometry by using a Side-scatter^hi^ CD66^+^ CD11b^+^CD15^+^CD14^-^ gate and assessed for cell membrane CD62L expression. Data (n=6-7/group) are means ± SEM. *P<0.05, **P<0.01, ***P<0.001 by 2-tailed Student *t* test.

### Higher Neutrophil Activation in Prediabetic CD36 Deficient Subjects

We previously reported that CD36 deletion specific to blood and lymphatic endothelial cells (*Tie2Cre;Cd36^-/-^*), but not in enterocytes (*VillinCre;Cd36^-/-^*), induces neutrophil activation and extravasation into several tissues (52). Neutrophil rolling and interaction with the inflamed endothelium occurs via CD62L recruitment of a CD63/integrin β1 complex (53), then CD62L is shed upon neutrophil activation to slow movement of the neutrophil along the endothelium (54). To explore potential relevance of our findings to individuals with CD36 deficiency, we recruited unrelated pre-diabetic obese body mass index (BMI)-matched individuals (**Table 1**), with or without a coding single nucleotide polymorphism (SNP) in the *CD36* gene, rs3211938 (T>G). This SNP, relatively common in populations of African ancestry, causes total CD36 protein deficiency in homozygous (G/G) (∼10% frequency) (55). We then examined whether neutrophils isolated from CD36 deficient individuals might be more activated by analyzing levels of CD62L, a selectin molecule that promotes vascular adhesion, that is shed from activated leukocytes to facilitate transmigration into inflamed tissues. Peripheral blood polymorphonuclear cells were isolated by density centrifugation and analyzed for neutrophil CD62L levels by flow cytometric analysis (**Figure 5f-g**). Neutrophils from CD36-deficient as compared to non-carriers with normal CD36 had lower levels of CD62L indicative of ongoing neutrophil activation.

### Comparison of Insulin Regulated Palmitoylation Profile between LECs and BECs

We examined how the human LEC palmitoylation profile compares with that of human BECs by interrogating an available BEC palmitoylation data set (22). Although a similar number of palmitoylated targets are detected in LECs and BECs at baseline, a higher percentage of palmitoylated protein is regulated by insulin in LECs (40% of all palmitoylated proteins) than BECs (10% of all palmitoylated proteins) (**Figure 6a**). Palmitoylation is catalyzed by ∼20 DHHCs acyltransferases (16) while depalmitoylation is performed by a family of depalmitoylation enzymes that includes APT-1 and -2(18) and ABHD17-A, -B and -C (19). Data mining of available RNA sequencing data in BECs and LECs (39) show that the vascular and lymphatic beds are equipped with specific sets of palmitoylation enzymes (**Figure 6b**) suggesting different palmitoylation targets in line with their specific physiological roles. Expression of genes encoding for *ZDHHC 1, 2, 9, 13, 16, 18* is significantly higher in BECs, while expression of genes encoding for *ZDHHC 3, 12, 14*, and *23* is significantly higher in LECs. No changes are found in expression of genes encoding for *ZDHHC 4, 5, 6, 7, 8, 20, 21,* and *24*. Gene expression of depalmitoylation enzymes *APT-1, APT-2, ABHD17 A*, and *ABHD17C* are similar for LECs and BECs except for *ABHD17B*, which is significantly higher in LECs (**Figure 6b**). Insulin receptor (*INSR*) and *CD36* expression levels are significantly increased in LECs as compared to BECs (p<0.005 and p<0.0001, respectively) (**Figure 6c-d**) while *FASN* expression (p=0.0033) is higher in BECs (**Figure 6e**).

**Figure 6:**
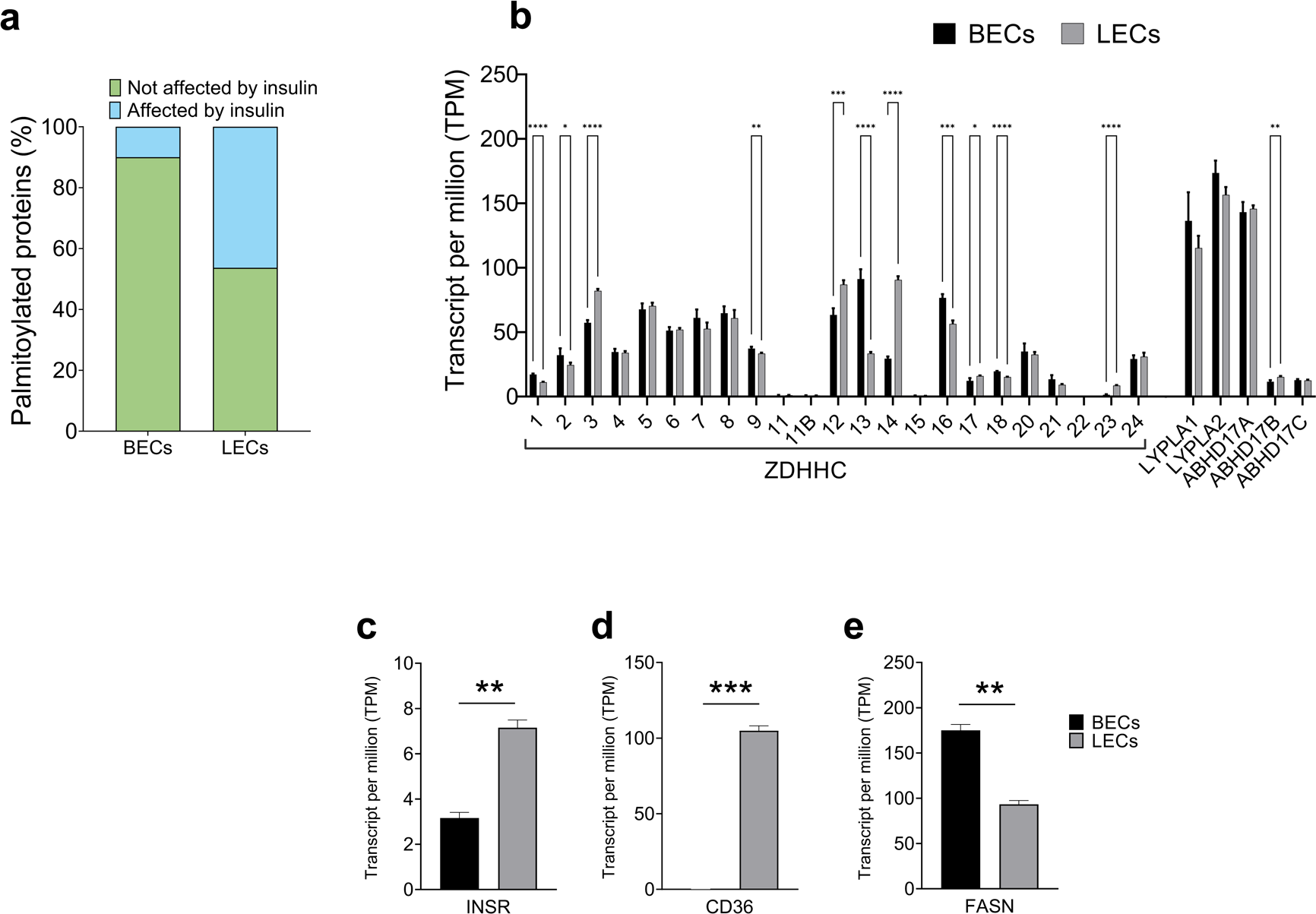
BECs and LECs have distinct i-regulated palmitoylation profile. (**a**) Number of palmitoylated targets in BECs and LECs at baseline and after insulin treatment (both at 100 nM for 6h) was obtained by comparing our data set to a previous palmitoylation proteomics study(22). (**b-e**) Expression levels of *ZDHHc*, depalmitoylation enzymes, *INSR, CD36* and *FASN* obtained from RNA sequencing in BECs and LECs(39). *P<0.05, **P<0.01, ***P<0.001 by 2-tailed Student *t* test.

## DISCUSSION

The metabolic syndrome, characterized by an abnormal lipid profile and insulin resistance, is associated with impaired lymphatic function in peripheral tissues (27, 41, 56). An abnormal lipid profile is detrimental for LEC survival (57, 58) and monolayer integrity, in line with reports in rodent models of hypercholesterolemia (8, 9), dyslipidemia (59, 60) and diet-induced obesity (26, 61) supporting the existence of a link between impaired lipid metabolism and loss of lymphatic vessel integrity.

LECs have higher rates of FA uptake and FAO compared to BECs, along with increased expression of the FA transporter CD36 and of proteins involved in FA import into mitochondria; FAO is essential for LEC differentiation by supplying acetyl-CoA as a substrate for acetylation of histones at lymphangiogenic genes to enhance transcription (2, 62). Acetyl-CoA can also be converted to palmitoyl-CoA to support palmitoylation, a reversible post-translational modification that involves linkage of a FA chain via a thioester bond predominantly to a cysteine (16). Reversible palmitoylation modulates spatiotemporal control of protein function and activity as well as protein–protein and membrane–protein interactions (14). Our study demonstrates highly active protein palmitoylation in LECs regulated by insulin and by CD36. CD36 silencing and the associated insulin resistance dramatically increase overall protein palmitoylation. Dysregulated palmitoylation is reported in a plethora of pathologies including neurological diseases (63), cancer (64), metabolic syndrome (20–22, 65), bacterial and viral infections(66), and histamine-mediated endothelial inflammation (17), and is indicative of disrupted lipid homeostasis.

Since insulin regulates lipid metabolism (67) and protein lipidation (22), we investigated whether insulin regulates LEC integrity via palmitoylation. Insulin remodeled lymphatic junctions by inducing linearization of VE-cadherin, an effect that was reversed by inhibition of palmitoylation with 2-bromopalmitate. VE-cadherin junctional morphological changes depend on junctional anchoring (42) and our proteomics and validation studies showed insulin-mediated increased palmitoylation of anchoring junction proteins catenin δ1, or p120-catenin, among additional members of the armadillo-related protein family (68) (i.e., plakophilin 3) known to facilitate protein-protein interaction (69). P120-catenin is enriched at adherens junctions and stabilizes cadherins via the cytoskeleton (70) as confirmed by our immunofluorescence analysis showing straighter p120-catenin at the plasma membrane following insulin treatment. Insulin also increased the palmitoylation of tight junction component claudin 5, which has overlapping distribution with VE-cadherin as previously reported (70, 71). Tight junctions are abundant in cerebrovascular endothelial cells and are necessary for blood-brain barrier integrity and maintenance (72). *Claudin-5^+/–^* mice exposed to ultraviolet B irradiation develop edema due to lymphatic vessel leakage (73) suggesting requirement of claudin-5 for the maintenance of collecting lymphatic vessel integrity in additional anatomical regions besides the brain. Interestingly, claudin 5 is negatively regulated by the transcriptional repressor forkhead box protein O1, FOXO1, whose activity increases during impaired insulin/AKT signaling (74). It is possible that FOXO1 activation during insulin resistance affects claudin-5 stability and turnover via altered palmitoylation.

Insulin induced robust palmitoylation of CD36, a transmembrane receptor with established functions in long-chain FA (i.e., palmitate) uptake and lipid metabolism (75). CD36 localization at the plasma membrane is finely regulated by metabolic status which dictates receptor turnover, degradation and cellular functions (76). Using immunofluorescence, we show that insulin treatment promoted CD36 plasma membrane localization in hLECs in line with a previous report showing CD36 translocation to the muscle cell plasma membrane following insulin stimulation (77). Since CD36 is a major route for the uptake of circulating palmitate into cells, we sought to determine whether its silencing in hLECs would affect global palmitoylation. We found that CD36 silencing in hLECs doubled rates of lymphatic palmitoylation, targeting proteins involved in inflammation, leukocyte recruitment, and neutrophil degranulation. Of note, palmitoylation of tetraspanin CD63 was significantly increased in siCD36 cells at baseline and following insulin treatment. Tetraspanins and many of their partner proteins, including CD36 (78), have palmitoylation sites to support interactions with other tetraspanins and integrins for the propagation of distinct signaling pathways (49, 79). We found that in siCD36 hLECs, insulin increased CD63 interaction with binding partner integrin β1, which previous studies have shown recruits and activates neutrophils via CD62L (53). Our human data show that circulating neutrophils isolated from individuals carrying the rs3211938 SNP, that decreases CD36 expression, have higher CD62L shedding than non-carriers, reminiscent of neutrophil activation. Since blood neutrophils have negligible CD36 expression (80), it is possible that more CD62L shedding and higher neutrophil activation in rs3211938 carriers might reflect neutrophil interaction with the endothelium devoid of CD36 which is in line with our previous studies showing neutrophil activation and infiltration in mice lacking CD36 globally or in the endothelium but not in enterocyte (52). Genetic variants in the CD36 gene that reduce protein expression are relatively common (81–83). In contrast to SNPs that might impact CD36 in a tissue-specific fashion (84), the coding SNP studied in our cohort impacts CD36 levels in all tissues. Thus, we cannot exclude that other cell types i.e., monocytes/macrophages with high expression of CD36 and CD63 (85) and undergo palmitoylation (86) might be involved in neutrophil activation in our cohort. However, the SNP rs3211938 SNP, which is exclusive to African Americans (minor allele incidence is ∼9%) is associated with increased incidence of endothelial dysfunction and vascular stiffening (87), stroke (88), and cardiovascular diseases (89). The contribution of rs3211938 to lymphatic endothelial dysfunction, however, is not known and warrants further investigation.

Our study found that CD36 silencing increased palmitoylation of proteins involved in glycolysis and gluconeogenesis, which may have contributed to the increased glycolytic levels and expression of genes encoding for glycolytic enzymes previously reported by our group in cultured hLECs and in isolated LECs from gut lymphatics (3). We previously showed that CD36 silencing in LECs associates with reduced VEGF-C mediated phosphorylation of VEGFR-2/AKT and impairs LEC migration, tube formation, and monolayer integrity. *In vivo*, mice with inducible deletion of CD36 in LECs manifested compromised integrity of the collectors, spillover of lymph in the mesentery and spontaneous obesity (3). Our current study adds a new mechanism for CD36 effects in LECs as CD36 silencing causes a loss of palmitoylation by members of the RAS family, important for lymphangiogenesis (90), and by claudin 11, highly enriched at valves in lymphatic collectors (91). How palmitoylation contributes to RAS or claudin 11 function in lymphatics is not known and warrants further investigation. However, our results highlight that CD36 might regulate lymphatic function via multiple mechanisms.

Insulin primarily regulated palmitoylation of proteins involved in angiogenesis and cell migration in BECs (22), while small GTPases, ubiquitination and cell-cell junction proteins were primarily affected in LECs. These results might be due to differences between BECs and LECs related to i) insulin sensitivity, ii) expression profile of zDHHCs and depalmitoylation enzymes, and iii) FAO flux (1). However, since regulation of palmitoylation is not fully understood, additional molecular and metabolic cues may underlie these differences.

In conclusion, our study determined that protein palmitoylation plays an important role in LEC function. The information related to the LEC palmitoylation profile, distinct from that of BECs, especially in relation to the regulation by insulin, provides a framework for future studies that will examine the role of lipid metabolism and insulin signaling in lymphatic system homeostasis and dysfunction. The unexpected effect of CD36 silencing in increasing the palmitoylation profile may help explain the endothelial dysfunction observed in a preclinical model of CD36 deficiency. We further speculate that the abnormal neutrophil activation observed in prediabetic CD36-deficient individuals is consequent to immune cell-endothelial interaction (92) mediated in part by changes in palmitoylation.

## DATA AVAILABILITY STATEMENT

The authors confirm that the data supporting the palmitoylation proteomics findings of this study are available within the article and its supplementary materials. The data that support additional findings of this study are available on request from the corresponding author.

## ACKNOLWDGEMENTS

We thank Petra Erdmann-Gilmore, Dr. Yiling Mi, Alan Davis and Rose Connors for their assistance with proteomics experiments. The cartoon in a figure was created using BioRender.

## AUTHOR INFORMATION

### Contributions

LLG and NA recruited human subjects. SGN, XW, JM, QZ, JZ, RM, DF, LLV, AEG, CFS, NA and VC conducted or supervised the studies. SGF, QZ, JZ and VC analyzed the data and performed the statistical analyses. VC designed the studies and wrote the manuscript. VC is the guarantor of this work and, as such, had full access to all the data in the study and takes responsibility for the integrity of the data and the accuracy of the data analysis. All authors critically reviewed and edited the manuscript.

## FUNDING SOURCES

This study was supported by the startup fund from Saint Louis University, SLU Institute for Drug & Biotherapeutic Innovation (IDBI) seed grant and from the SLU Research Institute Growth Fund, by NIH grants R56HL165199 (VC), R01HL157154 (CFS), and P30DK020579 (CFS). The proteomic experiments were performed at the Washington University Proteomics Shared Resource (WU-PSR), supported in part by the WU Institute of Clinical and Translational Sciences (NCATS UL1 TR000448), the Mass Spectrometry Research Resource (NIGMS P41 GM103422) and the Siteman Comprehensive Cancer Center Support Grant (NCI P30 CA091842).

## Notes

### Competing Interest Statement

The authors have declared no competing interest.

