## Supplemental Tables for "Insulin regulates lymphatic endothelial function via palmitoylation"

**Supplemental Table S1.** List of primary and secondary antibodies.

| Immunofluorescence | | | | | |
|---|---|---|---|---|---|
| **Antibody** | **Species** | **Identifier** | **Cat. No.** | **Clone** | **Dilution** |
| CD36 | Goat anti-human | R&D Systems | AF1955 | n/a | 1:100 |
| Claudin 5 | Rabbit anti-human | Abcam | Ab15106 | n/a | 1:250 |
| p120 | Mouse anti-human | Invitrogen | 33-9700 | 6H11 | 1:200 |
| VE-cadherin | Mouse anti-human | SCBT | Sc-9989 | F-8 | 1:100 |
| CD63 | Rabbit anti-human | Invitrogen | PA5-92370 | n/a | 1:100 |
| integrin β1 | Mouse anti-human | Invitrogen | 14-0299-82 | Ts2/16 | 1:100 |
| Secondary | Donkey anti-Rabbi AF488 | Invitrogen | A21206 | n/a | 1:250 |
| Secondary | Donkey anti-Mouse AF594 | Invitrogen | A11005 | n/a | 1:250 |
| Secondary | Donkey anti-Mouse AF568 | Invitrogen | A10037 | n/a | 1:250 |
| Secondary | Donkey anti-Goat AF647 | Invitrogen | A32849 | n/a | 1:250 |
| **Western Blotting** | | | | | |
| AKT | Rabbit anti-human | Cell Signaling | 4691 | C67E7 | 1:500 |
| p-AKT$^{ser473}$ | Rabbit anti-human | Cell Signaling | 4060 | D9E | 1:500 |
| CD36 | Goat anti-human | R&D Systems | AF1955 | n/a | 1:1000 |
| Claudin 5 | Mouse anti-human | Invitrogen | 35-2500 | 4C3C2 | 1:1000 |
| FASN | Rabbit anti-human | Abcam | ab128856 | EPR7465 | 1:1000 |
| p120 | Rabbit anti-human | Invitrogen | PA5-82545 | n/a | 1:500 |
| CD63 | Rabbit anti-human | Invitrogen | PA5-92370 | n/a | 1:500 |
| integrin β1 | Rabbit anti-human | Abcam | ab52971 | EP1041Y | 1:2000 |
| β-actin | Mouse anti-human | Cell Signaling | 3700 | 8H10D10 | 1:10,000 |
| Secondary | Donkey anti-mouse 800CW | LiCor | 926-32212 | n/a | 1:10,000 |
| Secondary | Donkey anti-goat 680RD | LiCor | 926-68074 | n/a | 1:10,000 |
| Secondary | Donkey anti-rabbit 680RD | LiCor | 926-68073 | n/a | 1:10,000 |
| Secondary | Donkey anti-rabbit 800CW | LiCor | 926-32213 | n/a | 1:10,000 |

### Supplemental Table S2. Palmitoylated proteins in human LECs.

| ENSEMBL ID | Gene Symbol | Description | UNIPROT | *p*-value | -Log Ratio (+/-HA) |
|---|---|---|---|---|---|
| ENSG00000120437 | ACAT2 | acetyl-CoA acetyltransferase 2 | Q9BWD1 | 9.70E-07 | 6.42 |
| ENSG00000168899 | VAMP5 | vesicle associated membrane protein 5 | O95183 | 5.50E-07 | 6.42 |
| ENSG00000049245 | VAMP3 | vesicle associated membrane protein 3 | Q15836 | 4.40E-07 | 6.39 |
| ENSG00000169908 | TM4SF1 | transmembrane 4 L six family member 1 | P30408 | 1.70E-06 | 6.29 |
| ENSG00000110651 | CD81 | CD81 molecule | E9PJK1 | 4.90E-06 | 6.21 |
| ENSG00000249751 | ECSCR | endothelial cell surface expressed chemotaxis and apoptosis regulator | Q19T08 | 1.10E-06 | 6.01 |
| ENSG00000142089 | IFITM3 | interferon induced transmembrane protein 3 | Q01628 | 5.00E-07 | 6.01 |
| ENSG00000002586 | CD99 | CD99 molecule (Xg blood group) | P14209 | 7.70E-07 | 5.67 |
| ENSG00000116754 | SRSF11 | serine and arginine rich splicing factor 11 | Q05519 | 3.10E-06 | 5.62 |
| ENSG00000129625 | REEP5 | receptor accessory protein 5 | Q00765 | 2.40E-06 | 5.56 |
| ENSG00000133818 | RRAS2 | RAS related 2 | P62070 | 9.10E-07 | 5.56 |
| ENSG00000134247 | PTGFRN | prostaglandin F2 receptor inhibitor | Q9P2B2 | 1.60E-06 | 5.55 |
| ENSG00000135404 | CD63 | CD63 molecule | A0A024RB05 | 3.10E-06 | 5.5 |
| ENSG00000148175 | STOM | stomatin | F8VSL7 | 1.20E-06 | 5.42 |
| ENSG00000141526 | SLC16A3 | solute carrier family 16 member 3 | A0A024R8U1 | 3.80E-06 | 5.41 |
| ENSG00000102007 | PLP2 | proteolipid protein 2 | A0A024QYW3 | 6.60E-05 | 5.4 |
| ENSG00000059573 | ALDH18A1 | aldehyde dehydrogenase 18 family member A1 | P54886 | 6.10E-06 | 5.25 |
| ENSG00000010278 | CD9 | CD9 molecule | A6NNI4 | 5.60E-06 | 5.22 |
| ENSG00000075239 | ACAT1 | acetyl-CoA acetyltransferase 1 | A0A140VJX1 | 1.90E-06 | 5.21 |
| ENSG00000076706 | MCAM | melanoma cell adhesion molecule | A0A024R3I5 | 2.10E-06 | 5.19 |
| ENSG00000078140 | UBE2K | ubiquitin conjugating enzyme E2 K | P61086 | 1.50E-06 | 5.17 |
| ENSG00000130725 | UBE2M | ubiquitin conjugating enzyme E2 M | A0A024R4T4 | 2.30E-06 | 5.04 |
| ENSG00000108848 | LUC7L3 | LUC7 like 3 pre-mRNA splicing factor | J3KPP4 | 3.30E-06 | 5.03 |
| ENSG00000115310 | RTN4 | reticulon 4 | Q9NQC3 | 2.40E-06 | 5.02 |
| ENSG00000177889 | UBE2N | ubiquitin conjugating enzyme E2 N | P61088 | 9.50E-06 | 4.87 |
| ENSG00000139921 | TMX1 | thioredoxin related transmembrane protein 1 | Q9H3N1 | 3.10E-06 | 4.7 |
| ENSG00000136156 | ITM2B | integral membrane protein 2B | Q9Y287 | 1.10E-05 | 4.64 |
| ENSG00000143878 | RHOB | ras homolog family member B | P62745 | 1.90E-05 | 4.56 |
| ENSG00000143222 | UFC1 | ubiquitin-fold modifier conjugating enzyme 1 | Q9Y3C8 | 1.30E-05 | 4.56 |
| ENSG00000188313 | PLSCR1 | phospholipid scramblase 1 | O15162 | 1.00E-05 | 4.54 |
| ENSG00000103018 | CYB5B | cytochrome b5 type B | J3KNF8 | 4.40E-06 | 4.49 |
| ENSG00000026508 | CD44 | CD44 molecule (Indian blood group) | P16070 | 3.70E-06 | 4.42 |
| ENSG00000243279 | PRAF2 | PRA1 domain family member 2 | A0A024QZ22 | 1.90E-05 | 4.42 |
| ENSG00000131037 | EPS8L1 | EPS8 like 1 | Q8TE68 | 0.0038 | 4.41 |

| | | | | | |
|---|---|---|---|---|---|
| ENSG00000197870 | PRB3 | proline rich protein BstNI subfamily 3 | A0A0G2JNB4 | 3.00E-04 | 4.41 |
| ENSG00000072274 | TFRC | transferrin receptor | P02786 | 1.70E-05 | 4.34 |
| ENSG00000112739 | PRPF4B | pre-mRNA processing factor 4B | A0A024QZY5 | 1.90E-05 | 4.26 |
| ENSG00000130985 | UBA1 | ubiquitin like modifier activating enzyme 1 | A0A024R1A3 | 5.70E-06 | 4.26 |
| ENSG00000185651 | UBE2L3 | ubiquitin conjugating enzyme E2 L3 | P68036 | 1.10E-05 | 4.25 |
| ENSG00000136026 | CKAP4 | cytoskeleton associated protein 4 | A0A024RBH2 | 1.90E-05 | 4.24 |
| ENSG00000198755 | RPL10A | ribosomal protein L10a | P62906 | 5.90E-05 | 4.24 |
| ENSG00000110090 | CPT1A | carnitine palmitoyltransferase 1A | P50416 | 1.70E-05 | 4.23 |
| ENSG00000072401 | UBE2D1 | ubiquitin conjugating enzyme E2 D1 | A0A087WW00 | 1.10E-04 | 4.19 |
| ENSG00000114353 | GNAI2 | G protein subunit alpha i2 | B3KP24 | 9.40E-06 | 4.14 |
| ENSG00000129353 | SLC44A2 | solute carrier family 44 member 2 | A0A088QCU6 | 1.00E-05 | 4.13 |
| ENSG00000006451 | RALA | RAS like proto-oncogene A | P11233 | 1.00E-05 | 4.12 |
| ENSG00000075142 | SRI | sorcin | P30626 | 5.30E-04 | 4.11 |
| ENSG00000135821 | GLUL | glutamate-ammonia ligase | A8YXX4 | 0.027 | 4.04 |
| ENSG00000198561 | CTNND1 | catenin delta 1 | O60716 | 4.70E-05 | 3.97 |
| ENSG00000182899 | RPL35A | ribosomal protein L35a | P18077 | 1.50E-04 | 3.83 |
| ENSG00000155380 | SLC16A1 | solute carrier family 16 member 1 | A0A024R0H1 | 1.50E-04 | 3.82 |
| ENSG00000059758 | CDK17 | cyclin dependent kinase 17 | Q00537 | 8.90E-05 | 3.79 |
| ENSG00000160221 | GATD3B | glutamine amidotransferase class 1 domain containing 3 | A0A0B4J2D5 | 1.50E-04 | 3.76 |
| ENSG00000067057 | PFKP | phosphofructokinase, platelet | Q01813 | 2.50E-05 | 3.7 |
| ENSG00000143545 | RAB13 | RAB13, member RAS oncogene family | A0A087WWB9 | 5.80E-04 | 3.67 |
| ENSG00000106263 | EIF3B | eukaryotic translation initiation factor 3 subunit B | A0A024R821 | 6.00E-05 | 3.66 |
| ENSG00000144746 | ARL6IP5 | ADP ribosylation factor like GTPase 6 interacting protein 5 | A0A024R371 | 4.20E-04 | 3.64 |
| ENSG00000131016 | AKAP12 | A-kinase anchoring protein 12 | Q02952 | 3.20E-05 | 3.62 |
| ENSG00000262814 | MRPL12 | mitochondrial ribosomal protein L12 | P52815 | 6.20E-05 | 3.62 |
| ENSG00000100201 | DDX17 | DEAD-box helicase 17 | A0A1W2PQ51 | 3.10E-04 | 3.61 |
| ENSG00000168703 | WFDC12 | WAP four-disulfide core domain 12 | Q8WWY7 | 0.0073 | 3.61 |
| ENSG00000167996 | FTH1 | ferritin heavy chain 1 | A0A024R525 | 2.40E-05 | 3.6 |
| ENSG00000198431 | TXNRD1 | thioredoxin reductase 1 | Q16881 | 1.40E-04 | 3.54 |
| ENSG00000265808 | SEC22B | SEC22 homolog B, vesicle trafficking protein | O75396 | 1.30E-04 | 3.52 |
| ENSG00000166479 | TMX3 | thioredoxin related transmembrane protein 3 | Q96JJ7 | 3.00E-05 | 3.51 |
| ENSG00000154473 | BUB3 | BUB3 mitotic checkpoint protein | A0A140VJF3 | 4.60E-05 | 3.49 |
| ENSG00000141959 | PFKL | phosphofructokinase, liver type | P17858 | 9.20E-04 | 3.49 |
| ENSG00000164163 | ABCE1 | ATP binding cassette subfamily E member 1 | P61221 | 1.00E-04 | 3.45 |
| ENSG00000177674 | AGTRAP | angiotensin II receptor associated protein | Q6RW13 | 7.00E-04 | 3.44 |
| ENSG00000170889 | RPS9 | ribosomal protein S9 | A0A024R4M0 | 9.20E-05 | 3.41 |
| ENSG00000106636 | YKT6 | YKT6 v-SNARE homolog | A4D2J0 | 8.70E-05 | 3.4 |
| ENSG00000142541 | RPL13A | ribosomal protein L13a | P40429 | 2.40E-04 | 3.37 |

| | | | | | |
|---|---|---|---|---|---|
| ENSG00000181163 | NPM1 | nucleophosmin 1 | A0A140VJQ2 | 5.70E-04 | 3.33 |
| ENSG00000108528 | SLC25A11 | solute carrier family 25 member 11 | Q6IBH0 | 5.90E-04 | 3.33 |
| ENSG00000127824 | TUBA4A | tubulin alpha 4a | P68366 | 1.20E-04 | 3.33 |
| ENSG00000172725 | CORO1B | coronin 1B | A0A024R5K1 | 3.80E-05 | 3.32 |
| ENSG00000159840 | ZYX | zyxin | Q15942 | 1.20E-04 | 3.31 |
| ENSG00000178597 | PSAPL1 | prosaposin like 1 | Q6NUJ1 | 0.0097 | 3.3 |
| ENSG00000112081 | SRSF3 | serine and arginine rich splicing factor 3 | B2R6F3 | 5.80E-04 | 3.29 |
| ENSG00000102886 | GDPD3 | glycerophosphodiester phosphodiesterase domain containing 3 | Q7L5L3 | 0.0019 | 3.25 |
| ENSG00000185627 | PSMD13 | proteasome 26S subunit, non-ATPase 13 | Q9UNM6 | 1.00E-04 | 3.16 |
| ENSG00000022267 | FHL1 | four and a half LIM domains 1 | Q13642 | 2.10E-04 | 3.1 |
| ENSG00000111737 | RAB35 | RAB35, member RAS oncogene family | Q15286 | 1.40E-04 | 3.08 |
| ENSG00000089248 | ERP29 | endoplasmic reticulum protein 29 | P30040 | 1.90E-04 | 3.07 |
| ENSG00000111667 | USP5 | ubiquitin specific peptidase 5 | A0A140VJZ1 | 8.00E-04 | 3.07 |
| ENSG00000007168 | PAFAH1B1 | platelet activating factor acetylhydrolase 1b regulatory subunit 1 | P43034 | 2.50E-04 | 3.06 |
| ENSG00000091136 | LAMB1 | laminin subunit beta 1 | P07942 | 1.70E-04 | 3.05 |
| ENSG00000213625 | LEPROT | leptin receptor overlapping transcript | A0A087X0N2 | 1.80E-04 | 3.04 |
| ENSG00000109919 | MTCH2 | mitochondrial carrier 2 | Q9Y6C9 | 1.60E-04 | 3.04 |
| ENSG00000130309 | COLGALT1 | collagen beta(1-O)galactosyltransferase 1 | Q8NBJ5 | 0.0013 | 3 |
| ENSG00000115524 | SF3B1 | splicing factor 3b subunit 1 | B4DGZ4 | 4.10E-04 | 2.98 |
| ENSG00000103381 | CPPED1 | calcineurin like phosphoesterase domain containing 1 | Q9BRF8 | 3.90E-04 | 2.97 |
| ENSG00000103187 | COTL1 | coactosin like F-actin binding protein 1 | Q14019 | 2.50E-04 | 2.96 |
| ENSG00000186566 | GPATCH8 | G-patch domain containing 8 | Q9UKJ3 | 2.80E-04 | 2.95 |
| ENSG00000171564 | FGB | fibrinogen beta chain | P02675 | 0.028 | 2.93 |
| ENSG00000133318 | RTN3 | reticulon 3 | O95197 | 4.10E-04 | 2.92 |
| ENSG00000063177 | RPL18 | ribosomal protein L18 | A0A024QZD1 | 7.10E-05 | 2.91 |
| ENSG00000117448 | AKR1A1 | aldo-keto reductase family 1 member A1 | P14550 | 1.20E-04 | 2.9 |
| ENSG00000169710 | FASN | fatty acid synthase | P49327 | 1.30E-04 | 2.9 |
| ENSG00000138760 | SCARB2 | scavenger receptor class B member 2 | Q14108 | 8.90E-05 | 2.9 |
| ENSG00000105612 | DNASE2 | deoxyribonuclease 2, lysosomal | A0A024R7F4 | 6.20E-04 | 2.89 |
| ENSG00000146963 | LUC7L2 | LUC7 like 2, pre-mRNA splicing factor | Q9Y383 | 1.40E-04 | 2.88 |
| ENSG00000138119 | MYOF | myoferlin | Q9NZM1 | 1.60E-04 | 2.88 |
| ENSG00000125977 | EIF2S2 | eukaryotic translation initiation factor 2 subunit beta | P20042 | 2.20E-04 | 2.87 |
| ENSG00000115380 | EFEMP1 | EGF containing fibulin extracellular matrix protein 1 | A0A0S2Z4F1 | 1.30E-04 | 2.83 |
| ENSG00000221914 | PPP2R2A | protein phosphatase 2 regulatory subunit Balpha | P63151 | 1.10E-04 | 2.83 |
| ENSG00000065978 | YBX1 | Y-box binding protein 1 | P67809 | 2.30E-04 | 2.83 |
| ENSG00000241553 | ARPC4 | actin related protein 2/3 complex subunit 4 | P59998 | 7.00E-04 | 2.81 |

| | | | | | |
|---|---|---|---|---|---|
| ENSG00000244687 | UBE2V1 | ubiquitin conjugating enzyme E2 V1 | Q13404 | 0.0013 | 2.8 |
| ENSG00000105953 | OGDH | oxoglutarate dehydrogenase | A0A140VJQ5 | 1.00E-04 | 2.79 |
| ENSG00000130255 | RPL36 | ribosomal protein L36 | Q9Y3U8 | 4.40E-04 | 2.77 |
| ENSG00000131469 | RPL27 | ribosomal protein L27 | A0A024R1V4 | 9.40E-04 | 2.74 |
| ENSG00000124570 | SERPINB6 | serpin family B member 6 | A0A024QZX5 | 0.001 | 2.74 |
| ENSG00000137486 | ARRB1 | arrestin beta 1 | B7Z1Q3 | 3.90E-04 | 2.73 |
| ENSG00000149218 | ENDOD1 | endonuclease domain containing 1 | O94919 | 5.00E-04 | 2.72 |
| ENSG00000198898 | CAPZA2 | capping actin protein of muscle Z-line subunit alpha 2 | A4D0V4 | 5.60E-04 | 2.7 |
| ENSG00000233276 | GPX1 | glutathione peroxidase 1 | P07203 | 1.60E-04 | 2.7 |
| ENSG00000138814 | PPP3CA | protein phosphatase 3 catalytic subunit alpha | A0A0S2Z4C6 | 0.032 | 2.7 |
| ENSG00000278637 | H4C1 | H4 clustered histone 1 | B2R4R0 | 2.80E-04 | 2.69 |
| ENSG00000104824 | HNRNPL | heterogeneous nuclear ribonucleoprotein L | P14866 | 2.40E-04 | 2.68 |
| ENSG00000147403 | RPL10 | ribosomal protein L10 | P27635 | 3.00E-04 | 2.67 |
| ENSG00000138029 | HADHB | hydroxyacyl-CoA dehydrogenase trifunctional multienzyme complex subunit beta | P55084 | 9.40E-04 | 2.66 |
| ENSG00000179091 | CYC1 | cytochrome c1 | P08574 | 1.70E-04 | 2.63 |
| ENSG00000105971 | CAV2 | caveolin 2 | P51636 | 6.70E-04 | 2.62 |
| ENSG00000182774 | RPS17 | ribosomal protein S17 | P08708 | 0.002 | 2.62 |
| ENSG00000101160 | CTSZ | cathepsin Z | Q9UBR2 | 6.30E-04 | 2.61 |
| ENSG00000138069 | RAB1A | RAB1A, member RAS oncogene family | P62820 | 0.0014 | 2.59 |
| ENSG00000165283 | STOML2 | stomatin like 2 | Q9UJZ1 | 9.80E-04 | 2.58 |
| ENSG00000072778 | ACADVL | acyl-CoA dehydrogenase very long chain | P49748 | 2.50E-04 | 2.56 |
| ENSG00000085491 | SLC25A24 | solute carrier family 25 member 24 | Q6NUK1 | 0.013 | 2.56 |
| ENSG00000079805 | DNM2 | dynamin 2 | P50570 | 0.015 | 2.55 |
| ENSG00000087460 | GNAS | GNAS complex locus | A0A0S2Z3H8 | 2.80E-04 | 2.55 |
| ENSG00000203879 | GDI1 | GDP dissociation inhibitor 1 | A0A0S2Z3X8 | 4.70E-04 | 2.53 |
| ENSG00000167792 | NDUFV1 | NADH:ubiquinone oxidoreductase core subunit V1 | P49821 | 6.40E-04 | 2.5 |
| ENSG00000124767 | GLO1 | glyoxalase I | Q04760 | 0.0019 | 2.47 |
| ENSG00000004455 | AK2 | adenylate kinase 2 | P54819 | 5.20E-04 | 2.46 |
| ENSG00000184009 | ACTG1 | actin gamma 1 | P63261 | 0.007 | 2.45 |
| ENSG00000031698 | SARS1 | seryl-tRNA synthetase 1 | Q5T5C7 | 3.60E-04 | 2.45 |
| ENSG00000105379 | ETFB | electron transfer flavoprotein subunit beta | P38117 | 3.80E-04 | 2.4 |
| ENSG00000161057 | PSMC2 | proteasome 26S subunit, ATPase 2 | B7Z571 | 4.90E-04 | 2.4 |
| ENSG00000165799 | RNASE7 | ribonuclease A family member 7 | Q9H1E1 | 0.026 | 2.4 |
| ENSG00000113407 | TARS1 | threonyl-tRNA synthetase 1 | P26639 | 0.0014 | 2.4 |
| ENSG00000185825 | BCAP31 | B cell receptor associated protein 31 | P51572 | 8.70E-04 | 2.38 |
| ENSG00000165637 | VDAC2 | voltage dependent anion channel 2 | P45880 | 5.70E-04 | 2.38 |
| ENSG00000088832 | FKBP1A | FKBP prolyl isomerase 1A | P62942 | 7.20E-04 | 2.37 |
| ENSG00000071082 | RPL31 | ribosomal protein L31 | P62899 | 0.009 | 2.36 |

| | | | | | |
|---|---|---|---|---|---|
| ENSG00000117519 | CNN3 | calponin 3 | Q15417 | 9.60E-04 | 2.34 |
| ENSG00000198712 | MT-CO2 | mitochondrially encoded cytochrome c oxidase II | P00403 | 0.002 | 2.34 |
| ENSG00000240386 | LCE1F | late cornified envelope 1F | Q5T754 | 0.028 | 2.33 |
| ENSG00000126432 | PRDX5 | peroxiredoxin 5 | P30044 | 0.016 | 2.33 |
| ENSG00000142676 | RPL11 | ribosomal protein L11 | P62913 | 9.20E-04 | 2.32 |
| ENSG00000109475 | RPL34 | ribosomal protein L34 | A0A024RDH8 | 0.0014 | 2.32 |
| ENSG00000139684 | ESD | esterase D | A0A140VJJ2 | 0.0089 | 2.31 |
| ENSG00000104388 | RAB2A | RAB2A, member RAS oncogene family | P61019 | 8.70E-04 | 2.31 |
| ENSG00000265681 | RPL17 | ribosomal protein L17 | A0A024R261 | 0.0064 | 2.3 |
| ENSG00000175793 | SFN | stratifin | P31947 | 0.0042 | 2.29 |
| ENSG00000088247 | KHSRP | KH-type splicing regulatory protein | Q92945 | 9.00E-04 | 2.28 |
| ENSG00000198363 | ASPH | aspartate beta-hydroxylase | Q12797 | 8.90E-04 | 2.26 |
| ENSG00000070831 | CDC42 | cell division cycle 42 | A0A024RAE4 | 4.20E-04 | 2.23 |
| ENSG00000172053 | QARS1 | glutaminyl-tRNA synthetase 1 | B7Z840 | 6.20E-04 | 2.22 |
| ENSG00000244038 | DDOST | dolichyl-diphosphooligosaccharide--protein glycosyltransferase non-catalytic subunit | A0A024RAD5 | 0.0041 | 2.18 |
| ENSG00000105974 | CAV1 | caveolin 1 | A0A024R757 | 7.80E-04 | 2.17 |
| ENSG00000092841 | MYL6 | myosin light chain 6 | P60660 | 0.0033 | 2.17 |
| ENSG00000161547 | SRSF2 | serine and arginine rich splicing factor 2 | A0A024R8U5 | 0.0053 | 2.17 |
| ENSG00000111275 | ALDH2 | aldehyde dehydrogenase 2 family member | P05091 | 0.0023 | 2.16 |
| ENSG00000100030 | MAPK1 | mitogen-activated protein kinase 1 | P28482 | 0.0014 | 2.16 |
| ENSG00000138674 | SEC31A | SEC31 homolog A, COPII coat complex component | O94979 | 0.0036 | 2.15 |
| ENSG00000174444 | RPL4 | ribosomal protein L4 | P36578 | 6.60E-04 | 2.13 |
| ENSG00000108298 | RPL19 | ribosomal protein L19 | P84098 | 0.0016 | 2.12 |
| ENSG00000123416 | TUBA1B | tubulin alpha 1b | P68363 | 0.023 | 2.11 |
| ENSG00000197930 | ERO1A | endoplasmic reticulum oxidoreductase 1 alpha | Q96HE7 | 0.0035 | 2.1 |
| ENSG00000090863 | GLG1 | golgi glycoprotein 1 | Q92896 | 0.0013 | 2.1 |
| ENSG00000077147 | TM9SF3 | transmembrane 9 superfamily member 3 | A0A024QYS2 | 9.80E-04 | 2.09 |
| ENSG00000140740 | UQCRC2 | ubiquinol-cytochrome c reductase core protein 2 | P22695 | 0.0016 | 2.08 |
| ENSG00000135624 | CCT7 | chaperonin containing TCP1 subunit 7 | Q99832 | 0.0038 | 2.05 |
| ENSG00000064666 | CNN2 | calponin 2 | B4DDF4 | 0.029 | 2.05 |
| ENSG00000178035 | IMPDH2 | inosine monophosphate dehydrogenase 2 | P12268 | 0.0028 | 2.05 |
| ENSG00000165092 | ALDH1A1 | aldehyde dehydrogenase 1 family member A1 | P00352 | 8.60E-04 | 2.04 |
| ENSG00000023228 | NDUFS1 | NADH:ubiquinone oxidoreductase core subunit S1 | P28331 | 9.50E-04 | 2.03 |
| ENSG00000186468 | RPS23 | ribosomal protein S23 | A8K517 | 0.014 | 2.03 |
| ENSG00000142910 | TINAGL1 | tubulointerstitial nephritis antigen like 1 | Q9GZM7 | 0.0011 | 2.03 |
| ENSG00000075624 | ACTB | actin beta | P60709 | 0.001 | 2.01 |

| | | | | | |
|---|---|---|---|---|---|
| ENSG00000184743 | ATL3 | atlastin GTPase 3 | B4DXC4 | 0.0011 | 2.01 |
| ENSG00000196230 | TUBB | tubulin beta class I | B4DY90 | 0.013 | 2.01 |
| ENSG00000115541 | HSPE1 | heat shock protein family E (Hsp10) member 1 | P61604 | 0.0019 | 2 |
| ENSG00000197958 | RPL12 | ribosomal protein L12 | P30050 | 0.001 | 2 |
| ENSG00000128335 | APOL2 | apolipoprotein L2 | A0A024R1M8 | 0.012 | 1.98 |
| ENSG00000261150 | EPPK1 | epiplakin 1 | A0A087X1U6 | 0.031 | 1.98 |
| ENSG00000182199 | SHMT2 | serine hydroxymethyltransferase 2 | P34897 | 0.0032 | 1.98 |
| ENSG00000177688 | SUMO4 | small ubiquitin like modifier 4 | Q6EEV6 | 0.026 | 1.97 |
| ENSG00000133731 | IMPA1 | inositol monophosphatase 1 | A0A140VJL8 | 0.0089 | 1.96 |
| ENSG00000101150 | TPD52L2 | TPD52 like 2 | O43399 | 0.032 | 1.96 |
| ENSG00000143870 | PDIA6 | protein disulfide isomerase family A member 6 | Q15084 | 0.0036 | 1.95 |
| ENSG00000182809 | CRIP2 | cysteine rich protein 2 | P52943 | 0.0015 | 1.94 |
| ENSG00000170323 | FABP4 | fatty acid binding protein 4 | E7DVW4 | 0.002 | 1.94 |
| ENSG00000211677 | IGLC2 | immunoglobulin lambda constant 2 | P0DOY2 | 0.014 | 1.94 |
| ENSG00000197956 | S100A6 | S100 calcium binding protein A6 | P06703 | 0.0019 | 1.94 |
| ENSG00000175130 | MARCKSL1 | MARCKS like 1 | P49006 | 0.0037 | 1.92 |
| ENSG00000148834 | GSTO1 | glutathione S-transferase omega 1 | P78417 | 0.0023 | 1.9 |
| ENSG00000108671 | PSMD11 | proteasome 26S subunit, non-ATPase 11 | O00231 | 0.0013 | 1.9 |
| ENSG00000168385 | SEPTIN2 | septin 2 | Q15019 | 0.013 | 1.9 |
| ENSG00000122545 | SEPTIN7 | septin 7 | A8K3D0 | 0.003 | 1.9 |
| ENSG00000057757 | PITHD1 | PITH domain containing 1 | Q9GZP4 | 0.0019 | 1.88 |
| ENSG00000233927 | RPS28 | ribosomal protein S28 | B2R4R9 | 0.03 | 1.88 |
| ENSG00000167526 | RPL13 | ribosomal protein L13 | A8K4C8 | 0.0024 | 1.86 |
| ENSG00000086598 | TMED2 | transmembrane p24 trafficking protein 2 | Q15363 | 0.014 | 1.86 |
| ENSG00000165280 | VCP | valosin containing protein | P55072 | 0.0032 | 1.86 |
| ENSG00000112306 | RPS12 | ribosomal protein S12 | P25398 | 0.0031 | 1.85 |
| ENSG00000108774 | RAB5C | RAB5C, member RAS oncogene family | P51148 | 0.0096 | 1.84 |
| ENSG00000138385 | SSB | small RNA binding exonuclease protection factor La | P05455 | 0.0018 | 1.84 |
| ENSG00000126524 | SBDS | SBDS ribosome maturation factor | A0A0S2Z5I7 | 0.003 | 1.83 |
| ENSG00000140612 | SEC11A | SEC11 homolog A, signal peptidase complex subunit | P67812 | 0.022 | 1.83 |
| ENSG00000176986 | SEC24C | SEC24 homolog C, COPII coat complex component | A0A024QZM6 | 0.017 | 1.83 |
| ENSG00000023318 | ERP44 | endoplasmic reticulum protein 44 | Q9BS26 | 0.0021 | 1.82 |
| ENSG00000100034 | PPM1F | protein phosphatase, Mg2+/Mn2+ dependent 1F | P49593 | 0.048 | 1.82 |
| ENSG00000067560 | RHOA | ras homolog family member A | A0A024R324 | 0.0014 | 1.82 |
| ENSG00000177600 | RPLP2 | ribosomal protein lateral stalk subunit P2 | A0A024RCA7 | 0.0071 | 1.82 |
| ENSG00000144848 | ATG3 | autophagy related 3 | Q9NT62 | 0.0019 | 1.81 |
| ENSG00000114867 | EIF4G1 | eukaryotic translation initiation factor 4 gamma 1 | B2RU06 | 0.0023 | 1.8 |
| ENSG00000106803 | SEC61B | SEC61 translocon subunit beta | P60468 | 0.0059 | 1.8 |

| | | | | | |
|---|---|---|---|---|---|
| ENSG00000058668 | ATP2B4 | ATPase plasma membrane Ca2+ transporting 4 | P23634 | 0.003 | 1.79 |
| ENSG00000214078 | CPNE1 | copine 1 | B0QZ18 | 0.042 | 1.77 |
| ENSG00000144713 | RPL32 | ribosomal protein L32 | A0A024R2G7 | 0.0022 | 1.77 |
| ENSG00000101474 | APMAP | adipocyte plasma membrane associated protein | Q9HDC9 | 0.0041 | 1.76 |
| ENSG00000150753 | CCT5 | chaperonin containing TCP1 subunit 5 | B4DX08 | 0.0029 | 1.76 |
| ENSG00000126457 | PRMT1 | protein arginine methyltransferase 1 | A0A087X1W2 | 0.0043 | 1.76 |
| ENSG00000069329 | VPS35 | VPS35 retromer complex component | Q96QK1 | 0.0037 | 1.76 |
| ENSG00000159176 | CSRP1 | cysteine and glycine rich protein 1 | B4DY28 | 0.0026 | 1.75 |
| ENSG00000137509 | PRCP | prolylcarboxypeptidase | B7Z7Q6 | 0.043 | 1.75 |
| ENSG00000130429 | ARPC1B | actin related protein 2/3 complex subunit 1B | A4D275 | 0.002 | 1.74 |
| ENSG00000136688 | IL36G | interleukin 36 gamma | Q9NZH8 | 0.023 | 1.74 |
| ENSG00000132341 | RAN | RAN, member RAS oncogene family | B4DV51 | 0.0032 | 1.74 |
| ENSG00000126602 | TRAP1 | TNF receptor associated protein 1 | Q12931 | 0.025 | 1.73 |
| ENSG00000213585 | VDAC1 | voltage dependent anion channel 1 | A0A1L1UHR1 | 0.0093 | 1.73 |
| ENSG00000110880 | CORO1C | coronin 1C | Q9ULV4 | 0.01 | 1.72 |
| ENSG00000104529 | EEF1D | eukaryotic translation elongation factor 1 delta | B2RAR6 | 0.0069 | 1.71 |
| ENSG00000239672 | NME1 | NME/NM23 nucleoside diphosphate kinase 1 | P15531 | 0.01 | 1.71 |
| ENSG00000168906 | MAT2A | methionine adenosyltransferase 2A | A0A140VJP5 | 0.013 | 1.7 |
| ENSG00000122406 | RPL5 | ribosomal protein L5 | A2RUM7 | 0.0035 | 1.7 |
| ENSG00000138768 | USO1 | USO1 vesicle transport factor | O60763 | 0.026 | 1.7 |
| ENSG00000111530 | CAND1 | cullin associated and neddylation dissociated 1 | Q86VP6 | 0.004 | 1.69 |
| ENSG00000134440 | NARS1 | asparaginyl-tRNA synthetase 1 | O43776 | 0.022 | 1.69 |
| ENSG00000189221 | MAOA | monoamine oxidase A | P21397 | 0.0046 | 1.68 |
| ENSG00000131828 | PDHA1 | pyruvate dehydrogenase E1 subunit alpha 1 | A0A024RBX9 | 0.013 | 1.68 |
| ENSG00000172354 | GNB2 | G protein subunit beta 2 | P62879 | 0.013 | 1.67 |
| ENSG00000100387 | RBX1 | ring-box 1 | P62877 | 0.0054 | 1.67 |
| ENSG00000136942 | RPL35 | ribosomal protein L35 | P42766 | 0.023 | 1.66 |
| ENSG00000072210 | ALDH3A2 | aldehyde dehydrogenase 3 family member A2 | P51648 | 0.0027 | 1.65 |
| ENSG00000117592 | PRDX6 | peroxiredoxin 6 | P30041 | 0.0034 | 1.64 |
| ENSG00000164733 | CTSB | cathepsin B | Q5HYG5 | 0.0039 | 1.62 |
| ENSG00000273749 | CYFIP1 | cytoplasmic FMR1 interacting protein 1 | Q7L576 | 0.0083 | 1.61 |
| ENSG00000116560 | SFPQ | splicing factor proline and glutamine rich | P23246 | 0.014 | 1.61 |
| ENSG00000110108 | TMEM109 | transmembrane protein 109 | Q9BVC6 | 0.018 | 1.6 |
| ENSG00000177469 | CAVIN1 | caveolae associated protein 1 | Q6NZI2 | 0.0068 | 1.59 |
| ENSG00000021355 | SERPINB1 | serpin family B member 1 | P30740 | 0.0056 | 1.59 |
| ENSG00000099797 | TECR | trans-2,3-enoyl-CoA reductase | B3KSQ1 | 0.022 | 1.59 |
| ENSG00000172757 | CFL1 | cofilin 1 | P23528 | 0.0053 | 1.58 |

| | | | | | | |
|---|---|---|---|---|---|---|
| ENSG00000119396 | RAB14 | RAB14, member RAS oncogene family | A0A024R845 | 0.017 | | 1.58 |
| ENSG00000075785 | RAB7A | RAB7A, member RAS oncogene family | A0A158RFU6 | 0.008 | | 1.58 |
| ENSG00000275895 | U2AF1L5 | U2 small nuclear RNA auxiliary factor 1-like 5 | B5BU08 | 0.0091 | | 1.57 |
| ENSG00000111640 | GAPDH | glyceraldehyde-3-phosphate dehydrogenase | P04406 | 0.0042 | | 1.56 |
| ENSG00000111144 | LTA4H | leukotriene A4 hydrolase | A0A140VK27 | 0.0071 | | 1.56 |
| ENSG00000130755 | GMFG | glia maturation factor gamma | M0R1D2 | 0.0056 | | 1.55 |
| ENSG00000150593 | PDCD4 | programmed cell death 4 | B4DKX4 | 0.026 | | 1.55 |
| ENSG00000152952 | PLOD2 | procollagen-lysine,2-oxoglutarate 5-dioxygenase 2 | O00469 | 0.0051 | | 1.55 |
| ENSG00000158710 | TAGLN2 | transgelin 2 | P37802 | 0.006 | | 1.55 |
| ENSG00000205542 | TMSB4X | thymosin beta 4 X-linked | A2VCK8 | 0.035 | | 1.55 |
| ENSG00000124557 | BTN1A1 | butyrophilin subfamily 1 member A1 | Q13410 | 0.0076 | | 1.54 |
| ENSG00000102393 | GLA | galactosidase alpha | P06280 | 0.0083 | | 1.54 |
| ENSG00000116251 | RPL22 | ribosomal protein L22 | P35268 | 0.0053 | | 1.54 |
| ENSG00000084774 | CAD | carbamoyl-phosphate synthetase 2, aspartate transcarbamylase, and dihydroorotase | F8VPD4 | 0.045 | | 1.52 |
| ENSG00000130811 | EIF3G | eukaryotic translation initiation factor 3 subunit G | O75821 | 0.0091 | | 1.52 |
| ENSG00000078369 | GNB1 | G protein subunit beta 1 | B3KVK2 | 0.02 | | 1.52 |
| ENSG00000100028 | SNRPD3 | small nuclear ribonucleoprotein D3 polypeptide | P62318 | 0.0079 | | 1.52 |
| ENSG00000206503 | HLA-A | major histocompatibility complex, class I, A | P30443 | 0.0044 | | 1.51 |
| ENSG00000146731 | CCT6A | chaperonin containing TCP1 subunit 6A | P40227 | 0.0097 | | 1.49 |
| ENSG00000167656 | LY6D | lymphocyte antigen 6 family member D | Q14210 | 0.036 | | 1.49 |
| ENSG00000147604 | RPL7 | ribosomal protein L7 | P18124 | 0.0059 | | 1.48 |
| ENSG00000128245 | YWHAH | tyrosine 3-monooxygenase/tryptophan 5-monooxygenase activation protein eta | A0A024R1K7 | 0.026 | | 1.48 |
| ENSG00000163466 | ARPC2 | actin related protein 2/3 complex subunit 2 | O15144 | 0.016 | | 1.47 |
| ENSG00000129351 | ILF3 | interleukin enhancer binding factor 3 | Q12906 | 0.0071 | | 1.47 |
| ENSG00000058262 | SEC61A1 | SEC61 translocon subunit alpha 1 | B3KME8 | 0.015 | | 1.46 |
| ENSG00000085063 | CD59 | CD59 molecule (CD59 blood group) | P13987 | 0.012 | | 1.44 |
| ENSG00000134333 | LDHA | lactate dehydrogenase A | P00338 | 0.014 | | 1.44 |
| ENSG00000015479 | MATR3 | matrin 3 | A0A0R4J2E8 | 0.022 | 1.43 | |
| ENSG00000141522 | ARHGDIA | Rho GDP dissociation inhibitor alpha | P52565 | 0.03 | 1.42 | |
| ENSG00000179085 | DPM3 | dolichyl-phosphate mannosyltransferase subunit 3, regulatory | A0A140VJI4 | 0.015 | 1.42 | |
| ENSG00000125868 | DSTN | destrin, actin depolymerizing factor | P60981 | 0.0082 | 1.42 | |
| ENSG00000239264 | TXNDC5 | thioredoxin domain containing 5 | A0A024QZV0 | 0.019 | 1.42 | |
| ENSG00000145425 | RPS3A | ribosomal protein S3A | P61247 | 0.018 | 1.41 | |

| | | | | | |
|---|---|---|---|---|---|
| ENSG00000196419 | XRCC6 | X-ray repair cross complementing 6 | A0A024R1N4 | 0.041 | 1.41 |
| ENSG00000134308 | YWHAQ | tyrosine 3-monooxygenase/tryptophan 5-monooxygenase activation protein theta | P27348 | 0.0093 | 1.41 |
| ENSG00000133313 | CNDP2 | carnosine dipeptidase 2 | Q96KP4 | 0.0093 | 1.4 |
| ENSG00000108654 | DDX5 | DEAD-box helicase 5 | P17844 | 0.0095 | 1.4 |
| ENSG00000259207 | ITGB3 | integrin subunit beta 3 | P05106 | 0.048 | 1.4 |
| ENSG00000107438 | PDLIM1 | PDZ and LIM domain 1 | O00151 | 0.018 | 1.4 |
| ENSG00000108107 | RPL28 | ribosomal protein L28 | P46779 | 0.04 | 1.4 |
| ENSG00000124939 | SCGB2A1 | secretoglobin family 2A member 1 | O75556 | 0.047 | 1.4 |
| ENSG00000108828 | VAT1 | vesicle amine transport 1 | A0A024R1Z6 | 0.037 | 1.4 |
| ENSG00000083845 | RPS5 | ribosomal protein S5 | A0A024R4Q8 | 0.024 | 1.38 |
| ENSG00000153187 | HNRNPU | heterogeneous nuclear ribonucleoprotein U | Q00839 | 0.02 | 1.37 |
| ENSG00000142864 | SERBP1 | SERPINE1 mRNA binding protein 1 | Q8NC51 | 0.039 | 1.36 |
| ENSG00000213719 | CLIC1 | chloride intracellular channel 1 | O00299 | 0.017 | 1.35 |
| ENSG00000106105 | GARS1 | glycyl-tRNA synthetase 1 | A0A090N8G0 | 0.047 | 1.35 |
| ENSG00000118680 | MYL12B | myosin light chain 12B | O14950 | 0.0089 | 1.35 |
| ENSG00000149273 | RPS3 | ribosomal protein S3 | P23396 | 0.025 | 1.35 |
| ENSG00000130402 | ACTN4 | actinin alpha 4 | A0A0S2Z3G9 | 0.0097 | 1.34 |
| ENSG00000148672 | GLUD1 | glutamate dehydrogenase 1 | P00367 | 0.015 | 1.34 |
| ENSG00000168028 | RPSA | ribosomal protein SA | A0A0C4DG17 | 0.015 | 1.34 |
| ENSG00000100823 | APEX1 | apurinic/apyrimidinic endodeoxyribonuclease 1 | P27695 | 0.011 | 1.33 |
| ENSG00000167460 | TPM4 | tropomyosin 4 | P67936 | 0.028 | 1.33 |
| ENSG00000254772 | EEF1G | eukaryotic translation elongation factor 1 gamma | P26641 | 0.0097 | 1.32 |
| ENSG00000100097 | LGALS1 | galectin 1 | P09382 | 0.015 | 1.32 |
| ENSG00000089157 | RPLP0 | ribosomal protein lateral stalk subunit P0 | A0A024RBS2 | 0.01 | 1.32 |
| ENSG00000104763 | ASAH1 | N-acylsphingosine amidohydrolase 1 | A8K0B6 | 0.031 | 1.31 |
| ENSG00000136160 | EDNRB | endothelin receptor type B | P24530 | 0.031 | 1.29 |
| ENSG00000196531 | NACA | nascent polypeptide associated complex subunit alpha | A0A024RB41 | 0.021 | 1.29 |
| ENSG00000067113 | PLPP1 | phospholipid phosphatase 1 | A0A024QZS3 | 0.047 | 1.29 |
| ENSG00000105568 | PPP2R1A | protein phosphatase 2 scaffold subunit Aalpha | A8K7B7 | 0.047 | 1.29 |
| ENSG00000134871 | COL4A2 | collagen type IV alpha 2 chain | A0A024RDW8 | 0.029 | 1.28 |
| ENSG00000145349 | CAMK2D | calcium/calmodulin dependent protein kinase II delta | A0A024RDK3 | 0.031 | 1.27 |
| ENSG00000167468 | GPX4 | glutathione peroxidase 4 | P36969 | 0.039 | 1.27 |
| ENSG00000103415 | HMOX2 | heme oxygenase 2 | P30519 | 0.012 | 1.27 |
| ENSG00000162909 | CAPN2 | calpain 2 | B4DN77 | 0.032 | 1.25 |
| ENSG00000137154 | RPS6 | ribosomal protein S6 | A2A3R6 | 0.019 | 1.25 |
| ENSG00000108518 | PFN1 | profilin 1 | P07737 | 0.047 | 1.24 |
| ENSG00000169504 | CLIC4 | chloride intracellular channel 4 | Q6FIC5 | 0.012 | 1.23 |

| | | | | | |
|---|---|---|---|---|---|
| ENSG00000138095 | LRPPRC | leucine rich pentatricopeptide repeat containing | E5KNY5 | 0.023 | 1.23 |
| ENSG00000089009 | RPL6 | ribosomal protein L6 | A0A024RBK3 | 0.025 | 1.23 |
| ENSG00000164924 | YWHAZ | tyrosine 3-monooxygenase/tryptophan 5-monooxygenase activation protein zeta | D0PNI1 | 0.015 | 1.23 |
| ENSG00000128591 | FLNC | filamin C | Q14315 | 0.013 | 1.22 |
| ENSG00000148484 | RSU1 | Ras suppressor protein 1 | Q15404 | 0.02 | 1.22 |
| ENSG00000100934 | SEC23A | SEC23 homolog A, COPII coat complex component | Q15436 | 0.025 | 1.22 |
| ENSG00000204628 | RACK1 | receptor for activated C kinase 1 | E9KL35 | 0.028 | 1.21 |
| ENSG00000100714 | MTHFD1 | methylenetetrahydrofolate dehydrogenase, cyclohydrolase and formyltetrahydrofolate synthetase 1 | P11586 | 0.021 | 1.2 |
| ENSG00000135636 | DYSF | dysferlin | O75923 | 0.014 | 1.19 |
| ENSG00000161960 | EIF4A1 | eukaryotic translation initiation factor 4A1 | P60842 | 0.021 | 1.19 |
| ENSG00000185624 | P4HB | prolyl 4-hydroxylase subunit beta | A0A024R8S5 | 0.034 | 1.19 |
| ENSG00000079246 | XRCC5 | X-ray repair cross complementing 5 | P13010 | 0.035 | 1.19 |
| ENSG00000129562 | DAD1 | defender against cell death 1 | P61803 | 0.047 | 1.18 |
| ENSG00000168374 | ARF4 | ADP ribosylation factor 4 | P18085 | 0.036 | 1.16 |
| ENSG00000092964 | DPYSL2 | dihydropyrimidinase like 2 | A0A1C7CYX9 | 0.026 | 1.16 |
| ENSG00000196586 | MYO6 | myosin VI | Q9UM54 | 0.026 | 1.16 |
| ENSG00000060138 | YBX3 | Y-box binding protein 3 | A0A024RAV4 | 0.043 | 1.16 |
| ENSG00000171314 | PGAM1 | phosphoglycerate mutase 1 | B7Z9E5 | 0.019 | 1.15 |
| ENSG00000004700 | RECQL | RecQ like helicase | A0A024RAV2 | 0.047 | 1.15 |
| ENSG00000163468 | CCT3 | chaperonin containing TCP1 subunit 3 | B3KX11 | 0.02 | 1.14 |
| ENSG00000100316 | RPL3 | ribosomal protein L3 | P39023 | 0.039 | 1.14 |
| ENSG00000075415 | SLC25A3 | solute carrier family 25 member 3 | A0A024RBE8 | 0.039 | 1.14 |
| ENSG00000174437 | ATP2A2 | ATPase sarcoplasmic/endoplasmic reticulum Ca2+ transporting 2 | P16615 | 0.031 | 1.12 |
| ENSG00000196205 | EEF1A1P5 | eukaryotic translation elongation factor 1 alpha 1 pseudogene 5 | Q5VTE0 | 0.046 | 1.11 |
| ENSG00000120694 | HSPH1 | heat shock protein family H (Hsp110) member 1 | A0A024RDQ0 | 0.034 | 1.11 |
| ENSG00000175166 | PSMD2 | proteasome 26S subunit ubiquitin receptor, non-ATPase 2 | Q13200 | 0.032 | 1.11 |
| ENSG00000170759 | KIF5B | kinesin family member 5B | P33176 | 0.046 | 1.1 |
| ENSG00000092010 | PSME1 | proteasome activator subunit 1 | Q06323 | 0.023 | 1.1 |
| ENSG00000197157 | SND1 | staphylococcal nuclease and tudor domain containing 1 | A0A140VK49 | 0.031 | 1.1 |
| ENSG00000135829 | DHX9 | DExH-box helicase 9 | B3KU66 | 0.025 | 1.09 |
| ENSG00000100401 | RANGAP1 | Ran GTPase activating protein 1 | A0A024R1U0 | 0.044 | 1.09 |
| ENSG00000198959 | TGM2 | transglutaminase 2 | P21980 | 0.02 | 1.09 |
| ENSG00000130741 | EIF2S3 | eukaryotic translation initiation factor 2 subunit gamma | P41091 | 0.044 | 1.08 |
| ENSG00000137106 | GRHPR | glyoxylate and hydroxypyruvate reductase | Q9UBQ7 | 0.021 | 1.08 |
| ENSG00000142798 | HSPG2 | heparan sulfate proteoglycan 2 | P98160 | 0.027 | 1.08 |

| | | | | | |
|---|---|---|---|---|---|
| ENSG00000105835 | NAMPT | nicotinamide phosphoribosyltransferase | A0A024R718 | 0.032 | 1.07 |
| ENSG00000024422 | EHD2 | EH domain containing 2 | A0A024R0S6 | 0.029 | 1.06 |
| ENSG00000187514 | PTMA | prothymosin alpha | P06454 | 0.046 | 1.04 |
| ENSG00000189171 | S100A13 | S100 calcium binding protein A13 | Q99584 | 0.033 | 1.03 |
| ENSG00000096384 | HSP90AB1 | heat shock protein 90 alpha family class B member 1 | A0A024RD80 | 0.034 | 1.02 |
| ENSG00000163902 | RPN1 | ribophorin I | P04843 | 0.046 | 1.02 |
| ENSG00000138668 | HNRNPD | heterogeneous nuclear ribonucleoprotein D | Q14103 | 0.047 | 1.01 |
| ENSG00000167085 | PHB1 | prohibitin 1 | P35232 | 0.043 | 1.01 |
| ENSG00000137767 | SQOR | sulfide quinone oxidoreductase | A0A024R5X2 | 0.039 | 1.01 |
| ENSG00000110799 | VWF | von Willebrand factor | P04275 | 0.042 | 0.99 |
| ENSG00000167658 | EEF2 | eukaryotic translation elongation factor 2 | P13639 | 0.042 | 0.98 |
| ENSG00000136068 | FLNB | filamin B | O75369 | 0.04 | 0.96 |
| ENSG00000138772 | ANXA3 | annexin A3 | P12429 | 0.045 | 0.94 |
| ENSG00000127022 | CANX | calnexin | P27824 | 0.049 | 0.94 |
| ENSG00000135316 | SYNCRIP | synaptotagmin binding cytoplasmic RNA interacting protein | B7Z645 | 0.045 | 0.94 |
| ENSG00000261371 | PECAM1 | platelet and endothelial cell adhesion molecule 1 | A0A075B738 | 0.047 | 0.93 |
| ENSG00000140988 | RPS2 | ribosomal protein S2 | P15880 | 0.044 | 0.92 |
| ENSG00000077549 | CAPZB | capping actin protein of muscle Z-line subunit beta | P47756 | 0.049 | 0.87 |

Adjustments of p-values for multiple comparisons were used with Benjamini-Hochberg (BH) correction.

### Supplemental Table S3. Insulin induced palmitoylated proteins in human LECs.

| ENSEMBL ID | Gene Symbol | Description | UNIPROT | *p*-value | -Log Ratio (+/-HA) |
|---|---|---|---|---|---|
| ENSG00000054116 | TRAPPC3 | trafficking protein particle complex subunit 3 | A0A087WWM0 | 1.10E-06 | 6.86 |
| ENSG00000102007 | PLP2 | proteolipid protein 2 | A0A024QYW3 | 2.80E-05 | 6.11 |
| ENSG00000114698 | PLSCR4 | phospholipid scramblase 4 | Q9NRQ2 | 5.40E-07 | 5.97 |
| ENSG00000126458 | RRAS | RAS related | A0A024QZF2 | 7.40E-07 | 5.94 |
| ENSG00000156052 | GNAQ | G protein subunit alpha q | A0A024R240 | 8.20E-07 | 5.91 |
| ENSG00000120063 | GNA13 | G protein subunit alpha 13 | Q14344 | 2.40E-06 | 5.81 |
| ENSG00000099864 | PALM | paralemmin | A0A024R207 | 1.30E-06 | 5.69 |
| ENSG00000162618 | ADGRL4 | adhesion G protein-coupled receptor L4 | Q9HBW9 | 2.00E-06 | 5.66 |
| ENSG00000108219 | TSPAN14 | tetraspanin 14 | Q8NG11 | 3.20E-06 | 5.47 |
| ENSG00000183726 | TMEM50A | transmembrane protein 50A | O95807 | 1.20E-05 | 5.46 |
| ENSG00000111897 | SERINC1 | serine incorporator 1 | Q9NRX5 | 2.80E-06 | 5.39 |
| ENSG00000135047 | CTSL | cathepsin L | A0A024R276 | 2.00E-06 | 5.08 |
| ENSG00000153551 | CMTM7 | CKLF like MARVEL transmembrane domain containing 7 | A0A024R2L3 | 7.20E-06 | 4.95 |
| ENSG00000099203 | TMED1 | transmembrane p24 trafficking protein 1 | Q13445 | 3.30E-06 | 4.91 |
| ENSG00000243279 | PRAF2 | PRA1 domain family member 2 | A0A024QZ22 | 9.30E-06 | 4.88 |
| ENSG00000169908 | TM4SF1 | transmembrane 4 L six family member 1 | P30408 | 1.10E-05 | 4.84 |
| ENSG00000227500 | SCAMP4 | secretory carrier membrane protein 4 | Q969E2 | 4.00E-06 | 4.82 |
| ENSG00000002586 | CD99 | CD99 molecule (Xg blood group) | P14209 | 2.70E-06 | 4.78 |
| ENSG00000116754 | SRSF11 | serine and arginine rich splicing factor 11 | Q05519 | 1.10E-05 | 4.73 |
| ENSG00000249751 | ECSCR | endothelial cell surface expressed chemotaxis and apoptosis regulator | Q19T08 | 6.70E-06 | 4.65 |
| ENSG00000013297 | CLDN11 | claudin 11 | O75508 | 7.10E-06 | 4.63 |
| ENSG00000147533 | GOLGA7 | golgin A7 | Q7Z5G4 | 5.90E-06 | 4.62 |
| ENSG00000142089 | IFITM3 | interferon induced transmembrane protein 3 | Q01628 | 4.90E-06 | 4.39 |
| ENSG00000026508 | CD44 | CD44 molecule (Indian blood group) | P16070 | 4.10E-06 | 4.36 |
| ENSG00000092531 | SNAP23 | synaptosome associated protein 23 | A8K287 | 1.30E-05 | 4.34 |
| ENSG00000159140 | SON | SON DNA and RNA binding protein | P18583 | 2.70E-05 | 4.28 |
| ENSG00000177697 | CD151 | CD151 molecule (Raph blood group) | A0A024RCB3 | 1.20E-05 | 4.21 |
| ENSG00000139921 | TMX1 | thioredoxin related transmembrane protein 1 | Q9H3N1 | 7.10E-06 | 4.19 |
| ENSG00000136156 | ITM2B | integral membrane protein 2B | Q9Y287 | 2.60E-05 | 4.11 |
| ENSG00000135821 | GLUL | glutamate-ammonia ligase | A8YXX4 | 0.021 | 4.1 |
| ENSG00000140497 | SCAMP2 | secretory carrier membrane protein 2 | A8K769 | 8.40E-06 | 4.1 |
| ENSG00000088256 | GNA11 | G protein subunit alpha 11 | P29992 | 3.70E-05 | 4.08 |

| | | | | | |
|---|---|---|---|---|---|
| ENSG00000123728 | RAP2C | RAP2C, member of RAS oncogene family | Q9Y3L5 | 6.50E-06 | 4.04 |
| ENSG00000129625 | REEP5 | receptor accessory protein 5 | Q00765 | 2.30E-05 | 4.04 |
| ENSG00000213699 | SLC35F6 | solute carrier family 35 member F6 | Q8N357 | 4.00E-05 | 3.97 |
| ENSG00000133818 | RRAS2 | RAS related 2 | P62070 | 1.30E-05 | 3.84 |
| ENSG00000144959 | NCEH1 | neutral cholesterol ester hydrolase 1 | A0A0A0MTJ9 | 4.60E-05 | 3.82 |
| ENSG00000010278 | CD9 | CD9 molecule | A6NNI4 | 5.10E-05 | 3.81 |
| ENSG00000114062 | UBE3A | ubiquitin protein ligase E3A | Q05086 | 4.10E-05 | 3.79 |
| ENSG00000185651 | UBE2L3 | ubiquitin conjugating enzyme E2 L3 | P68036 | 2.80E-05 | 3.73 |
| ENSG00000057149 | SERPINB3 | serpin family B member 3 | P29508 | 0.0068 | 3.7 |
| ENSG00000132388 | UBE2G1 | ubiquitin conjugating enzyme E2 G1 | P62253 | 2.40E-04 | 3.69 |
| ENSG00000181104 | F2R | coagulation factor II thrombin receptor | P25116 | 2.70E-05 | 3.67 |
| ENSG00000004468 | CD38 | CD38 molecule | B4E006 | 8.00E-05 | 3.66 |
| ENSG00000143878 | RHOB | ras homolog family member B | P62745 | 8.60E-05 | 3.66 |
| ENSG00000100258 | LMF2 | lipase maturation factor 2 | Q9BU23 | 4.20E-05 | 3.65 |
| ENSG00000133226 | SRRM1 | serine and arginine repetitive matrix 1 | B7Z7U0 | 1.90E-05 | 3.65 |
| ENSG00000136026 | CKAP4 | cytoskeleton associated protein 4 | A0A024RBH2 | 5.50E-05 | 3.64 |
| ENSG00000110651 | CD81 | CD81 molecule | E9PJK1 | 2.00E-04 | 3.63 |
| ENSG00000108848 | LUC7L3 | LUC7 like 3 pre-mRNA splicing factor | J3KPP4 | 3.40E-05 | 3.61 |
| ENSG00000003056 | M6PR | mannose-6-phosphate receptor, cation dependent | F5GX30 | 1.70E-04 | 3.58 |
| ENSG00000148175 | STOM | stomatin | F8VSL7 | 2.40E-05 | 3.58 |
| ENSG00000104852 | SNRNP70 | small nuclear ribonucleoprotein U1 subunit 70 | P08621 | 2.70E-05 | 3.46 |
| ENSG00000213625 | LEPROT | leptin receptor overlapping transcript | A0A087X0N2 | 8.20E-05 | 3.42 |
| ENSG00000206073 | SERPINB4 | serpin family B member 4 | P48594 | 0.023 | 3.42 |
| ENSG00000076706 | MCAM | melanoma cell adhesion molecule | A0A024R3I5 | 4.20E-05 | 3.41 |
| ENSG00000188313 | PLSCR1 | phospholipid scramblase 1 | O15162 | 7.40E-05 | 3.41 |
| ENSG00000063854 | HAGH | hydroxyacylglutathione hydrolase | Q16775 | 0.022 | 3.4 |
| ENSG00000130725 | UBE2M | ubiquitin conjugating enzyme E2 M | A0A024R4T4 | 3.90E-05 | 3.37 |
| ENSG00000146411 | SLC2A12 | solute carrier family 2 member 12 | Q8TD20 | 5.50E-04 | 3.36 |
| ENSG00000049245 | VAMP3 | vesicle associated membrane protein 3 | Q15836 | 4.40E-05 | 3.36 |
| ENSG00000120437 | ACAT2 | acetyl-CoA acetyltransferase 2 | Q9BWD1 | 9.60E-05 | 3.35 |
| ENSG00000137497 | NUMA1 | nuclear mitotic apparatus protein 1 | A0A024R5M9 | 1.80E-04 | 3.35 |
| ENSG00000143222 | UFC1 | ubiquitin-fold modifier conjugating enzyme 1 | Q9Y3C8 | 1.10E-04 | 3.35 |
| ENSG00000132589 | FLOT2 | flotillin 2 | J3QLD9 | 1.20E-04 | 3.31 |
| ENSG00000078140 | UBE2K | ubiquitin conjugating enzyme E2 K | P61086 | 4.10E-05 | 3.25 |

| | | | | | |
|---|---|---|---|---|---|
| ENSG00000135124 | P2RX4 | purinergic receptor P2X 4 | Q99571 | 9.10E-05 | 3.2 |
| ENSG00000110917 | MLEC | malectin | F5GX14 | 7.10E-05 | 3.17 |
| ENSG00000112739 | PRPF4B | pre-mRNA processing factor 4B | A0A024QZY5 | 1.40E-04 | 3.17 |
| ENSG00000109084 | TMEM97 | transmembrane protein 97 | Q5BJF2 | 0.0093 | 3.16 |
| ENSG00000184470 | TXNRD2 | thioredoxin reductase 2 | E7EWK1 | 9.70E-05 | 3.14 |
| ENSG00000112531 | QKI | QKI, KH domain containing RNA binding | Q96PU8 | 1.20E-04 | 3.13 |
| ENSG00000072401 | UBE2D1 | ubiquitin conjugating enzyme E2 D1 | A0A087WW00 | 7.30E-04 | 3.11 |
| ENSG00000125148 | MT2A | metallothionein 2A | P02795 | 8.90E-04 | 3.08 |
| ENSG00000168899 | VAMP5 | vesicle associated membrane protein 5 | O95183 | 1.10E-04 | 3.04 |
| ENSG00000136688 | IL36G | interleukin 36 gamma | Q9NZH8 | 8.40E-04 | 2.96 |
| ENSG00000157227 | MMP14 | matrix metallopeptidase 14 | P50281 | 7.70E-05 | 2.96 |
| ENSG00000254087 | LYN | LYN proto-oncogene, Src family tyrosine kinase | P07948 | 2.20E-04 | 2.92 |
| ENSG00000134247 | PTGFRN | prostaglandin F2 receptor inhibitor | Q9P2B2 | 1.50E-04 | 2.91 |
| ENSG00000072274 | TFRC | transferrin receptor | P02786 | 2.60E-04 | 2.9 |
| ENSG00000145833 | DDX46 | DEAD-box helicase 46 | A0A0C4DG89 | 1.90E-04 | 2.85 |
| ENSG00000000003 | TSPAN6 | tetraspanin 6 | A0A024RCI0 | 0.0021 | 2.85 |
| ENSG00000177889 | UBE2N | ubiquitin conjugating enzyme E2 N | P61088 | 4.10E-04 | 2.8 |
| ENSG00000143546 | S100A8 | S100 calcium binding protein A8 | P05109 | 0.046 | 2.79 |
| ENSG00000139433 | GLTP | glycolipid transfer protein | A0A024RBI7 | 0.0011 | 2.78 |
| ENSG00000130985 | UBA1 | ubiquitin like modifier activating enzyme 1 | A0A024R1A3 | 1.10E-04 | 2.77 |
| ENSG00000198833 | UBE2J1 | ubiquitin conjugating enzyme E2 J1 | Q9Y385 | 3.10E-04 | 2.76 |
| ENSG00000187688 | TRPV2 | transient receptor potential cation channel subfamily V member 2 | Q9Y5S1 | 1.40E-04 | 2.73 |
| ENSG00000084674 | APOB | apolipoprotein B | P04114 | 0.0062 | 2.7 |
| ENSG00000205076 | LGALS7 | galectin 7 | P47929 | 0.041 | 2.7 |
| ENSG00000184363 | PKP3 | plakophilin 3 | Q9Y446 | 0.031 | 2.61 |
| ENSG00000184113 | CLDN5 | claudin 5 | D3DX19 | 4.80E-04 | 2.58 |
| ENSG00000146963 | LUC7L2 | LUC7 like 2, pre-mRNA splicing factor | Q9Y383 | 3.10E-04 | 2.55 |
| ENSG00000205937 | RNPS1 | RNA binding protein with serine rich domain 1 | D3DU92 | 0.0054 | 2.53 |
| ENSG00000204568 | MRPS18B | mitochondrial ribosomal protein S18B | B0S7P4 | 0.0011 | 2.52 |
| ENSG00000144848 | ATG3 | autophagy related 3 | Q9NT62 | 2.60E-04 | 2.5 |
| ENSG00000131378 | RFTN1 | raftlin, lipid raft linker 1 | Q14699 | 4.30E-04 | 2.5 |
| ENSG00000066056 | TIE1 | tyrosine kinase with immunoglobulin like and EGF like domains 1 | B4DTW8 | 0.001 | 2.47 |
| ENSG00000014257 | ACP3 | acid phosphatase 3 | P15309 | 0.035 | 2.46 |
| ENSG00000168394 | TAP1 | transporter 1, ATP binding cassette subfamily B member | A0A0S2Z5A6 | 2.80E-04 | 2.39 |
| ENSG00000004961 | HCCS | holocytochrome c synthase | A0A024RBY9 | 7.70E-04 | 2.33 |
| ENSG00000138760 | SCARB2 | scavenger receptor class B member 2 | Q14108 | 3.90E-04 | 2.32 |

| | | | | | |
|---|---|---|---|---|---|
| ENSG00000198431 | TXNRD1 | thioredoxin reductase 1 | Q16881 | 0.002 | 2.32 |
| ENSG00000101160 | CTSZ | cathepsin Z | Q9UBR2 | 0.0013 | 2.3 |
| ENSG00000129353 | SLC44A2 | solute carrier family 44 member 2 | A0A088QCU6 | 5.40E-04 | 2.3 |
| ENSG00000074696 | HACD3 | 3-hydroxyacyl-CoA dehydratase 3 | Q9P035 | 0.0012 | 2.28 |
| ENSG00000139180 | NDUFA9 | NADH:ubiquinone oxidoreductase subunit A9 | Q16795 | 0.034 | 2.27 |
| ENSG00000150768 | DLAT | dihydrolipoamide S-acetyltransferase | P10515 | 0.0011 | 2.26 |
| ENSG00000183291 | SELENOF | selenoprotein F | O60613 | 5.70E-04 | 2.26 |
| ENSG00000233276 | GPX1 | glutathione peroxidase 1 | P07203 | 5.10E-04 | 2.25 |
| ENSG00000159202 | UBE2Z | ubiquitin conjugating enzyme E2 Z | Q9H832 | 6.60E-04 | 2.24 |
| ENSG00000144118 | RALB | RAS like proto-oncogene B | A0A024RAG3 | 6.60E-04 | 2.23 |
| ENSG00000144744 | UBA3 | ubiquitin like modifier activating enzyme 3 | Q8TBC4 | 0.0015 | 2.2 |
| ENSG00000197728 | RPS26 | ribosomal protein S26 | A0A024RB14 | 0.0096 | 2.19 |
| ENSG00000065135 | GNAI3 | G protein subunit alpha i3 | P08754 | 7.80E-04 | 2.18 |
| ENSG00000170889 | RPS9 | ribosomal protein S9 | A0A024R4M0 | 0.0016 | 2.18 |
| ENSG00000017260 | ATP2C1 | ATPase secretory pathway Ca2+ transporting 1 | P98194 | 8.00E-04 | 2.16 |
| ENSG00000025708 | TYMP | thymidine phosphorylase | B2RBL3 | 0.048 | 2.16 |
| ENSG00000135218 | CD36 | CD36 molecule | A4D1B1 | 0.0014 | 2.13 |
| ENSG00000126261 | UBA2 | ubiquitin like modifier activating enzyme 2 | Q9UBT2 | 0.0011 | 2.12 |
| ENSG00000239306 | RBM14 | RNA binding motif protein 14 | A0A0S2Z567 | 0.038 | 2.09 |
| ENSG00000147419 | CCDC25 | coiled-coil domain containing 25 | G3V121 | 0.04 | 2.05 |
| ENSG00000147649 | MTDH | metadherin | A0A024R9D2 | 7.80E-04 | 2.05 |
| ENSG00000167754 | KLK5 | kallikrein related peptidase 5 | Q9Y337 | 0.0093 | 2.04 |
| ENSG00000103018 | CYB5B | cytochrome b5 type B | J3KNF8 | 0.00093 | 2.02 |
| ENSG00000138119 | MYOF | myoferlin | Q9NZM1 | 0.0016 | 2 |
| ENSG00000156026 | MCU | mitochondrial calcium uniporter | Q8NE86 | 0.0053 | 1.99 |
| ENSG00000033178 | UBA6 | ubiquitin like modifier activating enzyme 6 | A0A024RDB0 | 0.0016 | 1.93 |
| ENSG00000114353 | GNAI2 | G protein subunit alpha i2 | B3KP24 | 0.0018 | 1.86 |
| ENSG00000155380 | SLC16A1 | solute carrier family 16 member 1 | A0A024R0H1 | 0.01 | 1.86 |
| ENSG00000136160 | EDNRB | endothelin receptor type B | P24530 | 0.0066 | 1.82 |
| ENSG00000105971 | CAV2 | caveolin 2 | P51636 | 0.0056 | 1.81 |
| ENSG00000063177 | RPL18 | ribosomal protein L18 | A0A024QZD1 | 0.0015 | 1.81 |
| ENSG00000006451 | RALA | RAS like proto-oncogene A | P11233 | 0.0025 | 1.77 |
| ENSG00000087086 | FTL | ferritin light chain | P02792 | 0.0073 | 1.76 |
| ENSG00000130193 | THEM6 | thioesterase superfamily member 6 | Q8WUY1 | 0.0058 | 1.76 |
| ENSG00000177674 | AGTRAP | angiotensin II receptor associated protein | Q6RW13 | 0.026 | 1.73 |
| ENSG00000131016 | AKAP12 | A-kinase anchoring protein 12 | Q02952 | 0.0036 | 1.72 |
| ENSG00000066654 | THUMPD1 | THUMP domain containing 1 | A0A024R388 | 0.018 | 1.72 |

| | | | | | |
|---|---|---|---|---|---|
| ENSG00000206341 | HLA-H | major histocompatibility complex, class I, H (pseudogene) | P01893 | 0.043 | 1.71 |
| ENSG00000169710 | FASN | fatty acid synthase | P49327 | 0.0037 | 1.67 |
| ENSG00000073008 | PVR | PVR cell adhesion molecule | P15151 | 0.0049 | 1.65 |
| ENSG00000196329 | GIMAP5 | GTPase, IMAP family member 5 | A0A090N8P9 | 0.022 | 1.64 |
| ENSG00000115310 | RTN4 | reticulon 4 | Q9NQC3 | 0.0039 | 1.63 |
| ENSG00000141526 | SLC16A3 | solute carrier family 16 member 3 | A0A024R8U1 | 0.008 | 1.63 |
| ENSG00000127824 | TUBA4A | tubulin alpha 4a | P68366 | 0.0081 | 1.63 |
| ENSG00000113441 | LNPEP | leucyl and cystinyl aminopeptidase | Q9UIQ6 | 0.012 | 1.62 |
| ENSG00000116350 | SRSF4 | serine and arginine rich splicing factor 4 | Q08170 | 0.025 | 1.59 |
| ENSG00000130255 | RPL36 | ribosomal protein L36 | Q9Y3U8 | 0.012 | 1.55 |
| ENSG00000075239 | ACAT1 | acetyl-CoA acetyltransferase 1 | A0A140VJX1 | 0.0063 | 1.5 |
| ENSG00000158270 | COLEC12 | collectin subfamily member 12 | Q5KU26 | 0.013 | 1.48 |
| ENSG00000138029 | HADHB | hydroxyacyl-CoA dehydrogenase trifunctional multienzyme complex subunit beta | P55084 | 0.021 | 1.48 |
| ENSG00000074755 | ZZEF1 | zinc finger ZZ-type and EF-hand domain containing 1 | O43149 | 0.0096 | 1.45 |
| ENSG00000167996 | FTH1 | ferritin heavy chain 1 | A0A024R525 | 0.0071 | 1.44 |
| ENSG00000145349 | CAMK2D | calcium/calmodulin dependent protein kinase II delta | A0A024RDK3 | 0.019 | 1.43 |
| ENSG00000111640 | GAPDH | glyceraldehyde-3-phosphate dehydrogenase | P04406 | 0.0076 | 1.39 |
| ENSG00000148634 | HERC4 | HECT and RLD domain containing E3 ubiquitin protein ligase 4 | Q5GLZ8 | 0.007 | 1.36 |
| ENSG00000198561 | CTNND1 | catenin delta 1 | O60716 | 0.024 | 1.35 |
| ENSG00000144713 | RPL32 | ribosomal protein L32 | A0A024R2G7 | 0.0093 | 1.35 |
| ENSG00000133318 | RTN3 | reticulon 3 | O95197 | 0.026 | 1.35 |
| ENSG00000117298 | ECE1 | endothelin converting enzyme 1 | A0A024RAB0 | 0.013 | 1.34 |
| ENSG00000262814 | MRPL12 | mitochondrial ribosomal protein L12 | P52815 | 0.019 | 1.34 |
| ENSG00000174444 | RPL4 | ribosomal protein L4 | P36578 | 0.0096 | 1.32 |
| ENSG00000105974 | CAV1 | caveolin 1 | A0A024R757 | 0.013 | 1.3 |
| ENSG00000103415 | HMOX2 | heme oxygenase 2 | P30519 | 0.012 | 1.28 |
| ENSG00000109475 | RPL34 | ribosomal protein L34 | A0A024RDH8 | 0.029 | 1.27 |
| ENSG00000197081 | IGF2R | insulin like growth factor 2 receptor | P11717 | 0.034 | 1.19 |
| ENSG00000115159 | GPD2 | glycerol-3-phosphate dehydrogenase 2 | P43304 | 0.024 | 1.18 |
| ENSG00000198380 | GFPT1 | glutamine--fructose-6-phosphate transaminase 1 | Q06210 | 0.026 | 1.16 |
| ENSG00000166479 | TMX3 | thioredoxin related transmembrane protein 3 | Q96JJ7 | 0.027 | 1.08 |
| ENSG00000184584 | STING1 | stimulator of interferon response cGAMP interactor 1 | V5V0K2 | 0.044 | 1.07 |

| | | | | | |
|---|---|---|---|---|---|
| ENSG00000119711 | ALDH6A1 | aldehyde dehydrogenase 6 family member A1 | Q02252 | 0.036 | 1.06 |
| ENSG00000087460 | GNAS | GNAS complex locus | A0A0S2Z3H8 | 0.031 | 1.06 |
| ENSG00000137845 | ADAM10 | ADAM metallopeptidase domain 10 | A0A024R5U5 | 0.042 | 1.01 |
| ENSG00000137767 | SQOR | sulfide quinone oxidoreductase | A0A024R5X2 | 0.042 | 0.99 |
| ENSG00000161011 | SQSTM1 | sequestosome 1 | Q13501 | 0.038 | 0.94 |

Adjustments of p-values for multiple comparisons were used with Benjamini-Hochberg (BH) correction.

### Supplemental Table S4. Palmitoylated proteins in human LECs.

| ENSEMBL ID | Gene Symbol | Description | UNIPROT | *p*-value | -Log Ratio (+/- HA) |
|---|---|---|---|---|---|
| ENSG00000054116 | TRAPPC3 | trafficking protein particle complex subunit 3 | A0A087WWM0 | 2.70E-07 | 7.2 |
| ENSG00000116521 | SCAMP3 | secretory carrier membrane protein 3 | O14828 | 3.40E-07 | 6.63 |
| ENSG00000136156 | ITM2B | integral membrane protein 2B | Q9Y287 | 3.10E-07 | 6.58 |
| ENSG00000137312 | FLOT1 | flotillin 1 | O75955 | 6.90E-07 | 6.57 |
| ENSG00000114698 | PLSCR4 | phospholipid scramblase 4 | Q9NRQ2 | 5.10E-07 | 6.53 |
| ENSG00000110651 | CD81 | CD81 molecule | E9PJK1 | 3.40E-07 | 6.44 |
| ENSG00000133818 | RRAS2 | RAS related 2 | P62070 | 6.10E-07 | 6.44 |
| ENSG00000002586 | CD99 | CD99 molecule (Xg blood group) | P14209 | 3.20E-07 | 6.35 |
| ENSG00000132824 | SERINC3 | serine incorporator 3 | Q13530 | 5.10E-07 | 6.3 |
| ENSG00000213699 | SLC35F6 | solute carrier family 35 member F6 | Q8N357 | 1.00E-06 | 6.2 |
| ENSG00000169908 | TM4SF1 | transmembrane 4 L six family member 1 | P30408 | 4.50E-07 | 6.03 |
| ENSG00000177889 | UBE2N | ubiquitin conjugating enzyme E2 N | P61088 | 4.10E-07 | 6.02 |
| ENSG00000177697 | CD151 | CD151 molecule (Raph blood group) | A0A024RCB3 | 2.00E-06 | 6 |
| ENSG00000085365 | SCAMP1 | secretory carrier membrane protein 1 | A0A087WXB0 | 4.80E-05 | 5.98 |
| ENSG00000102007 | PLP2 | proteolipid protein 2 | A0A024QYW3 | 4.90E-06 | 5.97 |
| ENSG00000148175 | STOM | stomatin | F8VSL7 | 9.60E-07 | 5.96 |
| ENSG00000139921 | TMX1 | thioredoxin related transmembrane protein 1 | Q9H3N1 | 5.50E-07 | 5.95 |
| ENSG00000135047 | CTSL | cathepsin L | A0A024R276 | 6.70E-07 | 5.92 |
| ENSG00000129353 | SLC44A2 | solute carrier family 44 member 2 | A0A088QCU6 | 4.70E-06 | 5.92 |
| ENSG00000113811 | SELENOK | selenoprotein K | Q9Y6D0 | 1.50E-06 | 5.89 |
| ENSG00000076706 | MCAM | melanoma cell adhesion molecule | A0A024R3I5 | 9.40E-07 | 5.88 |
| ENSG00000142188 | TMEM50B | transmembrane protein 50B | P56557 | 7.90E-07 | 5.87 |
| ENSG00000087191 | PSMC5 | proteasome 26S subunit, ATPase 5 | P62195 | 5.70E-07 | 5.82 |
| ENSG00000142089 | IFITM3 | interferon induced transmembrane protein 3 | Q01628 | 1.30E-06 | 5.8 |
| ENSG00000134247 | PTGFRN | prostaglandin F2 receptor inhibitor | Q9P2B2 | 2.50E-06 | 5.78 |
| ENSG00000213281 | NRAS | NRAS proto-oncogene, GTPase | P01111 | 4.20E-06 | 5.77 |
| ENSG00000243279 | PRAF2 | PRA1 domain family member 2 | A0A024QZ22 | 2.90E-06 | 5.76 |
| ENSG00000010278 | CD9 | CD9 molecule | A6NNI4 | 7.00E-07 | 5.75 |
| ENSG00000092531 | SNAP23 | synaptosome associated protein 23 | A8K287 | 6.40E-07 | 5.72 |
| ENSG00000112972 | HMGCS1 | 3-hydroxy-3-methylglutaryl-CoA synthase 1 | A0A024R059 | 1.60E-06 | 5.71 |
| ENSG00000129625 | REEP5 | receptor accessory protein 5 | Q00765 | 1.30E-06 | 5.68 |
| ENSG00000187838 | PLSCR3 | phospholipid scramblase 3 | Q9NRY6 | 2.80E-06 | 5.66 |
| ENSG00000111897 | SERINC1 | serine incorporator 1 | Q9NRX5 | 1.60E-06 | 5.58 |
| ENSG00000168899 | VAMP5 | vesicle associated membrane protein 5 | O95183 | 2.80E-06 | 5.58 |
| ENSG00000078140 | UBE2K | ubiquitin conjugating enzyme E2 K | P61086 | 1.40E-06 | 5.56 |
| ENSG00000204632 | HLA-G | major histocompatibility complex, class I, G | P17693 | 3.10E-06 | 5.55 |
| ENSG00000049245 | VAMP3 | vesicle associated membrane protein 3 | Q15836 | 1.70E-06 | 5.53 |
| ENSG00000130725 | UBE2M | ubiquitin conjugating enzyme E2 M | A0A024R4T4 | 3.10E-06 | 5.47 |
| ENSG00000099864 | PALM | paralemmin | A0A024R207 | 3.00E-06 | 5.46 |

| | | | | | |
|---|---|---|---|---|---|
| ENSG00000175792 | RUVBL1 | RuvB like AAA ATPase 1 | Q9Y265 | 1.50E-06 | 5.37 |
| ENSG00000137288 | UQCC2 | ubiquinol-cytochrome c reductase complex assembly factor 2 | Q9BRT2 | 1.30E-06 | 5.34 |
| ENSG00000073060 | SCARB1 | scavenger receptor class B member 1 | Q8WTV0 | 2.60E-06 | 5.29 |
| ENSG00000143878 | RHOB | ras homolog family member B | P62745 | 6.40E-06 | 5.22 |
| ENSG00000033178 | UBA6 | ubiquitin like modifier activating enzyme 6 | A0A024RDB0 | 1.70E-06 | 5.19 |
| ENSG00000126261 | UBA2 | ubiquitin like modifier activating enzyme 2 | Q9UBT2 | 2.50E-06 | 5.14 |
| ENSG00000099282 | TSPAN15 | tetraspanin 15 | O95858 | 1.80E-05 | 5.11 |
| ENSG00000233276 | GPX1 | glutathione peroxidase 1 | P07203 | 3.20E-06 | 5.08 |
| ENSG00000144118 | RALB | RAS like proto-oncogene B | A0A024RAG3 | 2.70E-06 | 5.06 |
| ENSG00000185651 | UBE2L3 | ubiquitin conjugating enzyme E2 L3 | P68036 | 1.70E-05 | 5.04 |
| ENSG00000132470 | ITGB4 | integrin subunit beta 4 | A0A024R8T0 | 5.10E-06 | 4.92 |
| ENSG00000163466 | ARPC2 | actin related protein 2/3 complex subunit 2 | O15144 | 3.60E-06 | 4.91 |
| ENSG00000135404 | CD63 | CD63 molecule | A0A024RB05 | 2.70E-05 | 4.91 |
| ENSG00000132388 | UBE2G1 | ubiquitin conjugating enzyme E2 G1 | P62253 | 2.80E-06 | 4.89 |
| ENSG00000227500 | SCAMP4 | secretory carrier membrane protein 4 | Q969E2 | 3.30E-06 | 4.88 |
| ENSG00000204525 | HLA-C | major histocompatibility complex, class I, C | O19617 | 3.10E-05 | 4.86 |
| ENSG00000163762 | TM4SF18 | transmembrane 4 L six family member 18 | Q96CE8 | 2.10E-05 | 4.81 |
| ENSG00000108219 | TSPAN14 | tetraspanin 14 | Q8NG11 | 9.70E-06 | 4.8 |
| ENSG00000183726 | TMEM50A | transmembrane protein 50A | O95807 | 8.80E-06 | 4.77 |
| ENSG00000078369 | GNB1 | G protein subunit beta 1 | B3KVK2 | 3.00E-05 | 4.72 |
| ENSG00000132589 | FLOT2 | flotillin 2 | J3QLD9 | 1.70E-05 | 4.71 |
| ENSG00000120437 | ACAT2 | acetyl-CoA acetyltransferase 2 | Q9BWD1 | 1.50E-05 | 4.7 |
| ENSG00000184113 | CLDN5 | claudin 5 | D3DX19 | 1.50E-05 | 4.7 |
| ENSG00000166479 | TMX3 | thioredoxin related transmembrane protein 3 | Q96JJ7 | 4.60E-06 | 4.68 |
| ENSG00000087460 | GNAS | GNAS complex locus | A0A0S2Z3H8 | 1.80E-05 | 4.65 |
| ENSG00000136026 | CKAP4 | cytoskeleton associated protein 4 | A0A024RBH2 | 1.10E-05 | 4.62 |
| ENSG00000163110 | PDLIM5 | PDZ and LIM domain 5 | Q96HC4 | 9.60E-06 | 4.61 |
| ENSG00000010270 | STARD3NL | STARD3 N-terminal like | A0A024RA89 | 6.50E-05 | 4.6 |
| ENSG00000157227 | MMP14 | matrix metallopeptidase 14 | P50281 | 1.10E-04 | 4.58 |
| ENSG00000112531 | QKI | QKI, KH domain containing RNA binding | Q96PU8 | 3.30E-06 | 4.55 |
| ENSG00000112739 | PRPF4B | pre-mRNA processing factor 4B | A0A024QZY5 | 5.40E-06 | 4.52 |
| ENSG00000167996 | FTH1 | ferritin heavy chain 1 | A0A024R525 | 2.00E-05 | 4.5 |
| ENSG00000100823 | APEX1 | apurinic/apyrimidinic endodeoxyribonuclease 1 | P27695 | 1.90E-05 | 4.49 |
| ENSG00000066322 | ELOVL1 | ELOVL fatty acid elongase 1 | Q9BW60 | 1.20E-05 | 4.48 |
| ENSG00000114353 | GNAI2 | G protein subunit alpha i2 | B3KP24 | 9.30E-06 | 4.46 |
| ENSG00000133706 | LARS1 | leucyl-tRNA synthetase 1 | B4E266 | 6.10E-06 | 4.45 |
| ENSG00000006451 | RALA | RAS like proto-oncogene A | P11233 | 1.50E-05 | 4.44 |
| ENSG00000121774 | KHDRBS1 | KH RNA binding domain containing, signal transduction associated 1 | Q07666 | 4.00E-05 | 4.43 |
| ENSG00000100764 | PSMC1 | proteasome 26S subunit, ATPase 1 | P62191 | 2.00E-05 | 4.39 |
| ENSG00000037280 | FLT4 | fms related receptor tyrosine kinase 4 | P35916 | 1.50E-05 | 4.35 |

| | | | | | |
|---|---|---|---|---|---|
| ENSG00000110917 | MLEC | malectin | F5GX14 | 7.70E-06 | 4.34 |
| ENSG00000035687 | ADSS2 | adenylosuccinate synthase 2 | A0A024R5Q7 | 9.00E-06 | 4.33 |
| ENSG00000133789 | SWAP70 | switching B cell complex subunit SWAP70 | B3KUB9 | 8.50E-06 | 4.31 |
| ENSG00000131711 | MAP1B | microtubule associated protein 1B | A2BDK6 | 2.40E-05 | 4.29 |
| ENSG00000142676 | RPL11 | ribosomal protein L11 | P62913 | 9.40E-05 | 4.29 |
| ENSG00000115310 | RTN4 | reticulon 4 | Q9NQC3 | 3.10E-05 | 4.28 |
| ENSG00000101294 | HM13 | histocompatibility minor 13 | A0A0S2Z5V7 | 1.20E-05 | 4.24 |
| ENSG00000177674 | AGTRAP | angiotensin II receptor associated protein | Q6RW13 | 1.40E-05 | 4.21 |
| ENSG00000136111 | TBC1D4 | TBC1 domain family member 4 | O60343 | 1.80E-04 | 4.2 |
| ENSG00000130193 | THEM6 | thioesterase superfamily member 6 | Q8WUY1 | 2.00E-05 | 4.15 |
| ENSG00000144848 | ATG3 | autophagy related 3 | Q9NT62 | 1.50E-05 | 4.1 |
| ENSG00000198561 | CTNND1 | catenin delta 1 | O60716 | 1.70E-05 | 4.1 |
| ENSG00000138029 | HADHB | hydroxyacyl-CoA dehydrogenase trifunctional multienzyme complex subunit beta | P55084 | 1.10E-05 | 4.1 |
| ENSG00000112118 | MCM3 | minichromosome maintenance complex component 3 | B4DUQ9 | 7.40E-04 | 4.1 |
| ENSG00000198431 | TXNRD1 | thioredoxin reductase 1 | Q16881 | 1.80E-05 | 4.1 |
| ENSG00000149218 | ENDOD1 | endonuclease domain containing 1 | O94919 | 9.90E-05 | 4.05 |
| ENSG00000188313 | PLSCR1 | phospholipid scramblase 1 | O15162 | 6.90E-05 | 4.05 |
| ENSG00000101160 | CTSZ | cathepsin Z | Q9UBR2 | 4.90E-05 | 4.03 |
| ENSG00000150768 | DLAT | dihydrolipoamide S-acetyltransferase | P10515 | 4.00E-05 | 4.01 |
| ENSG00000177469 | CAVIN1 | caveolae associated protein 1 | Q6NZI2 | 3.30E-05 | 4 |
| ENSG00000126088 | UROD | uroporphyrinogen decarboxylase | P06132 | 6.70E-05 | 4 |
| ENSG00000078668 | VDAC3 | voltage dependent anion channel 3 | Q9Y277 | 1.30E-05 | 4 |
| ENSG00000126214 | KLC1 | kinesin light chain 1 | Q07866 | 1.40E-05 | 3.93 |
| ENSG00000169045 | HNRNPH1 | heterogeneous nuclear ribonucleoprotein H1 | P31943 | 3.20E-05 | 3.91 |
| ENSG00000182809 | CRIP2 | cysteine rich protein 2 | P52943 | 1.10E-04 | 3.9 |
| ENSG00000117298 | ECE1 | endothelin converting enzyme 1 | A0A024RAB0 | 1.00E-04 | 3.9 |
| ENSG00000143321 | HDGF | heparin binding growth factor | P51858 | 2.90E-05 | 3.89 |
| ENSG00000142634 | EFHD2 | EF-hand domain family member D2 | A0A024QZ77 | 5.20E-05 | 3.88 |
| ENSG00000141458 | NPC1 | NPC intracellular cholesterol transporter 1 | O15118 | 8.70E-05 | 3.88 |
| ENSG00000108671 | PSMD11 | proteasome 26S subunit, non-ATPase 11 | O00231 | 1.70E-05 | 3.88 |
| ENSG00000105971 | CAV2 | caveolin 2 | P51636 | 7.30E-05 | 3.87 |
| ENSG00000197956 | S100A6 | S100 calcium binding protein A6 | P06703 | 0.0015 | 3.87 |
| ENSG00000197756 | RPL37A | ribosomal protein L37a | P61513 | 0.014 | 3.85 |
| ENSG00000138119 | MYOF | myoferlin | Q9NZM1 | 5.10E-05 | 3.84 |
| ENSG00000179091 | CYC1 | cytochrome c1 | P08574 | 2.60E-05 | 3.83 |
| ENSG00000203879 | GDI1 | GDP dissociation inhibitor 1 | A0A0S2Z3X8 | 1.20E-05 | 3.8 |
| ENSG00000138760 | SCARB2 | scavenger receptor class B member 2 | Q14108 | 2.00E-05 | 3.8 |
| ENSG00000178695 | KCTD12 | potassium channel tetramerization domain containing 12 | A0A140VJM4 | 1.80E-05 | 3.79 |
| ENSG00000007168 | PAFAH1B1 | platelet activating factor acetylhydrolase 1b regulatory subunit 1 | P43034 | 4.90E-05 | 3.78 |

| | | | | | |
|---|---|---|---|---|---|
| ENSG00000069345 | DNAJA2 | DnaJ heat shock protein family (Hsp40) member A2 | A0A024R6S1 | 0.0013 | 3.75 |
| ENSG00000107796 | ACTA2 | actin alpha 2, smooth muscle | D2JYH4 | 8.30E-05 | 3.74 |
| ENSG00000112306 | RPS12 | ribosomal protein S12 | P25398 | 1.10E-04 | 3.74 |
| ENSG00000111737 | RAB35 | RAB35, member RAS oncogene family | Q15286 | 3.10E-05 | 3.7 |
| ENSG00000158467 | AHCYL2 | adenosylhomocysteinase like 2 | Q96HN2 | 2.60E-04 | 3.68 |
| ENSG00000159176 | CSRP1 | cysteine and glycine rich protein 1 | B4DY28 | 7.20E-05 | 3.68 |
| ENSG00000136937 | NCBP1 | nuclear cap binding protein subunit 1 | A0A024R179 | 1.20E-04 | 3.68 |
| ENSG00000159840 | ZYX | zyxin | Q15942 | 3.30E-05 | 3.68 |
| ENSG00000184584 | STING1 | stimulator of interferon response cGAMP interactor 1 | V5V0K2 | 2.50E-05 | 3.67 |
| ENSG00000165637 | VDAC2 | voltage dependent anion channel 2 | P45880 | 5.40E-05 | 3.67 |
| ENSG00000070831 | CDC42 | cell division cycle 42 | A0A024RAE4 | 3.80E-04 | 3.66 |
| ENSG00000008988 | RPS20 | ribosomal protein S20 | P60866 | 1.40E-04 | 3.65 |
| ENSG00000005194 | CIAPIN1 | cytokine induced apoptosis inhibitor 1 | Q6FI81 | 9.40E-05 | 3.63 |
| ENSG00000103335 | PIEZO1 | piezo type mechanosensitive ion channel component 1 | Q92508 | 1.90E-04 | 3.62 |
| ENSG00000167461 | RAB8A | RAB8A, member RAS oncogene family | A0A024R7I3 | 6.80E-05 | 3.61 |
| ENSG00000172757 | CFL1 | cofilin 1 | P23528 | 9.70E-05 | 3.6 |
| ENSG00000196329 | GIMAP5 | GTPase, IMAP family member 5 | A0A090N8P9 | 5.40E-05 | 3.6 |
| ENSG00000013375 | PGM3 | phosphoglucomutase 3 | O95394 | 1.70E-04 | 3.6 |
| ENSG00000144746 | ARL6IP5 | ADP ribosylation factor like GTPase 6 interacting protein 5 | A0A024R371 | 0.0017 | 3.59 |
| ENSG00000103415 | HMOX2 | heme oxygenase 2 | P30519 | 1.10E-04 | 3.59 |
| ENSG00000136160 | EDNRB | endothelin receptor type B | P24530 | 4.20E-05 | 3.58 |
| ENSG00000169564 | PCBP1 | poly(rC) binding protein 1 | Q15365 | 9.90E-05 | 3.57 |
| ENSG00000124570 | SERPINB6 | serpin family B member 6 | A0A024QZX5 | 1.10E-04 | 3.57 |
| ENSG00000124767 | GLO1 | glyoxalase I | Q04760 | 1.30E-04 | 3.55 |
| ENSG00000108774 | RAB5C | RAB5C, member RAS oncogene family | P51148 | 4.10E-05 | 3.55 |
| ENSG00000137962 | ARHGAP29 | Rho GTPase activating protein 29 | Q52LW3 | 2.30E-05 | 3.54 |
| ENSG00000164305 | CASP3 | caspase 3 | P42574 | 3.40E-04 | 3.53 |
| ENSG00000139726 | DENR | density regulated re-initiation and release factor | A0A024RBR3 | 5.10E-04 | 3.52 |
| ENSG00000142864 | SERBP1 | SERPINE1 mRNA binding protein 1 | Q8NC51 | 3.50E-04 | 3.52 |
| ENSG00000119335 | SET | SET nuclear proto-oncogene | Q01105 | 1.10E-04 | 3.52 |
| ENSG00000130985 | UBA1 | ubiquitin like modifier activating enzyme 1 | A0A024R1A3 | 1.10E-04 | 3.52 |
| ENSG00000108883 | EFTUD2 | elongation factor Tu GTP binding domain containing 2 | B3KX19 | 5.10E-05 | 3.49 |
| ENSG00000166128 | RAB8B | RAB8B, member RAS oncogene family | Q92930 | 8.90E-05 | 3.49 |
| ENSG00000120802 | TMPO | thymopoietin | A0A024RBE7 | 4.30E-04 | 3.49 |
| ENSG00000136813 | ECPAS | Ecm29 proteasome adaptor and scaffold | NA | 3.20E-04 | 3.48 |
| ENSG00000104852 | SNRNP70 | small nuclear ribonucleoprotein U1 subunit 70 | P08621 | 4.80E-05 | 3.47 |
| ENSG00000141756 | FKBP10 | FKBP prolyl isomerase 10 | A0A024R1W3 | 2.60E-04 | 3.45 |
| ENSG00000067704 | IARS2 | isoleucyl-tRNA synthetase 2, mitochondrial | Q9NSE4 | 1.00E-04 | 3.45 |

| | | | | | |
|---|---|---|---|---|---|
| ENSG00000198682 | PAPSS2 | 3'-phosphoadenosine 5'-phosphosulfate synthase 2 | O95340 | 5.30E-05 | 3.45 |
| ENSG00000158195 | WASF2 | WASP family member 2 | Q9Y6W5 | 0.0014 | 3.45 |
| ENSG00000239672 | NME1 | NME/NM23 nucleoside diphosphate kinase 1 | P15531 | 7.60E-05 | 3.44 |
| ENSG00000165434 | PGM2L1 | phosphoglucomutase 2 like 1 | Q6PCE3 | 1.00E-04 | 3.44 |
| ENSG00000072274 | TFRC | transferrin receptor | P02786 | 2.90E-04 | 3.44 |
| ENSG00000114942 | EEF1B2 | eukaryotic translation elongation factor 1 beta 2 | A0A024R3W7 | 1.40E-04 | 3.43 |
| ENSG00000240972 | MIF | macrophage migration inhibitory factor | I4AY87 | 6.70E-05 | 3.43 |
| ENSG00000184009 | ACTG1 | actin gamma 1 | P63261 | 9.70E-05 | 3.42 |
| ENSG00000135269 | TES | testin LIM domain protein | A4D0U5 | 5.10E-05 | 3.4 |
| ENSG00000131016 | AKAP12 | A-kinase anchoring protein 12 | Q02952 | 5.10E-05 | 3.39 |
| ENSG00000137497 | NUMA1 | nuclear mitotic apparatus protein 1 | A0A024R5M9 | 1.80E-04 | 3.39 |
| ENSG00000213719 | CLIC1 | chloride intracellular channel 1 | O00299 | 8.40E-05 | 3.37 |
| ENSG00000131143 | COX4I1 | cytochrome c oxidase subunit 4I1 | P13073 | 5.60E-05 | 3.37 |
| ENSG00000129562 | DAD1 | defender against cell death 1 | P61803 | 2.20E-04 | 3.37 |
| ENSG00000168872 | DDX19A | DEAD-box helicase 19A | B4DS24 | 3.20E-04 | 3.37 |
| ENSG00000099901 | RANBP1 | RAN binding protein 1 | F6WQW2 | 4.70E-05 | 3.36 |
| ENSG00000135318 | NT5E | 5'-nucleotidase ecto | P21589 | 0.018 | 3.35 |
| ENSG00000127314 | RAP1B | RAP1B, member of RAS oncogene family | A0A024RB87 | 1.70E-04 | 3.35 |
| ENSG00000197894 | ADH5 | alcohol dehydrogenase 5 (class III), chi polypeptide | P11766 | 6.10E-05 | 3.34 |
| ENSG00000149781 | FERMT3 | FERM domain containing kindlin 3 | Q86UX7 | 1.20E-04 | 3.34 |
| ENSG00000144959 | NCEH1 | neutral cholesterol ester hydrolase 1 | A0A0A0MTJ9 | 6.50E-04 | 3.34 |
| ENSG00000100300 | TSPO | translocator protein | O76068 | 6.00E-05 | 3.34 |
| ENSG00000101361 | NOP56 | NOP56 ribonucleoprotein | O00567 | 6.40E-05 | 3.33 |
| ENSG00000164828 | SUN1 | Sad1 and UNC84 domain containing 1 | O94901 | 1.40E-04 | 3.32 |
| ENSG00000128245 | YWHAH | tyrosine 3-monooxygenase/tryptophan 5-monooxygenase activation protein eta | A0A024R1K7 | 1.20E-04 | 3.32 |
| ENSG00000104687 | GSR | glutathione-disulfide reductase | P00390 | 2.10E-04 | 3.31 |
| ENSG00000100664 | EIF5 | eukaryotic translation initiation factor 5 | A0A024R6Q1 | 0.003 | 3.3 |
| ENSG00000087470 | DNM1L | dynamin 1 like | B4DYR6 | 1.90E-04 | 3.29 |
| ENSG00000125691 | RPL23 | ribosomal protein L23 | A0A024R1Q8 | 0.00093 | 3.29 |
| ENSG00000087274 | ADD1 | adducin 1 | P35611 | 1.50E-04 | 3.28 |
| ENSG00000134748 | PRPF38A | pre-mRNA processing factor 38A | Q8NAV1 | 1.40E-04 | 3.28 |
| ENSG00000108848 | LUC7L3 | LUC7 like 3 pre-mRNA splicing factor | J3KPP4 | 4.80E-05 | 3.27 |
| ENSG00000151247 | EIF4E | eukaryotic translation initiation factor 4E | P06730 | 0.0033 | 3.26 |
| ENSG00000114480 | GBE1 | 1,4-alpha-glucan branching enzyme 1 | Q04446 | 1.20E-04 | 3.26 |
| ENSG00000141543 | EIF4A3 | eukaryotic translation initiation factor 4A3 | A0A024R8W0 | 1.60E-04 | 3.25 |
| ENSG00000170027 | YWHAG | tyrosine 3-monooxygenase/tryptophan 5-monooxygenase activation protein gamma | P61981 | 1.70E-04 | 3.25 |
| ENSG00000185825 | BCAP31 | B cell receptor associated protein 31 | P51572 | 1.30E-04 | 3.24 |
| ENSG00000169710 | FASN | fatty acid synthase | P49327 | 9.10E-05 | 3.24 |

| | | | | | |
|---|---|---|---|---|---|
| ENSG00000170348 | TMED10 | transmembrane p24 trafficking protein 10 | A0A024R6I3 | 2.90E-04 | 3.24 |
| ENSG00000105568 | PPP2R1A | protein phosphatase 2 scaffold subunit Aalpha | A8K7B7 | 8.50E-05 | 3.23 |
| ENSG00000010256 | UQCRC1 | ubiquinol-cytochrome c reductase core protein 1 | P31930 | 6.20E-04 | 3.23 |
| ENSG00000105974 | CAV1 | caveolin 1 | A0A024R757 | 4.80E-05 | 3.22 |
| ENSG00000100519 | PSMC6 | proteasome 26S subunit, ATPase 6 | A0A087X2I1 | 4.70E-05 | 3.22 |
| ENSG00000119707 | RBM25 | RNA binding motif protein 25 | P49756 | 1.70E-04 | 3.22 |
| ENSG00000100075 | SLC25A1 | solute carrier family 25 member 1 | D9HTE9 | 6.60E-05 | 3.22 |
| ENSG00000079785 | DDX1 | DEAD-box helicase 1 | A3RJH1 | 8.90E-05 | 3.21 |
| ENSG00000125868 | DSTN | destrin, actin depolymerizing factor | P60981 | 6.50E-04 | 3.21 |
| ENSG00000100097 | LGALS1 | galectin 1 | P09382 | 4.50E-04 | 3.2 |
| ENSG00000126561 | STAT5A | signal transducer and activator of transcription 5A | A8K6I5 | 0.013 | 3.2 |
| ENSG00000265808 | SEC22B | SEC22 homolog B, vesicle trafficking protein | O75396 | 1.70E-04 | 3.19 |
| ENSG00000178035 | IMPDH2 | inosine monophosphate dehydrogenase 2 | P12268 | 5.80E-04 | 3.18 |
| ENSG00000095139 | ARCN1 | archain 1 | P48444 | 1.20E-04 | 3.17 |
| ENSG00000146731 | CCT6A | chaperonin containing TCP1 subunit 6A | P40227 | 2.60E-04 | 3.17 |
| ENSG00000122545 | SEPTIN7 | septin 7 | A8K3D0 | 1.80E-04 | 3.17 |
| ENSG00000135316 | SYNCRIP | synaptotagmin binding cytoplasmic RNA interacting protein | B7Z645 | 3.90E-04 | 3.17 |
| ENSG00000100934 | SEC23A | SEC23 homolog A, COPII coat complex component | Q15436 | 0.0012 | 3.16 |
| ENSG00000176014 | TUBB6 | tubulin beta 6 class V | Q9BUF5 | 4.30E-04 | 3.15 |
| ENSG00000055609 | KMT2C | lysine methyltransferase 2C | Q8NEZ4 | 0.0011 | 3.14 |
| ENSG00000178921 | PFAS | phosphoribosylformylglycinamidine synthase | A8K9T9 | 2.10E-04 | 3.14 |
| ENSG00000109084 | TMEM97 | transmembrane protein 97 | Q5BJF2 | 4.10E-04 | 3.14 |
| ENSG00000134186 | PRPF38B | pre-mRNA processing factor 38B | Q5VTL8 | 6.20E-05 | 3.13 |
| ENSG00000117592 | PRDX6 | peroxiredoxin 6 | P30041 | 2.60E-04 | 3.12 |
| ENSG00000091409 | ITGA6 | integrin subunit alpha 6 | P23229 | 2.80E-04 | 3.11 |
| ENSG00000166181 | API5 | apoptosis inhibitor 5 | Q9BZZ5 | 5.10E-04 | 3.1 |
| ENSG00000211893 | IGHG2 | immunoglobulin heavy constant gamma 2 (G2m marker) | NA | 0.0021 | 3.08 |
| ENSG00000111581 | NUP107 | nucleoporin 107 | P57740 | 0.0037 | 3.08 |
| ENSG00000175582 | RAB6A | RAB6A, member RAS oncogene family | P20340 | 0.0011 | 3.08 |
| ENSG00000163636 | PSMD6 | proteasome 26S subunit, non-ATPase 6 | Q15008 | 3.10E-04 | 3.07 |
| ENSG00000116754 | SRSF11 | serine and arginine rich splicing factor 11 | Q05519 | 1.10E-04 | 3.07 |
| ENSG00000179218 | CALR | calreticulin | P27797 | 1.90E-04 | 3.06 |
| ENSG00000100030 | MAPK1 | mitogen-activated protein kinase 1 | P28482 | 1.30E-04 | 3.06 |
| ENSG00000100813 | ACIN1 | apoptotic chromatin condensation inducer 1 | Q9UKV3 | 9.50E-05 | 3.05 |
| ENSG00000146376 | ARHGAP18 | Rho GTPase activating protein 18 | Q8N392 | 4.40E-04 | 3.05 |
| ENSG00000125944 | HNRNPR | heterogeneous nuclear ribonucleoprotein R | Q0VGD6 | 3.10E-04 | 3.04 |
| ENSG00000167553 | TUBA1C | tubulin alpha 1c | B7Z1K5 | 0.022 | 3.01 |

| | | | | | |
|---|---|---|---|---|---|
| ENSG00000130309 | COLGALT1 | collagen beta(1-O)galactosyltransferase 1 | Q8NBJ5 | 2.40E-04 | 2.99 |
| ENSG00000062485 | CS | citrate synthase | A0A024RB75 | 0.0014 | 2.98 |
| ENSG00000143815 | LBR | lamin B receptor | Q14739 | 2.30E-04 | 2.98 |
| ENSG00000109501 | WFS1 | wolframin ER transmembrane glycoprotein | A0A0S2Z4V6 | 1.00E-04 | 2.98 |
| ENSG00000102898 | NUTF2 | nuclear transport factor 2 | A0A024R6Y2 | 0.0012 | 2.97 |
| ENSG00000075624 | ACTB | actin beta | P60709 | 3.30E-04 | 2.95 |
| ENSG00000134440 | NARS1 | asparaginyl-tRNA synthetase 1 | O43776 | 1.70E-04 | 2.95 |
| ENSG00000104325 | DECR1 | 2,4-dienoyl-CoA reductase 1 | Q16698 | 4.70E-04 | 2.94 |
| ENSG00000169919 | GUSB | glucuronidase beta | P08236 | 1.30E-04 | 2.93 |
| ENSG00000169504 | CLIC4 | chloride intracellular channel 4 | Q6FIC5 | 1.90E-04 | 2.92 |
| ENSG00000172725 | CORO1B | coronin 1B | A0A024R5K1 | 1.20E-04 | 2.92 |
| ENSG00000070756 | PABPC1 | poly(A) binding protein cytoplasmic 1 | A0A024R9C1 | 3.80E-04 | 2.92 |
| ENSG00000130313 | PGLS | 6-phosphogluconolactonase | A0A0K0K1K7 | 8.80E-05 | 2.92 |
| ENSG00000108946 | PRKAR1A | protein kinase cAMP-dependent type I regulatory subunit alpha | B2R5T5 | 0.0031 | 2.91 |
| ENSG00000132341 | RAN | RAN, member RAS oncogene family | B4DV51 | 0.0012 | 2.91 |
| ENSG00000100504 | PYGL | glycogen phosphorylase L | P06737 | 0.0037 | 2.9 |
| ENSG00000022267 | FHL1 | four and a half LIM domains 1 | Q13642 | 2.20E-04 | 2.89 |
| ENSG00000176658 | MYO1D | myosin ID | J3QRN6 | 2.20E-04 | 2.89 |
| ENSG00000213639 | PPP1CB | protein phosphatase 1 catalytic subunit beta | P62140 | 2.90E-04 | 2.89 |
| ENSG00000092201 | SUPT16H | SPT16 homolog, facilitates chromatin remodeling subunit | Q9Y5B9 | 1.80E-04 | 2.88 |
| ENSG00000169067 | ACTBL2 | actin beta like 2 | Q562R1 | 3.90E-04 | 2.87 |
| ENSG00000074696 | HACD3 | 3-hydroxyacyl-CoA dehydratase 3 | Q9P035 | 2.30E-04 | 2.87 |
| ENSG00000106397 | PLOD3 | procollagen-lysine,2-oxoglutarate 5-dioxygenase 3 | O60568 | 3.10E-04 | 2.87 |
| ENSG00000198363 | ASPH | aspartate beta-hydroxylase | Q12797 | 3.10E-04 | 2.86 |
| ENSG00000133318 | RTN3 | reticulon 3 | O95197 | 1.90E-04 | 2.86 |
| ENSG00000171560 | FGA | fibrinogen alpha chain | P02671 | 0.013 | 2.84 |
| ENSG00000132507 | EIF5A | eukaryotic translation initiation factor 5A | P63241 | 2.70E-04 | 2.83 |
| ENSG00000169738 | DCXR | dicarbonyl and L-xylulose reductase | Q7Z4W1 | 4.70E-04 | 2.81 |
| ENSG00000197111 | PCBP2 | poly(rC) binding protein 2 | Q15366 | 3.40E-04 | 2.81 |
| ENSG00000101558 | VAPA | VAMP associated protein A | Q9P0L0 | 3.80E-04 | 2.81 |
| ENSG00000130429 | ARPC1B | actin related protein 2/3 complex subunit 1B | A4D275 | 2.70E-04 | 2.8 |
| ENSG00000198668 | CALM1 | calmodulin 1 | B4DJ51 | 6.00E-04 | 2.8 |
| ENSG00000075856 | SART3 | spliceosome associated factor 3, U4/U6 recycling protein | Q15020 | 0.0099 | 2.79 |
| ENSG00000138772 | ANXA3 | annexin A3 | P12429 | 1.50E-04 | 2.77 |
| ENSG00000150593 | PDCD4 | programmed cell death 4 | B4DKX4 | 4.50E-04 | 2.77 |
| ENSG00000239264 | TXNDC5 | thioredoxin domain containing 5 | A0A024QZV0 | 2.90E-04 | 2.77 |
| ENSG00000072778 | ACADVL | acyl-CoA dehydrogenase very long chain | P49748 | 4.20E-04 | 2.76 |
| ENSG00000154473 | BUB3 | BUB3 mitotic checkpoint protein | A0A140VJF3 | 0.0011 | 2.75 |
| ENSG00000143621 | ILF2 | interleukin enhancer binding factor 2 | B4DY09 | 4.10E-04 | 2.74 |
| ENSG00000103642 | LACTB | lactamase beta | P83111 | 0.0022 | 2.74 |

| | | | | | |
|---|---|---|---|---|---|
| ENSG00000167004 | PDIA3 | protein disulfide isomerase family A member 3 | P30101 | 4.20E-04 | 2.74 |
| ENSG00000157020 | SEC13 | SEC13 homolog, nuclear pore and COPII coat complex component | P55735 | 5.90E-04 | 2.74 |
| ENSG00000090273 | NUDC | nuclear distribution C, dynein complex regulator | Q9Y266 | 0.0013 | 2.73 |
| ENSG00000197702 | PARVA | parvin alpha | J3KNQ4 | 0.0021 | 2.73 |
| ENSG00000167699 | GLOD4 | glyoxalase domain containing 4 | Q9HC38 | 2.00E-04 | 2.72 |
| ENSG00000083845 | RPS5 | ribosomal protein S5 | A0A024R4Q8 | 5.30E-04 | 2.72 |
| ENSG00000149089 | APIP | APAF1 interacting protein | Q96GX9 | 0.0014 | 2.71 |
| ENSG00000158710 | TAGLN2 | transgelin 2 | P37802 | 0.0033 | 2.71 |
| ENSG00000122958 | VPS26A | VPS26 retromer complex component A | O75436 | 0.01 | 2.71 |
| ENSG00000169021 | UQCRFS1 | ubiquinol-cytochrome c reductase, Rieske iron-sulfur polypeptide 1 | P47985 | 0.0017 | 2.7 |
| ENSG00000130396 | AFDN | afadin, adherens junction formation factor | P55196 | 0.0022 | 2.69 |
| ENSG00000177731 | FLII | FLII actin remodeling protein | Q13045 | 4.60E-04 | 2.69 |
| ENSG00000120254 | MTHFD1L | methylenetetrahydrofolate dehydrogenase (NADP+ dependent) 1 like | B7ZM99 | 1.60E-04 | 2.69 |
| ENSG00000125977 | EIF2S2 | eukaryotic translation initiation factor 2 subunit beta | P20042 | 0.0011 | 2.68 |
| ENSG00000067113 | PLPP1 | phospholipid phosphatase 1 | A0A024QZS3 | 0.0028 | 2.68 |
| ENSG00000168003 | SLC3A2 | solute carrier family 3 member 2 | J3KPF3 | 2.90E-04 | 2.68 |
| ENSG00000166747 | AP1G1 | adaptor related protein complex 1 subunit gamma 1 | A0A140VJE7 | 3.70E-04 | 2.65 |
| ENSG00000143819 | EPHX1 | epoxide hydrolase 1 | P07099 | 9.70E-04 | 2.65 |
| ENSG00000105379 | ETFB | electron transfer flavoprotein subunit beta | P38117 | 6.10E-04 | 2.65 |
| ENSG00000119396 | RAB14 | RAB14, member RAS oncogene family | A0A024R845 | 0.0071 | 2.65 |
| ENSG00000140740 | UQCRC2 | ubiquinol-cytochrome c reductase core protein 2 | P22695 | 0.0031 | 2.65 |
| ENSG00000151693 | ASAP2 | ArfGAP with SH3 domain, ankyrin repeat and PH domain 2 | O43150 | 5.30E-04 | 2.64 |
| ENSG00000108094 | CUL2 | cullin 2 | A0A140VKB1 | 0.035 | 2.64 |
| ENSG00000154174 | TOMM70 | translocase of outer mitochondrial membrane 70 | O94826 | 0.0014 | 2.64 |
| ENSG00000091164 | TXNL1 | thioredoxin like 1 | O43396 | 0.0012 | 2.64 |
| ENSG00000164022 | AIMP1 | aminoacyl tRNA synthetase complex interacting multifunctional protein 1 | B4DNK3 | 5.60E-04 | 2.63 |
| ENSG00000148841 | ITPRIP | inositol 1,4,5-trisphosphate receptor interacting protein | Q8IWB1 | 3.20E-04 | 2.63 |
| ENSG00000099246 | RAB18 | RAB18, member RAS oncogene family | Q9NP72 | 6.00E-04 | 2.63 |
| ENSG00000132842 | AP3B1 | adaptor related protein complex 3 subunit beta 1 | A0A0S2Z5J4 | 4.20E-04 | 2.6 |
| ENSG00000138777 | PPA2 | inorganic pyrophosphatase 2 | Q9H2U2 | 0.0024 | 2.6 |
| ENSG00000084652 | TXLNA | taxilin alpha | P40222 | 3.70E-04 | 2.6 |
| ENSG00000145833 | DDX46 | DEAD-box helicase 46 | A0A0C4DG89 | 9.00E-04 | 2.59 |
| ENSG00000118257 | NRP2 | neuropilin 2 | O60462 | 3.20E-04 | 2.59 |
| ENSG00000100911 | PSME2 | proteasome activator subunit 2 | Q86SZ7 | 9.20E-04 | 2.59 |
| ENSG00000169398 | PTK2 | protein tyrosine kinase 2 | Q05397 | 3.30E-04 | 2.59 |
| ENSG00000137710 | RDX | radixin | P35241 | 2.40E-04 | 2.58 |

| | | | | | |
|---|---|---|---|---|---|
| ENSG00000073969 | NSF | N-ethylmaleimide sensitive factor, vesicle fusing ATPase | P46459 | 2.70E-04 | 2.57 |
| ENSG00000089157 | RPLP0 | ribosomal protein lateral stalk subunit P0 | A0A024RBS2 | 3.40E-04 | 2.57 |
| ENSG00000147140 | NONO | non-POU domain containing octamer binding | A0A0S2Z4Z9 | 0.0043 | 2.55 |
| ENSG00000031698 | SARS1 | seryl-tRNA synthetase 1 | Q5T5C7 | 8.10E-04 | 2.54 |
| ENSG00000170876 | TMEM43 | transmembrane protein 43 | A0A024R2F9 | 0.0023 | 2.54 |
| ENSG00000107862 | GBF1 | golgi brefeldin A resistant guanine nucleotide exchange factor 1 | Q92538 | 4.50E-04 | 2.53 |
| ENSG00000067560 | RHOA | ras homolog family member A | A0A024R324 | 0.011 | 2.53 |
| ENSG00000163191 | S100A11 | S100 calcium binding protein A11 | P31949 | 0.0022 | 2.53 |
| ENSG00000115233 | PSMD14 | proteasome 26S subunit, non-ATPase 14 | A0A140VKF2 | 0.003 | 2.52 |
| ENSG00000136450 | SRSF1 | serine and arginine rich splicing factor 1 | Q07955 | 0.0082 | 2.52 |
| ENSG00000188229 | TUBB4B | tubulin beta 4B class IVb | P68371 | 9.50E-04 | 2.52 |
| ENSG00000134308 | YWHAQ | tyrosine 3-monooxygenase/tryptophan 5-monooxygenase activation protein theta | P27348 | 6.60E-04 | 2.52 |
| ENSG00000148700 | ADD3 | adducin 3 | Q9UEY8 | 3.80E-04 | 2.5 |
| ENSG00000168374 | ARF4 | ADP ribosylation factor 4 | P18085 | 0.0018 | 2.5 |
| ENSG00000125107 | CNOT1 | CCR4-NOT transcription complex subunit 1 | A5YKK6 | 0.0065 | 2.5 |
| ENSG00000197728 | RPS26 | ribosomal protein S26 | A0A024RB14 | 0.013 | 2.5 |
| ENSG00000165092 | ALDH1A1 | aldehyde dehydrogenase 1 family member A1 | P00352 | 7.80E-04 | 2.49 |
| ENSG00000065485 | PDIA5 | protein disulfide isomerase family A member 5 | Q14554 | 0.0011 | 2.49 |
| ENSG00000187514 | PTMA | prothymosin alpha | P06454 | 0.015 | 2.49 |
| ENSG00000177105 | RHOG | ras homolog family member G | P84095 | 0.0016 | 2.47 |
| ENSG00000162695 | SLC30A7 | solute carrier family 30 member 7 | Q8NEW0 | 8.60E-04 | 2.47 |
| ENSG00000142192 | APP | amyloid beta precursor protein | A0A140VJC8 | 0.0035 | 2.46 |
| ENSG00000162616 | DNAJB4 | DnaJ heat shock protein family (Hsp40) member B4 | Q9UDY4 | 6.20E-04 | 2.46 |
| ENSG00000197892 | KIF13B | kinesin family member 13B | Q9NQT8 | 0.0038 | 2.46 |
| ENSG00000168259 | DNAJC7 | DnaJ heat shock protein family (Hsp40) member C7 | Q99615 | 0.0011 | 2.45 |
| ENSG00000176619 | LMNB2 | lamin B2 | Q03252 | 6.80E-04 | 2.45 |
| ENSG00000130726 | TRIM28 | tripartite motif containing 28 | Q13263 | 0.0035 | 2.45 |
| ENSG00000113810 | SMC4 | structural maintenance of chromosomes 4 | Q58F29 | 8.30E-04 | 2.44 |
| ENSG00000163931 | TKT | transketolase | P29401 | 4.40E-04 | 2.44 |
| ENSG00000101150 | TPD52L2 | TPD52 like 2 | O43399 | 4.80E-04 | 2.44 |
| ENSG00000100243 | CYB5R3 | cytochrome b5 reductase 3 | P00387 | 6.50E-04 | 2.42 |
| ENSG00000137509 | PRCP | prolylcarboxypeptidase | B7Z7Q6 | 0.008 | 2.42 |
| ENSG00000054118 | THRAP3 | thyroid hormone receptor associated protein 3 | Q9Y2W1 | 0.013 | 2.42 |
| ENSG00000140350 | ANP32A | acidic nuclear phosphoprotein 32 family member A | P39687 | 0.0012 | 2.41 |
| ENSG00000166825 | ANPEP | alanyl aminopeptidase, membrane | A0A024RC61 | 0.0042 | 2.41 |
| ENSG00000057757 | PITHD1 | PITH domain containing 1 | Q9GZP4 | 6.40E-04 | 2.41 |
| ENSG00000101474 | APMAP | adipocyte plasma membrane associated protein | Q9HDC9 | 0.0022 | 2.39 |

| | | | | | |
|---|---|---|---|---|---|
| ENSG00000273703 | H2BC14 | H2B clustered histone 14 | Q99879 | 0.044 | 2.39 |
| ENSG00000141959 | PFKL | phosphofructokinase, liver type | P17858 | 9.40E-04 | 2.39 |
| ENSG00000133226 | SRRM1 | serine and arginine repetitive matrix 1 | B7Z7U0 | 6.10E-04 | 2.39 |
| ENSG00000085063 | CD59 | CD59 molecule (CD59 blood group) | P13987 | 0.026 | 2.38 |
| ENSG00000166333 | ILK | integrin linked kinase | Q13418 | 0.0045 | 2.38 |
| ENSG00000112245 | PTP4A1 | protein tyrosine phosphatase 4A1 | A0A024R8J2 | 0.019 | 2.38 |
| ENSG00000075415 | SLC25A3 | solute carrier family 25 member 3 | A0A024RBE8 | 5.80E-04 | 2.38 |
| ENSG00000142937 | RPS8 | ribosomal protein S8 | P62241 | 8.40E-04 | 2.37 |
| ENSG00000143549 | TPM3 | tropomyosin 3 | A0A0S2Z4I4 | 5.90E-04 | 2.37 |
| ENSG00000102401 | ARMCX3 | armadillo repeat containing X-linked 3 | A0A024RCF9 | 7.30E-04 | 2.36 |
| ENSG00000241553 | ARPC4 | actin related protein 2/3 complex subunit 4 | P59998 | 0.0024 | 2.36 |
| ENSG00000133112 | TPT1 | tumor protein, translationally-controlled 1 | A0A0B4J2C3 | 0.026 | 2.36 |
| ENSG00000153904 | DDAH1 | dimethylarginine dimethylaminohydrolase 1 | B4E3V1 | 0.0011 | 2.35 |
| ENSG00000104408 | EIF3E | eukaryotic translation initiation factor 3 subunit E | P60228 | 5.20E-04 | 2.35 |
| ENSG00000169756 | LIMS1 | LIM zinc finger domain containing 1 | P48059 | 6.90E-04 | 2.35 |
| ENSG00000147274 | RBMX | RNA binding motif protein X-linked | P38159 | 0.0013 | 2.35 |
| ENSG00000197958 | RPL12 | ribosomal protein L12 | P30050 | 4.30E-04 | 2.35 |
| ENSG00000079332 | SAR1A | secretion associated Ras related GTPase 1A | Q5SQT9 | 0.0029 | 2.35 |
| ENSG00000028528 | SNX1 | sorting nexin 1 | Q13596 | 8.30E-04 | 2.35 |
| ENSG00000115380 | EFEMP1 | EGF containing fibulin extracellular matrix protein 1 | A0A0S2Z4F1 | 0.0015 | 2.34 |
| ENSG00000132780 | NASP | nuclear autoantigenic sperm protein | P49321 | 9.10E-04 | 2.34 |
| ENSG00000167792 | NDUFV1 | NADH:ubiquinone oxidoreductase core subunit V1 | P49821 | 9.70E-04 | 2.34 |
| ENSG00000075618 | FSCN1 | fascin actin-bundling protein 1 | B3KTA3 | 0.002 | 2.33 |
| ENSG00000103342 | GSPT1 | G1 to S phase transition 1 | P15170 | 5.40E-04 | 2.32 |
| ENSG00000092841 | MYL6 | myosin light chain 6 | P60660 | 0.0016 | 2.32 |
| ENSG00000023228 | NDUFS1 | NADH:ubiquinone oxidoreductase core subunit S1 | P28331 | 0.0035 | 2.32 |
| ENSG00000076043 | REXO2 | RNA exonuclease 2 | Q9Y3B8 | 0.0011 | 2.32 |
| ENSG00000072501 | SMC1A | structural maintenance of chromosomes 1A | G8JLG1 | 0.0014 | 2.32 |
| ENSG00000148834 | GSTO1 | glutathione S-transferase omega 1 | P78417 | 7.10E-04 | 2.31 |
| ENSG00000171863 | RPS7 | ribosomal protein S7 | P62081 | 0.0021 | 2.31 |
| ENSG00000127022 | CANX | calnexin | P27824 | 5.60E-04 | 2.3 |
| ENSG00000139684 | ESD | esterase D | A0A140VJJ2 | 0.0026 | 2.3 |
| ENSG00000167770 | OTUB1 | OTU deubiquitinase, ubiquitin aldehyde binding 1 | B3KUV5 | 5.70E-04 | 2.3 |
| ENSG00000166340 | TPP1 | tripeptidyl peptidase 1 | O14773 | 0.0046 | 2.3 |
| ENSG00000185624 | P4HB | prolyl 4-hydroxylase subunit beta | A0A024R8S5 | 0.004 | 2.29 |
| ENSG00000241973 | PI4KA | phosphatidylinositol 4-kinase alpha | B4DYG5 | 0.0077 | 2.29 |

Adjustments of p-values for multiple comparisons were used with Benjamini-Hochberg (BH) correction.

### Supplemental Table S5. Palmitoylated proteins in insulin treated siCD36 LECs.

| ENSEMBL ID | Gene Symbol | Description | UNIPROT | *p*-value | -Log Ratio (+/- HA) |
|---|---|---|---|---|---|
| ENSG00000213699 | SLC35F6 | solute carrier family 35 member F6 | Q8N357 | 0.00000017 | 7.76 |
| ENSG00000002586 | CD99 | CD99 molecule (Xg blood group) | P14209 | 0.00000014 | 7.02 |
| ENSG00000054116 | TRAPPC3 | trafficking protein particle complex subunit 3 | A0A087WWM0 | 0.00000041 | 6.82 |
| ENSG00000111897 | SERINC1 | serine incorporator 1 | Q9NRX5 | 0.00000036 | 6.71 |
| ENSG00000102007 | PLP2 | proteolipid protein 2 | A0A024QYW3 | 0.0000021 | 6.67 |
| ENSG00000136156 | ITM2B | integral membrane protein 2B | Q9Y287 | 0.00000029 | 6.63 |
| ENSG00000110651 | CD81 | CD81 molecule | E9PJK1 | 0.00000032 | 6.49 |
| ENSG00000177697 | CD151 | CD151 molecule (Raph blood group) | A0A024RCB3 | 0.0000012 | 6.41 |
| ENSG00000076706 | MCAM | melanoma cell adhesion molecule | A0A024R3I5 | 0.00000056 | 6.27 |
| ENSG00000143878 | RHOB | ras homolog family member B | P62745 | 0.0000016 | 6.24 |
| ENSG00000139921 | TMX1 | thioredoxin related transmembrane protein 1 | Q9H3N1 | 0.00000047 | 6.08 |
| ENSG00000132388 | UBE2G1 | ubiquitin conjugating enzyme E2 G1 | P62253 | 0.00000052 | 6.03 |
| ENSG00000049245 | VAMP3 | vesicle associated membrane protein 3 | Q15836 | 0.00000085 | 6.02 |
| ENSG00000114698 | PLSCR4 | phospholipid scramblase 4 | Q9NRQ2 | 0.00000099 | 6.02 |
| ENSG00000135404 | CD63 | CD63 molecule | A0A024RB05 | 0.0000058 | 6.01 |
| ENSG00000092531 | SNAP23 | synaptosome associated protein 23 | A8K287 | 0.00000044 | 5.99 |
| ENSG00000003056 | M6PR | mannose-6-phosphate receptor, cation dependent | F5GX30 | 0.00000081 | 5.96 |
| ENSG00000133818 | RRAS2 | RAS related 2 | P62070 | 0.0000011 | 5.96 |
| ENSG00000185651 | UBE2L3 | ubiquitin conjugating enzyme E2 L3 | P68036 | 0.0000049 | 5.9 |
| ENSG00000144744 | UBA3 | ubiquitin like modifier activating enzyme 3 | Q8TBC4 | 0.00000072 | 5.86 |
| ENSG00000110917 | MLEC | malectin | F5GX14 | 0.00000084 | 5.76 |
| ENSG00000187838 | PLSCR3 | phospholipid scramblase 3 | Q9NRY6 | 0.0000024 | 5.76 |
| ENSG00000143222 | UFC1 | ubiquitin-fold modifier conjugating enzyme 1 | Q9Y3C8 | 0.000012 | 5.74 |
| ENSG00000072778 | ACADVL | acyl-CoA dehydrogenase very long chain | P49748 | 0.0000019 | 5.71 |
| ENSG00000184113 | CLDN5 | claudin 5 | D3DX19 | 0.0000034 | 5.67 |
| ENSG00000169908 | TM4SF1 | transmembrane 4 L six family member 1 | P30408 | 0.00000083 | 5.59 |
| ENSG00000130725 | UBE2M | ubiquitin conjugating enzyme E2 M | A0A024R4T4 | 0.0000027 | 5.59 |
| ENSG00000148175 | STOM | stomatin | F8VSL7 | 0.0000016 | 5.58 |
| ENSG00000129625 | REEP5 | receptor accessory protein 5 | Q00765 | 0.0000016 | 5.54 |
| ENSG00000227500 | SCAMP4 | secretory carrier membrane protein 4 | Q969E2 | 0.0000014 | 5.44 |
| ENSG00000033178 | UBA6 | ubiquitin like modifier activating enzyme 6 | A0A024RDB0 | 0.0000013 | 5.38 |

| | | | | | |
|---|---|---|---|---|---|
| ENSG00000005194 | CIAPIN1 | cytokine induced apoptosis inhibitor 1 | Q6FI81 | 0.0000049 | 5.37 |
| ENSG00000141543 | EIF4A3 | eukaryotic translation initiation factor 4A3 | A0A024R8W0 | 0.0000039 | 5.35 |
| ENSG00000142089 | IFITM3 | interferon induced transmembrane protein 3 | Q01628 | 0.0000026 | 5.33 |
| ENSG00000143079 | CTTNBP2NL | CTTNBP2 N-terminal like | A0A024R0C7 | 0.000024 | 5.32 |
| ENSG00000108219 | TSPAN14 | tetraspanin 14 | Q8NG11 | 0.0000044 | 5.31 |
| ENSG00000168899 | VAMP5 | vesicle associated membrane protein 5 | O95183 | 0.0000043 | 5.28 |
| ENSG00000116521 | SCAMP3 | secretory carrier membrane protein 3 | O14828 | 0.0000023 | 5.21 |
| ENSG00000183726 | TMEM50A | transmembrane protein 50A | O95807 | 0.0000051 | 5.11 |
| ENSG00000168374 | ARF4 | ADP ribosylation factor 4 | P18085 | 0.000012 | 5.11 |
| ENSG00000169756 | LIMS1 | LIM zinc finger domain containing 1 | P48059 | 0.0000024 | 5.1 |
| ENSG00000129353 | SLC44A2 | solute carrier family 44 member 2 | A0A088QCU6 | 0.000015 | 5.08 |
| ENSG00000132589 | FLOT2 | flotillin 2 | J3QLD9 | 0.0000097 | 5.07 |
| ENSG00000204525 | HLA-C | major histocompatibility complex, class I, C | O19617 | 0.000024 | 5.05 |
| ENSG00000037280 | FLT4 | fms related receptor tyrosine kinase 4 | P35916 | 0.0000051 | 5.02 |
| ENSG00000115310 | RTN4 | reticulon 4 | Q9NQC3 | 0.000011 | 4.93 |
| ENSG00000243279 | PRAF2 | PRA1 domain family member 2 | A0A024QZ22 | 0.00001 | 4.91 |
| ENSG00000010278 | CD9 | CD9 molecule | A6NNI4 | 0.0000025 | 4.9 |
| ENSG00000204632 | HLA-G | major histocompatibility complex, class I, G | P17693 | 0.0000084 | 4.88 |
| ENSG00000249751 | ECSCR | endothelial cell surface expressed chemotaxis and apoptosis regulator | Q19T08 | 0.0000087 | 4.88 |
| ENSG00000113811 | SELENOK | selenoprotein K | Q9Y6D0 | 0.0000067 | 4.87 |
| ENSG00000141458 | NPC1 | NPC intracellular cholesterol transporter 1 | O15118 | 0.000016 | 4.86 |
| ENSG00000099282 | TSPAN15 | tetraspanin 15 | O95858 | 0.000027 | 4.86 |
| ENSG00000134247 | PTGFRN | prostaglandin F2 receptor inhibitor | Q9P2B2 | 0.00001 | 4.83 |
| ENSG00000006451 | RALA | RAS like proto-oncogene A | P11233 | 0.0000079 | 4.82 |
| ENSG00000078140 | UBE2K | ubiquitin conjugating enzyme E2 K | P61086 | 0.0000045 | 4.79 |
| ENSG00000105379 | ETFB | electron transfer flavoprotein subunit beta | P38117 | 0.0000096 | 4.72 |
| ENSG00000168515 | SCGB1D1 | secretoglobin family 1D member 1 | O95968 | 0.0019 | 4.69 |
| ENSG00000099864 | PALM | paralemmin | A0A024R207 | 0.00001 | 4.66 |
| ENSG00000114115 | RBP1 | retinol binding protein 1 | P09455 | 0.0000054 | 4.65 |
| ENSG00000188313 | PLSCR1 | phospholipid scramblase 1 | O15162 | 0.000024 | 4.65 |
| ENSG00000122378 | PRXL2A | peroxiredoxin like 2A | Q9BRX8 | 0.0000069 | 4.62 |
| ENSG00000177889 | UBE2N | ubiquitin conjugating enzyme E2 N | P61088 | 0.0000034 | 4.6 |
| ENSG00000087460 | GNAS | GNAS complex locus | A0A0S2Z3H8 | 0.00002 | 4.59 |
| ENSG00000143815 | LBR | lamin B receptor | Q14739 | 0.00001 | 4.56 |
| ENSG00000133318 | RTN3 | reticulon 3 | O95197 | 0.0000063 | 4.54 |

| | | | | | |
|---|---|---|---|---|---|
| ENSG00000163762 | TM4SF18 | transmembrane 4 L six family member 18 | Q96CE8 | 0.000034 | 4.49 |
| ENSG00000114353 | GNAI2 | G protein subunit alpha i2 | B3KP24 | 0.0000097 | 4.43 |
| ENSG00000178035 | IMPDH2 | inosine monophosphate dehydrogenase 2 | P12268 | 0.000058 | 4.42 |
| ENSG00000118873 | RAB3GAP2 | RAB3 GTPase activating non-catalytic protein subunit 2 | Q9H2M9 | 0.0000069 | 4.39 |
| ENSG00000233276 | GPX1 | glutathione peroxidase 1 | P07203 | 0.00001 | 4.38 |
| ENSG00000144118 | RALB | RAS like proto-oncogene B | A0A024RAG3 | 0.0000088 | 4.36 |
| ENSG00000135047 | CTSL | cathepsin L | A0A024R276 | 0.0000078 | 4.34 |
| ENSG00000144848 | ATG3 | autophagy related 3 | Q9NT62 | 0.000011 | 4.27 |
| ENSG00000213281 | NRAS | NRAS proto-oncogene, GTPase | P01111 | 0.000051 | 4.16 |
| ENSG00000168172 | HOOK3 | hook microtubule tethering protein 3 | Q86VS8 | 0.00085 | 4.16 |
| ENSG00000198431 | TXNRD1 | thioredoxin reductase 1 | Q16881 | 0.000017 | 4.15 |
| ENSG00000141526 | SLC16A3 | solute carrier family 16 member 3 | A0A024R8U1 | 0.000011 | 4.12 |
| ENSG00000167996 | FTH1 | ferritin heavy chain 1 | A0A024R525 | 0.00004 | 4.1 |
| ENSG00000129562 | DAD1 | defender against cell death 1 | P61803 | 0.000055 | 4.09 |
| ENSG00000120437 | ACAT2 | acetyl-CoA acetyltransferase 2 | Q9BWD1 | 0.000048 | 4.03 |
| ENSG00000101160 | CTSZ | cathepsin Z | Q9UBR2 | 0.000051 | 4.01 |
| ENSG00000182809 | CRIP2 | cysteine rich protein 2 | P52943 | 0.000091 | 4 |
| ENSG00000197956 | S100A6 | S100 calcium binding protein A6 | P06703 | 0.0013 | 3.97 |
| ENSG00000109133 | TMEM33 | transmembrane protein 33 | A0A024R9W7 | 0.000043 | 3.95 |
| ENSG00000117298 | ECE1 | endothelin converting enzyme 1 | A0A024RAB0 | 0.00012 | 3.83 |
| ENSG00000072274 | TFRC | transferrin receptor | P02786 | 0.00014 | 3.83 |
| ENSG00000152661 | GJA1 | gap junction protein alpha 1 | P17302 | 0.00014 | 3.81 |
| ENSG00000052749 | RRP12 | ribosomal RNA processing 12 homolog | B3KMR5 | 0.000047 | 3.77 |
| ENSG00000197747 | S100A10 | S100 calcium binding protein A10 | P60903 | 0.00012 | 3.75 |
| ENSG00000066322 | ELOVL1 | ELOVL fatty acid elongase 1 | Q9BW60 | 0.000046 | 3.73 |
| ENSG00000107796 | ACTA2 | actin alpha 2, smooth muscle | D2JYH4 | 0.000087 | 3.71 |
| ENSG00000078668 | VDAC3 | voltage dependent anion channel 3 | Q9Y277 | 0.000024 | 3.7 |
| ENSG00000164022 | AIMP1 | aminoacyl tRNA synthetase complex interacting multifunctional protein 1 | B4DNK3 | 0.000051 | 3.69 |
| ENSG00000126214 | KLC1 | kinesin light chain 1 | Q07866 | 0.000023 | 3.68 |
| ENSG00000138760 | SCARB2 | scavenger receptor class B member 2 | Q14108 | 0.000026 | 3.68 |
| ENSG00000104131 | EIF3J | eukaryotic translation initiation factor 3 subunit J | O75822 | 0.000059 | 3.68 |
| ENSG00000125868 | DSTN | destrin, actin depolymerizing factor | P60981 | 0.00026 | 3.68 |
| ENSG00000160285 | LSS | lanosterol synthase | B2R694 | 0.000021 | 3.63 |
| ENSG00000166747 | AP1G1 | adaptor related protein complex 1 subunit gamma 1 | A0A140VJE7 | 0.00004 | 3.62 |
| ENSG00000184575 | XPOT | exportin for tRNA | O43592 | 0.000033 | 3.6 |
| ENSG00000114480 | GBE1 | 1,4-alpha-glucan branching enzyme 1 | Q04446 | 0.000059 | 3.6 |

| | | | | | |
|---|---|---|---|---|---|
| ENSG00000115419 | GLS | glutaminase | O94925 | 0.00014 | 3.6 |
| ENSG00000101367 | MAPRE1 | microtubule associated protein RP/EB family member 1 | Q15691 | 0.00003 | 3.58 |
| ENSG00000136026 | CKAP4 | cytoskeleton associated protein 4 | A0A024RBH2 | 0.000074 | 3.58 |
| NA | 5MP2_HUMAN | NA | NA | 0.00081 | 3.56 |
| ENSG00000135269 | TES | testin LIM domain protein | A4D0U5 | 0.000038 | 3.54 |
| ENSG00000262246 | CORO7 | coronin 7 | P57737 | 0.00018 | 3.54 |
| ENSG00000151914 | DST | dystonin | B4DSS9 | 0.0021 | 3.52 |
| ENSG00000112531 | QKI | QKI, KH domain containing RNA binding | Q96PU8 | 0.000027 | 3.46 |
| ENSG00000197249 | SERPINA1 | serpin family A member 1 | E9KL23 | 0.0064 | 3.45 |
| ENSG00000163565 | IFI16 | interferon gamma inducible protein 16 | Q16666 | 0.00024 | 3.43 |
| ENSG00000179091 | CYC1 | cytochrome c1 | P08574 | 0.00006 | 3.42 |
| ENSG00000106636 | YKT6 | YKT6 v-SNARE homolog | A4D2J0 | 0.00026 | 3.42 |
| ENSG00000057757 | PITHD1 | PITH domain containing 1 | Q9GZP4 | 0.000058 | 3.4 |
| ENSG00000143549 | TPM3 | tropomyosin 3 | A0A0S2Z4I4 | 0.000048 | 3.38 |
| ENSG00000132470 | ITGB4 | integrin subunit beta 4 | A0A024R8T0 | 0.000088 | 3.37 |
| ENSG00000066056 | TIE1 | tyrosine kinase with immunoglobulin like and EGF like domains 1 | B4DTW8 | 0.000041 | 3.33 |
| ENSG00000179051 | RCC2 | regulator of chromosome condensation 2 | A0A024RAC5 | 0.00022 | 3.3 |
| ENSG00000157916 | RER1 | retention in endoplasmic reticulum sorting receptor 1 | O15258 | 0.00026 | 3.3 |
| ENSG00000147155 | EBP | EBP cholestenol delta-isomerase | A0A024QYX0 | 0.00033 | 3.3 |
| ENSG00000022267 | FHL1 | four and a half LIM domains 1 | Q13642 | 0.000089 | 3.29 |
| ENSG00000110841 | PPFIBP1 | PPFIA binding protein 1 | A0A024RB02 | 0.00042 | 3.28 |
| ENSG00000112739 | PRPF4B | pre-mRNA processing factor 4B | A0A024QZY5 | 0.000062 | 3.27 |
| ENSG00000153904 | DDAH1 | dimethylarginine dimethylaminohydrolase 1 | B4E3V1 | 0.00013 | 3.26 |
| ENSG00000169067 | ACTBL2 | actin beta like 2 | Q562R1 | 0.00016 | 3.25 |
| ENSG00000105971 | CAV2 | caveolin 2 | P51636 | 0.00025 | 3.25 |
| ENSG00000110619 | CARS1 | cysteinyl-tRNA synthetase 1 | P49589 | 0.00025 | 3.22 |
| ENSG00000114416 | FXR1 | FMR1 autosomal homolog 1 | P51114 | 0.000074 | 3.21 |
| ENSG00000017260 | ATP2C1 | ATPase secretory pathway Ca2+ transporting 1 | P98194 | 0.000057 | 3.2 |
| ENSG00000139684 | ESD | esterase D | A0A140VJJ2 | 0.00033 | 3.18 |
| ENSG00000134001 | EIF2S1 | eukaryotic translation initiation factor 2 subunit alpha | P05198 | 0.0004 | 3.18 |
| ENSG00000100519 | PSMC6 | proteasome 26S subunit, ATPase 6 | A0A087X2I1 | 0.000054 | 3.17 |
| ENSG00000100030 | MAPK1 | mitogen-activated protein kinase 1 | P28482 | 0.000098 | 3.17 |
| ENSG00000103342 | GSPT1 | G1 to S phase transition 1 | P15170 | 0.000061 | 3.16 |
| ENSG00000130985 | UBA1 | ubiquitin like modifier activating enzyme 1 | A0A024R1A3 | 0.00023 | 3.16 |
| ENSG00000112306 | RPS12 | ribosomal protein S12 | P25398 | 0.00036 | 3.15 |
| ENSG00000135048 | CEMIP2 | cell migration inducing hyaluronidase 2 | Q9UHN6 | 0.000067 | 3.11 |

| | | | | | |
|---|---|---|---|---|---|
| ENSG00000138029 | HADHB | hydroxyacyl-CoA dehydrogenase trifunctional multienzyme complex subunit beta | P55084 | 0.000086 | 3.1 |
| ENSG00000008988 | RPS20 | ribosomal protein S20 | P60866 | 0.00045 | 3.1 |
| ENSG00000172725 | CORO1B | coronin 1B | A0A024R5K1 | 0.000078 | 3.09 |
| ENSG00000184584 | STING1 | stimulator of interferon response cGAMP interactor 1 | V5V0K2 | 0.000093 | 3.08 |
| ENSG00000138119 | MYOF | myoferlin | Q9NZM1 | 0.00028 | 3.01 |
| ENSG00000231500 | RPS18 | ribosomal protein S18 | P62269 | 0.00014 | 3 |
| ENSG00000111275 | ALDH2 | aldehyde dehydrogenase 2 family member | P05091 | 0.000094 | 2.98 |
| ENSG00000185825 | BCAP31 | B cell receptor associated protein 31 | P51572 | 0.00024 | 2.98 |
| ENSG00000163110 | PDLIM5 | PDZ and LIM domain 5 | Q96HC4 | 0.00024 | 2.97 |
| ENSG00000159840 | ZYX | zyxin | Q15942 | 0.00016 | 2.96 |
| ENSG00000087365 | SF3B2 | splicing factor 3b subunit 2 | Q13435 | 0.00028 | 2.96 |
| ENSG00000103018 | CYB5B | cytochrome b5 type B | J3KNF8 | 0.00063 | 2.96 |
| ENSG00000211899 | IGHM | immunoglobulin heavy constant mu | NA | 0.0051 | 2.95 |
| ENSG00000075624 | ACTB | actin beta | P60709 | 0.00035 | 2.93 |
| ENSG00000172354 | GNB2 | G protein subunit beta 2 | P62879 | 0.00092 | 2.93 |
| ENSG00000137486 | ARRB1 | arrestin beta 1 | B7Z1Q3 | 0.0006 | 2.92 |
| ENSG00000124181 | PLCG1 | phospholipase C gamma 1 | P19174 | 0.0018 | 2.92 |
| ENSG00000101361 | NOP56 | NOP56 ribonucleoprotein | O00567 | 0.00017 | 2.9 |
| ENSG00000067560 | RHOA | ras homolog family member A | A0A024R324 | 0.0052 | 2.9 |
| ENSG00000060237 | WNK1 | WNK lysine deficient protein kinase 1 | Q9H4A3 | 0.0063 | 2.9 |
| ENSG00000100823 | APEX1 | apurinic/apyrimidinic endodeoxyribonuclease 1 | P27695 | 0.00044 | 2.89 |
| ENSG00000110047 | EHD1 | EH domain containing 1 | B2R5U3 | 0.0001 | 2.87 |
| ENSG00000126261 | UBA2 | ubiquitin like modifier activating enzyme 2 | Q9UBT2 | 0.0002 | 2.87 |
| ENSG00000125970 | RALY | RALY heterogeneous nuclear ribonucleoprotein | Q9UKM9 | 0.0068 | 2.87 |
| ENSG00000105974 | CAV1 | caveolin 1 | A0A024R757 | 0.00011 | 2.86 |
| ENSG00000150768 | DLAT | dihydrolipoamide S-acetyltransferase | P10515 | 0.00044 | 2.86 |
| ENSG00000149218 | ENDOD1 | endonuclease domain containing 1 | O94919 | 0.0011 | 2.86 |
| ENSG00000214078 | CPNE1 | copine 1 | B0QZ18 | 0.004 | 2.86 |
| ENSG00000048828 | FAM120A | family with sequence similarity 120 member A | Q9NZB2 | 0.0011 | 2.84 |
| ENSG00000062822 | POLD1 | DNA polymerase delta 1, catalytic subunit | A0A024R4F4 | 0.0012 | 2.82 |
| ENSG00000115286 | NDUFS7 | NADH:ubiquinone oxidoreductase core subunit S7 | O75251 | 0.00027 | 2.81 |
| ENSG00000091879 | ANGPT2 | angiopoietin 2 | O15123 | 0.00048 | 2.81 |
| ENSG00000178695 | KCTD12 | potassium channel tetramerization domain containing 12 | A0A140VJM4 | 0.00017 | 2.8 |
| ENSG00000107438 | PDLIM1 | PDZ and LIM domain 1 | O00151 | 0.00044 | 2.8 |
| ENSG00000142634 | EFHD2 | EF-hand domain family member D2 | A0A024QZ77 | 0.00053 | 2.8 |

| | | | | | |
|---|---|---|---|---|---|
| ENSG00000142676 | RPL11 | ribosomal protein L11 | P62913 | 0.0017 | 2.8 |
| ENSG00000104419 | NDRG1 | N-myc downstream regulated 1 | Q92597 | 0.00082 | 2.79 |
| ENSG00000175931 | UBE2O | ubiquitin conjugating enzyme E2 O | Q9C0C9 | 0.00015 | 2.78 |
| ENSG00000141522 | ARHGDIA | Rho GDP dissociation inhibitor alpha | P52565 | 0.00059 | 2.78 |
| ENSG00000117054 | ACADM | acyl-CoA dehydrogenase medium chain | A0A0S2Z366 | 0.00071 | 2.77 |
| ENSG00000113441 | LNPEP | leucyl and cystinyl aminopeptidase | Q9UIQ6 | 0.00018 | 2.76 |
| ENSG00000171314 | PGAM1 | phosphoglycerate mutase 1 | B7Z9E5 | 0.00014 | 2.75 |
| ENSG00000244687 | UBE2V1 | ubiquitin conjugating enzyme E2 V1 | Q13404 | 0.00023 | 2.75 |
| ENSG00000241973 | PI4KA | phosphatidylinositol 4-kinase alpha | B4DYG5 | 0.0028 | 2.74 |
| ENSG00000132507 | EIF5A | eukaryotic translation initiation factor 5A | P63241 | 0.00036 | 2.72 |
| ENSG00000196923 | PDLIM7 | PDZ and LIM domain 7 | Q9NR12 | 0.00057 | 2.72 |
| ENSG00000117592 | PRDX6 | peroxiredoxin 6 | P30041 | 0.00066 | 2.72 |
| ENSG00000087274 | ADD1 | adducin 1 | P35611 | 0.00055 | 2.71 |
| ENSG00000122406 | RPL5 | ribosomal protein L5 | A2RUM7 | 0.00023 | 2.7 |
| ENSG00000082516 | GEMIN5 | gem nuclear organelle associated protein 5 | B7ZLC9 | 0.00024 | 2.7 |
| ENSG00000092841 | MYL6 | myosin light chain 6 | P60660 | 0.00059 | 2.7 |
| ENSG00000167792 | NDUFV1 | NADH:ubiquinone oxidoreductase core subunit V1 | P49821 | 0.00039 | 2.68 |
| ENSG00000116251 | RPL22 | ribosomal protein L22 | P35268 | 0.00049 | 2.68 |
| ENSG00000186642 | PDE2A | phosphodiesterase 2A | O00408 | 0.00046 | 2.66 |
| ENSG00000111737 | RAB35 | RAB35, member RAS oncogene family | Q15286 | 0.00035 | 2.64 |
| ENSG00000150593 | PDCD4 | programmed cell death 4 | B4DKX4 | 0.00062 | 2.64 |
| ENSG00000100220 | RTCB | RNA 2',3'-cyclic phosphate and 5'-OH ligase | Q9Y3I0 | 0.00041 | 2.63 |
| ENSG00000104408 | EIF3E | eukaryotic translation initiation factor 3 subunit E | P60228 | 0.00025 | 2.62 |
| ENSG00000158467 | AHCYL2 | adenosylhomocysteinase like 2 | Q96HN2 | 0.0023 | 2.62 |
| ENSG00000102898 | NUTF2 | nuclear transport factor 2 | A0A024R6Y2 | 0.0034 | 2.62 |
| ENSG00000165609 | NUDT5 | nudix hydrolase 5 | Q9UKK9 | 0.00038 | 2.61 |
| ENSG00000105323 | HNRNPUL1 | heterogeneous nuclear ribonucleoprotein U like 1 | B7Z4B8 | 0.00046 | 2.61 |
| ENSG00000164163 | ABCE1 | ATP binding cassette subfamily E member 1 | P61221 | 0.0006 | 2.61 |
| ENSG00000023228 | NDUFS1 | NADH:ubiquinone oxidoreductase core subunit S1 | P28331 | 0.0017 | 2.61 |
| ENSG00000128245 | YWHAH | tyrosine 3-monooxygenase/tryptophan 5-monooxygenase activation protein eta | A0A024R1K7 | 0.00063 | 2.6 |
| ENSG00000184009 | ACTG1 | actin gamma 1 | P63261 | 0.00064 | 2.6 |
| ENSG00000172757 | CFL1 | cofilin 1 | P23528 | 0.0009 | 2.6 |
| ENSG00000130741 | EIF2S3 | eukaryotic translation initiation factor 2 subunit gamma | P41091 | 0.0014 | 2.6 |
| ENSG00000108848 | LUC7L3 | LUC7 like 3 pre-mRNA splicing factor | J3KPP4 | 0.00025 | 2.59 |
| ENSG00000011485 | PPP5C | protein phosphatase 5 catalytic subunit | A0A024R0Q7 | 0.00024 | 2.58 |

| | | | | | |
|---|---|---|---|---|---|
| ENSG00000140350 | ANP32A | acidic nuclear phosphoprotein 32 family member A | P39687 | 0.00078 | 2.58 |
| ENSG00000141429 | GALNT1 | polypeptide N-acetylgalactosaminyltransferase 1 | A0A024RC48 | 0.0011 | 2.58 |
| ENSG00000115091 | ACTR3 | actin related protein 3 | B4DXW1 | 0.0021 | 2.58 |
| ENSG00000104687 | GSR | glutathione-disulfide reductase | P00390 | 0.0012 | 2.57 |
| ENSG00000149480 | MTA2 | metastasis associated 1 family member 2 | O94776 | 0.0022 | 2.56 |
| ENSG00000136938 | ANP32B | acidic nuclear phosphoprotein 32 family member B | Q92688 | 0.003 | 2.56 |
| ENSG00000100243 | CYB5R3 | cytochrome b5 reductase 3 | P00387 | 0.00047 | 2.54 |
| ENSG00000162616 | DNAJB4 | DnaJ heat shock protein family (Hsp40) member B4 | Q9UDY4 | 0.00051 | 2.53 |
| ENSG00000197958 | RPL12 | ribosomal protein L12 | P30050 | 0.00026 | 2.52 |
| ENSG00000165637 | VDAC2 | voltage dependent anion channel 2 | P45880 | 0.00074 | 2.52 |
| ENSG00000070756 | PABPC1 | poly(A) binding protein cytoplasmic 1 | A0A024R9C1 | 0.001 | 2.51 |
| ENSG00000100097 | LGALS1 | galectin 1 | P09382 | 0.0021 | 2.51 |
| ENSG00000141456 | PELP1 | proline, glutamate and leucine rich protein 1 | B4DEX7 | 0.011 | 2.51 |
| ENSG00000130429 | ARPC1B | actin related protein 2/3 complex subunit 1B | A4D275 | 0.0006 | 2.49 |
| ENSG00000167699 | GLOD4 | glyoxalase domain containing 4 | Q9HC38 | 0.00038 | 2.48 |
| ENSG00000137497 | NUMA1 | nuclear mitotic apparatus protein 1 | A0A024R5M9 | 0.0015 | 2.48 |
| ENSG00000162521 | RBBP4 | RB binding protein 4, chromatin remodeling factor | Q09028 | 0.0022 | 2.48 |
| ENSG00000142507 | PSMB6 | proteasome 20S subunit beta 6 | A0A087X2I4 | 0.003 | 2.48 |
| ENSG00000213639 | PPP1CB | protein phosphatase 1 catalytic subunit beta | P62140 | 0.00085 | 2.47 |
| ENSG00000103335 | PIEZO1 | piezo type mechanosensitive ion channel component 1 | Q92508 | 0.0024 | 2.47 |
| ENSG00000120438 | TCP1 | t-complex 1 | E7EQR6 | 0.00061 | 2.45 |
| ENSG00000169710 | FASN | fatty acid synthase | P49327 | 0.00068 | 2.43 |
| ENSG00000108518 | PFN1 | profilin 1 | P07737 | 0.0016 | 2.43 |
| ENSG00000136930 | PSMB7 | proteasome 20S subunit beta 7 | E9KL30 | 0.0016 | 2.43 |
| ENSG00000090615 | GOLGA3 | golgin A3 | Q08378 | 0.022 | 2.43 |
| ENSG00000106397 | PLOD3 | procollagen-lysine,2-oxoglutarate 5-dioxygenase 3 | O60568 | 0.00096 | 2.42 |
| ENSG00000169021 | UQCRFS1 | ubiquinol-cytochrome c reductase, Rieske iron-sulfur polypeptide 1 | P47985 | 0.0032 | 2.42 |
| ENSG00000140750 | ARHGAP17 | Rho GTPase activating protein 17 | Q68EM7 | 0.02 | 2.42 |
| ENSG00000030066 | NUP160 | nucleoporin 160 | Q12769 | 0.00074 | 2.41 |
| ENSG00000240972 | MIF | macrophage migration inhibitory factor | I4AY87 | 0.00078 | 2.41 |
| ENSG00000130309 | COLGALT1 | collagen beta(1-O)galactosyltransferase 1 | Q8NBJ5 | 0.001 | 2.41 |
| ENSG00000126602 | TRAP1 | TNF receptor associated protein 1 | Q12931 | 0.017 | 2.41 |
| ENSG00000169504 | CLIC4 | chloride intracellular channel 4 | Q6FIC5 | 0.00078 | 2.38 |
| ENSG00000185236 | RAB11B | RAB11B, member RAS oncogene family | Q15907 | 0.00081 | 2.38 |

| | | | | | |
|---|---|---|---|---|---|
| ENSG00000138668 | HNRNPD | heterogeneous nuclear ribonucleoprotein D | Q14103 | 0.00068 | 2.37 |
| ENSG00000135926 | TMBIM1 | transmembrane BAX inhibitor motif containing 1 | A0A024R472 | 0.0009 | 2.36 |
| ENSG00000182774 | RPS17 | ribosomal protein S17 | P08708 | 0.0012 | 2.35 |
| ENSG00000001497 | LAS1L | LAS1 like ribosome biogenesis factor | Q9Y4W2 | 0.018 | 2.35 |
| ENSG00000119707 | RBM25 | RNA binding motif protein 25 | P49756 | 0.0015 | 2.34 |
| ENSG00000078369 | GNB1 | G protein subunit beta 1 | B3KVK2 | 0.0034 | 2.33 |
| ENSG00000120254 | MTHFD1L | methylenetetrahydrofolate dehydrogenase (NADP+ dependent) 1 like | B7ZM99 | 0.00045 | 2.32 |
| ENSG00000164924 | YWHAZ | tyrosine 3-monooxygenase/tryptophan 5-monooxygenase activation protein zeta | D0PNI1 | 0.0012 | 2.31 |
| ENSG00000198363 | ASPH | aspartate beta-hydroxylase | Q12797 | 0.0013 | 2.31 |
| ENSG00000067113 | PLPP1 | phospholipid phosphatase 1 | A0A024QZS3 | 0.0064 | 2.31 |
| ENSG00000133112 | TPT1 | tumor protein, translationally-controlled 1 | A0A0B4J2C3 | 0.028 | 2.31 |
| ENSG00000175792 | RUVBL1 | RuvB like AAA ATPase 1 | Q9Y265 | 0.00078 | 2.3 |
| ENSG00000135677 | GNS | glucosamine (N-acetyl)-6-sulfatase | A0A024RBC5 | 0.0032 | 2.3 |
| ENSG00000213719 | CLIC1 | chloride intracellular channel 1 | O00299 | 0.0012 | 2.29 |
| ENSG00000169714 | CNBP | CCHC-type zinc finger nucleic acid binding protein | P62633 | 0.0018 | 2.29 |
| ENSG00000132341 | RAN | RAN, member RAS oncogene family | B4DV51 | 0.0049 | 2.29 |
| ENSG00000142910 | TINAGL1 | tubulointerstitial nephritis antigen like 1 | Q9GZM7 | 0.00095 | 2.28 |
| ENSG00000083845 | RPS5 | ribosomal protein S5 | A0A024R4Q8 | 0.0017 | 2.27 |
| ENSG00000157020 | SEC13 | SEC13 homolog, nuclear pore and COPII coat complex component | P55735 | 0.0019 | 2.27 |
| ENSG00000167978 | SRRM2 | serine/arginine repetitive matrix 2 | A0A140VK53 | 0.0012 | 2.26 |
| NA | MT-CO2 | NA | NA | 0.0016 | 2.25 |
| ENSG00000124570 | SERPINB6 | serpin family B member 6 | A0A024QZX5 | 0.0024 | 2.25 |
| ENSG00000161011 | SQSTM1 | sequestosome 1 | Q13501 | 0.0085 | 2.25 |
| ENSG00000189403 | HMGB1 | high mobility group box 1 | A0A024RDR0 | 0.0012 | 2.24 |
| ENSG00000169100 | SLC25A6 | solute carrier family 25 member 6 | P12236 | 0.0026 | 2.24 |
| ENSG00000122565 | CBX3 | chromobox 3 | A4D177 | 0.0029 | 2.24 |
| ENSG00000143416 | SELENBP1 | selenium binding protein 1 | Q13228 | 0.03 | 2.23 |
| ENSG00000142937 | RPS8 | ribosomal protein S8 | P62241 | 0.0013 | 2.22 |
| ENSG00000116005 | PCYOX1 | prenylcysteine oxidase 1 | Q9UHG3 | 0.0023 | 2.22 |
| ENSG00000115380 | EFEMP1 | EGF containing fibulin extracellular matrix protein 1 | A0A0S2Z4F1 | 0.0021 | 2.21 |
| ENSG00000196230 | TUBB | tubulin beta class I | B4DY90 | 0.0054 | 2.21 |
| ENSG00000148700 | ADD3 | adducin 3 | Q9UEY8 | 0.00089 | 2.2 |
| ENSG00000121774 | KHDRBS1 | KH RNA binding domain containing, signal transduction associated 1 | Q07666 | 0.0041 | 2.2 |
| ENSG00000154473 | BUB3 | BUB3 mitotic checkpoint protein | A0A140VJF3 | 0.0042 | 2.19 |

| | | | | | |
|---|---|---|---|---|---|
| ENSG00000100504 | PYGL | glycogen phosphorylase L | P06737 | 0.016 | 2.18 |
| ENSG00000107862 | GBF1 | golgi brefeldin A resistant guanine nucleotide exchange factor 1 | Q92538 | 0.0013 | 2.16 |
| ENSG00000203879 | GDI1 | GDP dissociation inhibitor 1 | A0A0S2Z3X8 | 0.00073 | 2.15 |
| ENSG00000101294 | HM13 | histocompatibility minor 13 | A0A0S2Z5V7 | 0.0014 | 2.15 |
| ENSG00000176619 | LMNB2 | lamin B2 | Q03252 | 0.0016 | 2.15 |
| ENSG00000100280 | AP1B1 | adaptor related protein complex 1 subunit beta 1 | Q10567 | 0.0063 | 2.15 |
| ENSG00000148834 | GSTO1 | glutathione S-transferase omega 1 | P78417 | 0.0012 | 2.14 |
| ENSG00000168259 | DNAJC7 | DnaJ heat shock protein family (Hsp40) member C7 | Q99615 | 0.0025 | 2.14 |
| ENSG00000104325 | DECR1 | 2,4-dienoyl-CoA reductase 1 | Q16698 | 0.0036 | 2.13 |
| ENSG00000108671 | PSMD11 | proteasome 26S subunit, non-ATPase 11 | O00231 | 0.0012 | 2.12 |
| ENSG00000092199 | HNRNPC | heterogeneous nuclear ribonucleoprotein C | P07910 | 0.0022 | 2.12 |
| ENSG00000165280 | VCP | valosin containing protein | P55072 | 0.0048 | 2.12 |
| ENSG00000131016 | AKAP12 | A-kinase anchoring protein 12 | Q02952 | 0.0013 | 2.11 |
| ENSG00000197111 | PCBP2 | poly(rC) binding protein 2 | Q15366 | 0.0021 | 2.11 |
| ENSG00000163513 | TGFBR2 | transforming growth factor beta receptor 2 | D2JYI1 | 0.0022 | 2.11 |
| ENSG00000170027 | YWHAG | tyrosine 3-monooxygenase/tryptophan 5-monooxygenase activation protein gamma | P61981 | 0.0028 | 2.11 |
| ENSG00000076201 | PTPN23 | protein tyrosine phosphatase non-receptor type 23 | B4DST5 | 0.0055 | 2.11 |
| ENSG00000125977 | EIF2S2 | eukaryotic translation initiation factor 2 subunit beta | P20042 | 0.0047 | 2.1 |
| ENSG00000127483 | HP1BP3 | heterochromatin protein 1 binding protein 3 | Q5SSJ5 | 0.0066 | 2.1 |
| ENSG00000158710 | TAGLN2 | transgelin 2 | P37802 | 0.013 | 2.1 |
| ENSG00000112118 | MCM3 | minichromosome maintenance complex component 3 | B4DUQ9 | 0.028 | 2.1 |
| ENSG00000188257 | PLA2G2A | phospholipase A2 group IIA | A0A024RA96 | 0.036 | 2.1 |
| ENSG00000109501 | WFS1 | wolframin ER transmembrane glycoprotein | A0A0S2Z4V6 | 0.0012 | 2.09 |
| ENSG00000159176 | CSRP1 | cysteine and glycine rich protein 1 | B4DY28 | 0.0031 | 2.09 |
| ENSG00000204628 | RACK1 | receptor for activated C kinase 1 | E9KL35 | 0.0044 | 2.09 |
| ENSG00000150753 | CCT5 | chaperonin containing TCP1 subunit 5 | B4DX08 | 0.0012 | 2.06 |
| ENSG00000108515 | ENO3 | enolase 3 | P13929 | 0.0015 | 2.06 |
| ENSG00000198561 | CTNND1 | catenin delta 1 | O60716 | 0.002 | 2.06 |
| ENSG00000100911 | PSME2 | proteasome activator subunit 2 | Q86SZ7 | 0.0037 | 2.06 |
| ENSG00000116288 | PARK7 | Parkinsonism associated deglycase | Q99497 | 0.0015 | 2.05 |
| ENSG00000121152 | NCAPH | non-SMC condensin I complex subunit H | Q15003 | 0.0056 | 2.05 |
| ENSG00000067369 | TP53BP1 | tumor protein p53 binding protein 1 | Q12888 | 0.025 | 2.05 |
| ENSG00000166794 | PPIB | peptidylprolyl isomerase B | P23284 | 0.0015 | 2.04 |
| ENSG00000170323 | FABP4 | fatty acid binding protein 4 | E7DVW4 | 0.0036 | 2.04 |

| | | | | | |
|---|---|---|---|---|---|
| ENSG00000130158 | DOCK6 | dedicator of cytokinesis 6 | B7Z9U8 | 0.005 | 2.03 |
| ENSG00000095380 | NANS | N-acetylneuraminate synthase | Q9NR45 | 0.0038 | 2.02 |
| ENSG00000170348 | TMED10 | transmembrane p24 trafficking protein 10 | A0A024R6I3 | 0.0055 | 2.02 |
| ENSG00000084652 | TXLNA | taxilin alpha | P40222 | 0.0019 | 2.01 |
| ENSG00000198898 | CAPZA2 | capping actin protein of muscle Z-line subunit alpha 2 | A4D0V4 | 0.0024 | 2.01 |
| ENSG00000176986 | SEC24C | SEC24 homolog C, COPII coat complex component | A0A024QZM6 | 0.02 | 2.01 |
| ENSG00000146731 | CCT6A | chaperonin containing TCP1 subunit 6A | P40227 | 0.0048 | 2 |
| ENSG00000091164 | TXNL1 | thioredoxin like 1 | O43396 | 0.0059 | 2 |
| ENSG00000113569 | NUP155 | nucleoporin 155 | B4DLT2 | 0.0065 | 1.99 |
| ENSG00000144746 | ARL6IP5 | ADP ribosylation factor like GTPase 6 interacting protein 5 | A0A024R371 | 0.034 | 1.99 |
| ENSG00000113758 | DBN1 | drebrin 1 | Q16643 | 0.0032 | 1.97 |
| ENSG00000163131 | CTSS | cathepsin S | P25774 | 0.0017 | 1.96 |
| ENSG00000107719 | PALD1 | phosphatase domain containing paladin 1 | A0A024QZM5 | 0.0018 | 1.96 |
| ENSG00000161057 | PSMC2 | proteasome 26S subunit, ATPase 2 | B7Z571 | 0.0033 | 1.96 |
| ENSG00000004700 | RECQL | RecQ like helicase | A0A024RAV2 | 0.0039 | 1.96 |
| ENSG00000054654 | SYNE2 | spectrin repeat containing nuclear envelope protein 2 | Q8WXH0 | 0.0048 | 1.96 |
| ENSG00000140416 | TPM1 | tropomyosin 1 | A0A0K0K1I0 | 0.0016 | 1.95 |
| ENSG00000211623 | IGKV2D-26 | immunoglobulin kappa variable 2D-26 | NA | 0.027 | 1.95 |
| ENSG00000110107 | PRPF19 | pre-mRNA processing factor 19 | Q9UMS4 | 0.002 | 1.94 |
| ENSG00000062485 | CS | citrate synthase | A0A024RB75 | 0.015 | 1.94 |
| ENSG00000177674 | AGTRAP | angiotensin II receptor associated protein | Q6RW13 | 0.0029 | 1.93 |
| ENSG00000166333 | ILK | integrin linked kinase | Q13418 | 0.013 | 1.93 |
| ENSG00000111144 | LTA4H | leukotriene A4 hydrolase | A0A140VK27 | 0.0018 | 1.92 |
| ENSG00000140105 | WARS1 | tryptophanyl-tRNA synthetase 1 | A0A024R6K8 | 0.0019 | 1.92 |
| ENSG00000075415 | SLC25A3 | solute carrier family 25 member 3 | A0A024RBE8 | 0.0023 | 1.92 |
| ENSG00000085832 | EPS15 | epidermal growth factor receptor pathway substrate 15 | B7Z240 | 0.0068 | 1.92 |
| ENSG00000138772 | ANXA3 | annexin A3 | P12429 | 0.0017 | 1.91 |
| ENSG00000127022 | CANX | calnexin | P27824 | 0.0018 | 1.91 |
| ENSG00000173267 | SNCG | synuclein gamma | F8W754 | 0.0076 | 1.91 |
| ENSG00000137509 | PRCP | prolylcarboxypeptidase | B7Z7Q6 | 0.024 | 1.91 |
| ENSG00000073969 | NSF | N-ethylmaleimide sensitive factor, vesicle fusing ATPase | P46459 | 0.0019 | 1.9 |
| ENSG00000005022 | SLC25A5 | solute carrier family 25 member 5 | P05141 | 0.0029 | 1.9 |
| ENSG00000171793 | CTPS1 | CTP synthase 1 | B4E1E0 | 0.0038 | 1.9 |
| ENSG00000175203 | DCTN2 | dynactin subunit 2 | Q13561 | 0.0066 | 1.9 |
| ENSG00000168003 | SLC3A2 | solute carrier family 3 member 2 | J3KPF3 | 0.0028 | 1.89 |
| ENSG00000114573 | ATP6V1A | ATPase H+ transporting V1 subunit A | P38606 | 0.0031 | 1.89 |

| | | | | | |
|---|---|---|---|---|---|
| ENSG00000058262 | SEC61A1 | SEC61 translocon subunit alpha 1 | B3KME8 | 0.0051 | 1.89 |
| ENSG00000119689 | DLST | dihydrolipoamide S-succinyltransferase | B7Z6J1 | 0.012 | 1.88 |
| ENSG00000167674 | HDGFL2 | HDGF like 2 | Q7Z4V5 | 0.0055 | 1.87 |
| ENSG00000170144 | HNRNPA3 | heterogeneous nuclear ribonucleoprotein A3 | B4DDB6 | 0.0044 | 1.86 |
| ENSG00000165092 | ALDH1A1 | aldehyde dehydrogenase 1 family member A1 | P00352 | 0.0046 | 1.86 |
| ENSG00000168610 | STAT3 | signal transducer and activator of transcription 3 | P40763 | 0.0047 | 1.86 |
| ENSG00000145833 | DDX46 | DEAD-box helicase 46 | A0A0C4DG89 | 0.0065 | 1.86 |
| ENSG00000005893 | LAMP2 | lysosomal associated membrane protein 2 | P13473 | 0.012 | 1.86 |
| ENSG00000102119 | EMD | emerin | P50402 | 0.02 | 1.86 |
| ENSG00000110090 | CPT1A | carnitine palmitoyltransferase 1A | P50416 | 0.0036 | 1.85 |
| ENSG00000149084 | HSD17B12 | hydroxysteroid 17-beta dehydrogenase 12 | Q53GQ0 | 0.019 | 1.85 |
| ENSG00000151176 | PLBD2 | phospholipase B domain containing 2 | Q8NHP8 | 0.0044 | 1.84 |
| ENSG00000125166 | GOT2 | glutamic-oxaloacetic transaminase 2 | P00505 | 0.0036 | 1.83 |
| ENSG00000198755 | RPL10A | ribosomal protein L10a | P62906 | 0.0053 | 1.83 |
| ENSG00000119655 | NPC2 | NPC intracellular cholesterol transporter 2 | A0A024R6C0 | 0.0074 | 1.83 |
| ENSG00000067225 | PKM | pyruvate kinase M1/2 | P14618 | 0.004 | 1.81 |
| ENSG00000159140 | SON | SON DNA and RNA binding protein | P18583 | 0.0073 | 1.81 |
| ENSG00000113643 | RARS1 | arginyl-tRNA synthetase 1 | P54136 | 0.0051 | 1.8 |
| ENSG00000104388 | RAB2A | RAB2A, member RAS oncogene family | P61019 | 0.008 | 1.8 |
| ENSG00000197635 | DPP4 | dipeptidyl peptidase 4 | P27487 | 0.018 | 1.8 |
| ENSG00000140740 | UQCRC2 | ubiquinol-cytochrome c reductase core protein 2 | P22695 | 0.022 | 1.8 |
| ENSG00000023191 | RNH1 | ribonuclease/angiogenin inhibitor 1 | A0A140VJT8 | 0.0055 | 1.79 |
| ENSG00000169223 | LMAN2 | lectin, mannose binding 2 | Q12907 | 0.0059 | 1.79 |
| ENSG00000172037 | LAMB2 | laminin subunit beta 2 | A0A024R319 | 0.0066 | 1.78 |
| ENSG00000149273 | RPS3 | ribosomal protein S3 | P23396 | 0.0073 | 1.78 |
| ENSG00000162402 | USP24 | ubiquitin specific peptidase 24 | Q9UPU5 | 0.047 | 1.78 |
| ENSG00000123416 | TUBA1B | tubulin alpha 1b | P68363 | 0.0048 | 1.77 |
| ENSG00000066777 | ARFGEF1 | ADP ribosylation factor guanine nucleotide exchange factor 1 | A0A024R7X0 | 0.0079 | 1.77 |
| ENSG00000147140 | NONO | non-POU domain containing octamer binding | A0A0S2Z4Z9 | 0.026 | 1.77 |
| ENSG00000132780 | NASP | nuclear autoantigenic sperm protein | P49321 | 0.005 | 1.76 |
| ENSG00000196497 | IPO4 | importin 4 | B3KT38 | 0.01 | 1.76 |
| ENSG00000175130 | MARCKSL1 | MARCKS like 1 | P49006 | 0.017 | 1.76 |
| ENSG00000089157 | RPLP0 | ribosomal protein lateral stalk subunit P0 | A0A024RBS2 | 0.0039 | 1.75 |
| ENSG00000186468 | RPS23 | ribosomal protein S23 | A8K517 | 0.0069 | 1.75 |

Adjustments of p-values for multiple comparisons were used with Benjamini-Hochberg (BH) correction.

**Supplemental Table S6.** LEC protein palmitoylated enriched only in siCtrl and insulin treatment condition.

| ENSEMBL ID | Description | UNIPROT | Enrichment in siCtrl + Ins | |
|---|---|---|---|---|
| | | | *p*-value | -Log Ratio (+/- HA) |
| ENSG00000075239 | acetyl-CoA acetyltransferase 1 | A0A140VJX1 | 0.0063 | 1.5 |
| ENSG00000014257 | acid phosphatase 3 | P15309 | 0.035 | 2.46 |
| ENSG00000137845 | ADAM metallopeptidase domain 10 | A0A024R5U5 | 0.042 | 1.01 |
| ENSG00000162618 | adhesion G protein-coupled receptor L4 | Q9HBW9 | 2.00E-06 | 5.66 |
| ENSG00000119711 | aldehyde dehydrogenase 6 family member A1 | Q02252 | 0.036 | 1.06 |
| ENSG00000084674 | apolipoprotein B | P04114 | 0.0062 | 2.7 |
| ENSG00000145349 | calcium/calmodulin dependent protein kinase II delta | A0A024RDK3 | 0.019 | 1.43 |
| ENSG00000147419 | coiled-coil domain containing 25 | G3V121 | 0.04 | 2.05 |
| ENSG00000135218 | CD36 molecule | A4D1B1 | 0.0014 | 2.13 |
| ENSG00000004468 | CD38 molecule | B4E006 | 8.00E-05 | 3.66 |
| ENSG00000026508 | CD44 molecule (Indian blood group) | P16070 | 4.10E-06 | 4.36 |
| ENSG00000013297 | claudin 11 | O75508 | 7.10E-06 | 4.63 |
| ENSG00000153551 | CKLF like MARVEL transmembrane domain containing 7 | A0A024R2L3 | 7.20E-06 | 4.95 |
| ENSG00000158270 | collectin subfamily member 12 | Q5KU26 | 0.013 | 1.48 |
| ENSG00000136160 | endothelin receptor type B | P24530 | 0.0066 | 1.82 |
| ENSG00000181104 | coagulation factor II thrombin receptor | P25116 | 2.70E-05 | 3.67 |
| ENSG00000087086 | ferritin light chain | P02792 | 0.0073 | 1.76 |
| ENSG00000198380 | glutamine--fructose-6-phosphate transaminase 1 | Q06210 | 0.026 | 1.16 |
| ENSG00000196329 | GTPase, IMAP family member 5 | A0A090N8P9 | 0.022 | 1.64 |
| ENSG00000139433 | glycolipid transfer protein | A0A024RBI7 | 0.0011 | 2.78 |
| ENSG00000135821 | glutamate-ammonia ligase | A8YXX4 | 0.021 | 4.1 |
| ENSG00000088256 | G protein subunit alpha 11 | P29992 | 3.70E-05 | 4.08 |
| ENSG00000120063 | G protein subunit alpha 13 | Q14344 | 2.40E-06 | 5.81 |
| ENSG00000065135 | G protein subunit alpha i3 | P08754 | 0.00078 | 2.18 |
| ENSG00000156052 | G protein subunit alpha q | A0A024R240 | 8.20E-07 | 5.91 |
| ENSG00000147533 | golgin A7 | Q7Z5G4 | 5.90E-06 | 4.62 |
| ENSG00000115159 | glycerol-3-phosphate dehydrogenase 2 | P43304 | 0.024 | 1.18 |
| ENSG00000074696 | 3-hydroxyacyl-CoA dehydratase 3 | Q9P035 | 0.0012 | 2.28 |
| ENSG00000063854 | hydroxyacylglutathione hydrolase | Q16775 | 0.022 | 3.4 |
| ENSG00000004961 | holocytochrome c synthase | A0A024RBY9 | 0.00077 | 2.33 |
| ENSG00000148634 | HECT and RLD domain containing E3 ubiquitin protein ligase 4 | Q5GLZ8 | 0.007 | 1.36 |
| NA | major histocompatibility complex, class I, H (pseudogene) | NA | 0.043 | 1.71 |
| ENSG00000103415 | heme oxygenase 2 | P30519 | 0.012 | 1.28 |
| ENSG00000197081 | insulin like growth factor 2 receptor | P11717 | 0.034 | 1.19 |
| ENSG00000136688 | interleukin 36 gamma | Q9NZH8 | 0.00084 | 2.96 |
| ENSG00000167754 | kallikrein related peptidase 5 | Q9Y337 | 0.0093 | 2.04 |

| | | | | |
|---|---|---|---|---|
| ENSG00000213625 | leptin receptor overlapping transcript | A0A087X0N2 | 8.20E-05 | 3.42 |
| ENSG00000205076 | galectin 7 | P47929 | 0.041 | 2.7 |
| ENSG00000100258 | lipase maturation factor 2 | Q9BU23 | 4.20E-05 | 3.65 |
| ENSG00000254087 | LYN proto-oncogene, Src family tyrosine kinase | P07948 | 0.00022 | 2.92 |
| ENSG00000156026 | mitochondrial calcium uniporter | Q8NE86 | 0.0053 | 1.99 |
| ENSG00000157227 | matrix metallopeptidase 14 | P50281 | 7.70E-05 | 2.96 |
| ENSG00000262814 | mitochondrial ribosomal protein L12 | P52815 | 0.019 | 1.34 |
| ENSG00000204568 | mitochondrial ribosomal protein S18B | B0S7P4 | 0.0011 | 2.52 |
| ENSG00000125148 | metallothionein 2A | P02795 | 0.00089 | 3.08 |
| ENSG00000147649 | metadherin | A0A024R9D2 | 0.00078 | 2.05 |
| ENSG00000144959 | neutral cholesterol ester hydrolase 1 | A0A0A0MTJ9 | 4.60E-05 | 3.82 |
| ENSG00000139180 | NADH:ubiquinone oxidoreductase subunit A9 | Q16795 | 0.034 | 2.27 |
| ENSG00000135124 | purinergic receptor P2X 4 | Q99571 | 9.10E-05 | 3.2 |
| ENSG00000184363 | plakophilin 3 | Q9Y446 | 0.031 | 2.61 |
| ENSG00000073008 | PVR cell adhesion molecule | P15151 | 0.0049 | 1.65 |
| ENSG00000123728 | RAP2C, member of RAS oncogene family | Q9Y3L5 | 6.50E-06 | 4.04 |
| ENSG00000239306 | RNA binding motif protein 14 | A0A0S2Z567 | 0.038 | 2.09 |
| ENSG00000131378 | raftlin, lipid raft linker 1 | Q14699 | 0.00043 | 2.5 |
| ENSG00000205937 | RNA binding protein with serine rich domain 1 | D3DU92 | 0.0054 | 2.53 |
| ENSG00000063177 | ribosomal protein L18 | A0A024QZD1 | 0.0015 | 1.81 |
| ENSG00000144713 | ribosomal protein L32 | A0A024R2G7 | 0.0093 | 1.35 |
| ENSG00000109475 | ribosomal protein L34 | A0A024RDH8 | 0.029 | 1.27 |
| ENSG00000130255 | ribosomal protein L36 | Q9Y3U8 | 0.012 | 1.55 |
| ENSG00000174444 | ribosomal protein L4 | P36578 | 0.0096 | 1.32 |
| ENSG00000197728 | ribosomal protein S26 | A0A024RB14 | 0.0096 | 2.19 |
| ENSG00000170889 | ribosomal protein S9 | A0A024R4M0 | 0.0016 | 2.18 |
| ENSG00000126458 | RAS related | A0A024QZF2 | 7.40E-07 | 5.94 |
| ENSG00000143546 | S100 calcium binding protein A8 | P05109 | 0.046 | 2.79 |
| ENSG00000140497 | secretory carrier membrane protein 2 | A8K769 | 8.40E-06 | 4.1 |
| ENSG00000183291 | selenoprotein F | O60613 | 0.00057 | 2.26 |
| ENSG00000057149 | serpin family B member 3 | P29508 | 0.0068 | 3.7 |
| ENSG00000206073 | serpin family B member 4 | P48594 | 0.023 | 3.42 |
| ENSG00000155380 | solute carrier family 16 member 1 | A0A024R0H1 | 0.01 | 1.86 |
| ENSG00000146411 | solute carrier family 2 member 12 | Q8TD20 | 0.00055 | 3.36 |
| ENSG00000104852 | small nuclear ribonucleoprotein U1 subunit 70 | P08621 | 2.70E-05 | 3.46 |
| ENSG00000133226 | serine and arginine repetitive matrix 1 | B7Z7U0 | 1.90E-05 | 3.65 |
| ENSG00000116754 | serine and arginine rich splicing factor 11 | Q05519 | 1.10E-05 | 4.73 |
| ENSG00000116350 | serine and arginine rich splicing factor 4 | Q08170 | 0.025 | 1.59 |
| ENSG00000168394 | transporter 1, ATP binding cassette subfamily B member | A0A0S2Z5A6 | 0.00028 | 2.39 |
| ENSG00000130193 | thioesterase superfamily member 6 | Q8WUY1 | 0.0058 | 1.76 |
| ENSG00000066654 | THUMP domain containing 1 | A0A024R388 | 0.018 | 1.72 |
| ENSG00000099203 | transmembrane p24 trafficking protein 1 | Q13445 | 3.30E-06 | 4.91 |

| | | | | |
|---|---|---|---|---|
| ENSG00000109084 | transmembrane protein 97 | Q5BJF2 | 0.0093 | 3.16 |
| ENSG00000166479 | thioredoxin related transmembrane protein 3 | Q96JJ7 | 0.027 | 1.08 |
| ENSG00000187688 | transient receptor potential cation channel subfamily V member 2 | Q9Y5S1 | 0.00014 | 2.73 |
| ENSG00000000003 | tetraspanin 6 | A0A024RCI0 | 0.0021 | 2.85 |
| ENSG00000127824 | tubulin alpha 4a | P68366 | 0.0081 | 1.63 |
| ENSG00000184470 | thioredoxin reductase 2 | E7EWK1 | 9.70E-05 | 3.14 |
| ENSG00000025708 | thymidine phosphorylase | B2RBL3 | 0.048 | 2.16 |
| ENSG00000072401 | ubiquitin conjugating enzyme E2 D1 | A0A087WW00 | 0.00073 | 3.11 |
| ENSG00000198833 | ubiquitin conjugating enzyme E2 J1 | Q9Y385 | 0.00031 | 2.76 |
| ENSG00000159202 | ubiquitin conjugating enzyme E2 Z | Q9H832 | 0.00066 | 2.24 |
| ENSG00000114062 | ubiquitin protein ligase E3A | Q05086 | 4.10E-05 | 3.79 |
| ENSG00000074755 | zinc finger ZZ-type and EF-hand domain containing 1 | O43149 | 0.0096 | 1.45 |
