## Supplemental Figures S1-S3 for "Insulin regulates lymphatic endothelial function via palmitoylation"

### Slide 1
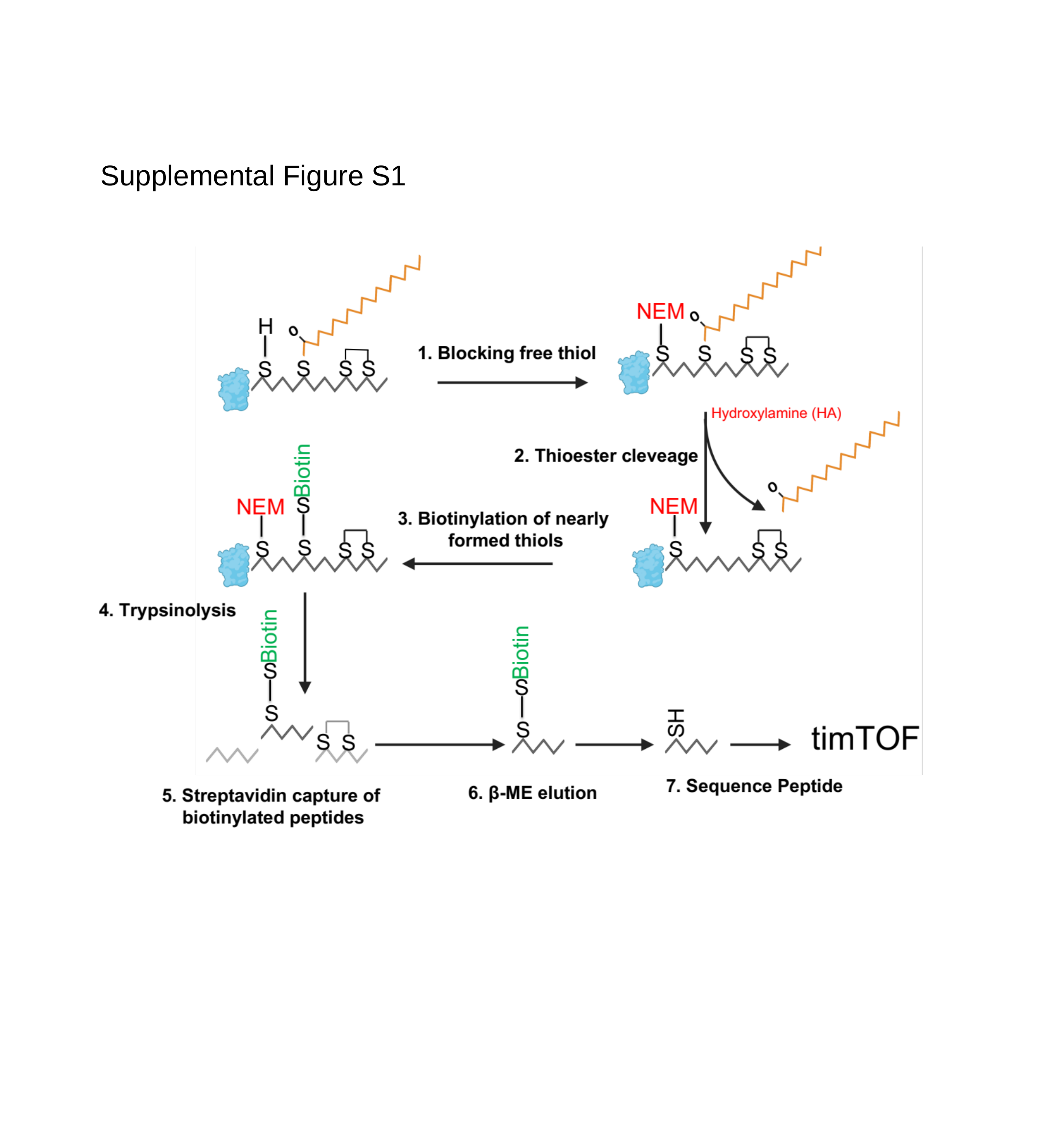

Supplemental Figure S1

### Slide 2
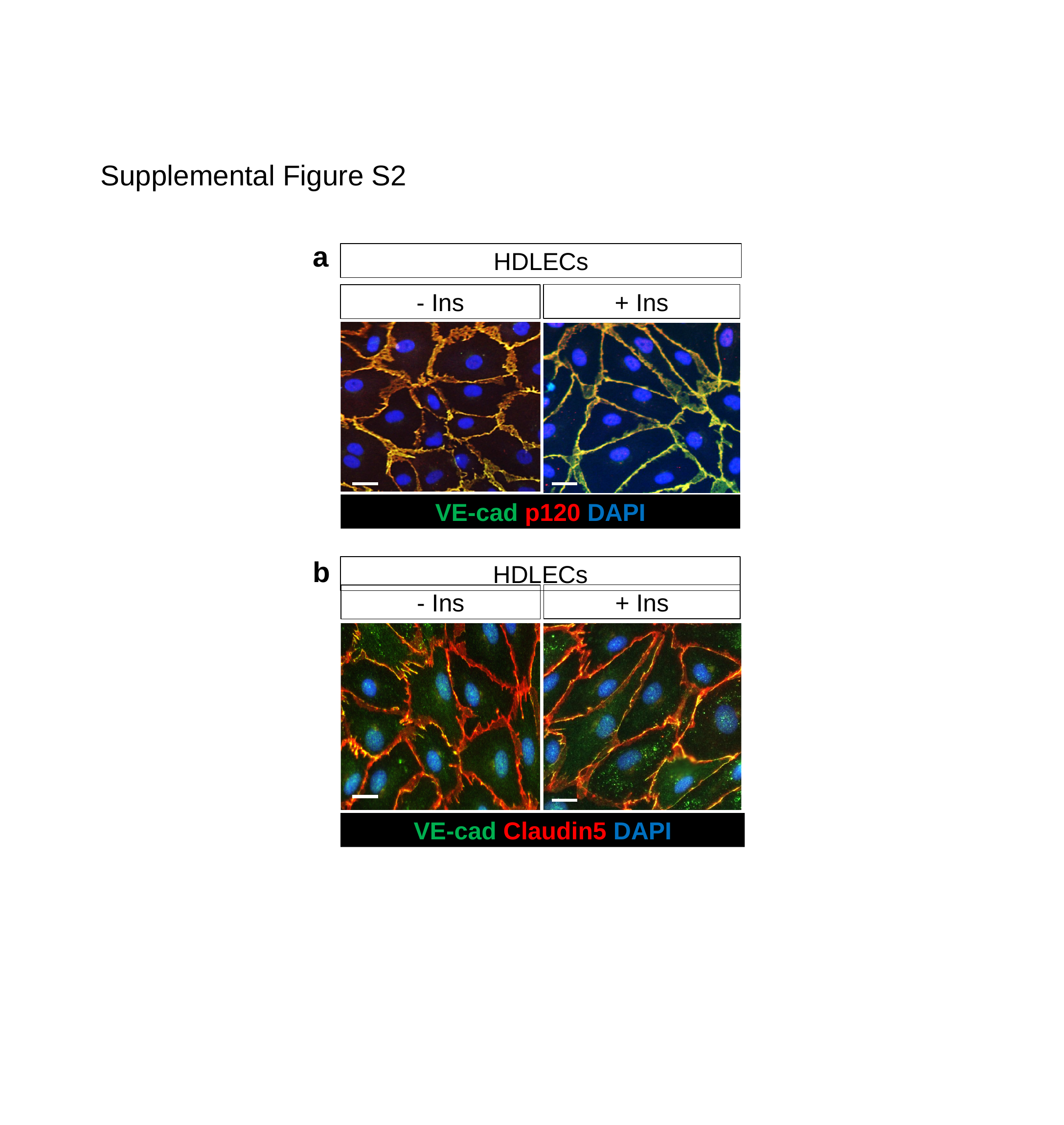

Supplemental Figure S2
a
HDLECs
+ Ins
- Ins
VE-cad p120 DAPI
b
HDLECs
+ Ins
- Ins
VE-cad Claudin5 DAPI

### Slide 3
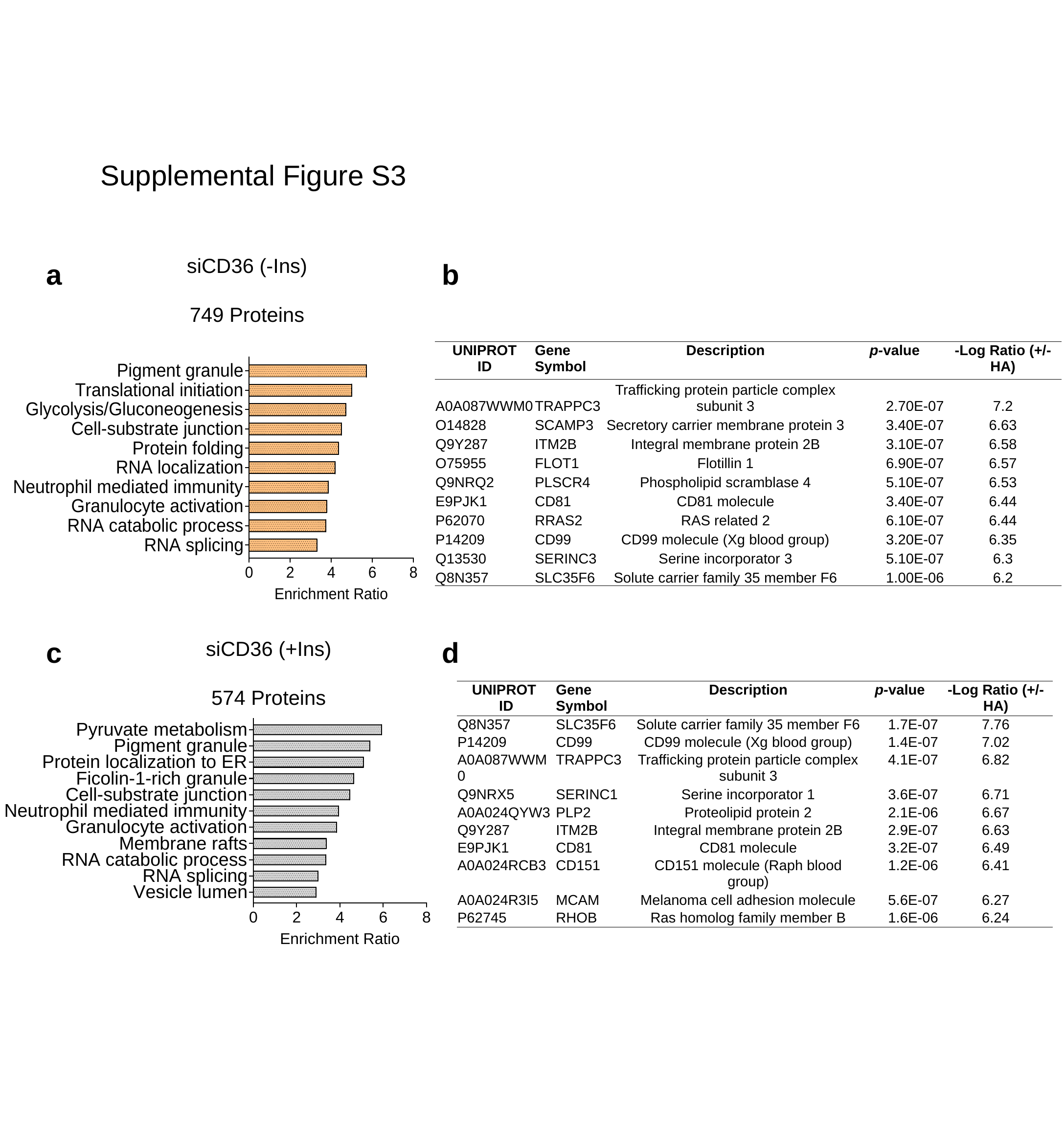

Supplemental Figure S3
siCD36 (-Ins)
749 Proteins
a
b
| UNIPROT ID | Gene Symbol | Description | p-value | -Log Ratio (+/- HA) |
| --- | --- | --- | --- | --- |
| A0A087WWM0 | TRAPPC3 | Trafficking protein particle complex subunit 3 | 2.70E-07 | 7.2 |
| O14828 | SCAMP3 | Secretory carrier membrane protein 3 | 3.40E-07 | 6.63 |
| Q9Y287 | ITM2B | Integral membrane protein 2B | 3.10E-07 | 6.58 |
| O75955 | FLOT1 | Flotillin 1 | 6.90E-07 | 6.57 |
| Q9NRQ2 | PLSCR4 | Phospholipid scramblase 4 | 5.10E-07 | 6.53 |
| E9PJK1 | CD81 | CD81 molecule | 3.40E-07 | 6.44 |
| P62070 | RRAS2 | RAS related 2 | 6.10E-07 | 6.44 |
| P14209 | CD99 | CD99 molecule (Xg blood group) | 3.20E-07 | 6.35 |
| Q13530 | SERINC3 | Serine incorporator 3 | 5.10E-07 | 6.3 |
| Q8N357 | SLC35F6 | Solute carrier family 35 member F6 | 1.00E-06 | 6.2 |
c
d
siCD36 (+Ins)
574 Proteins
| UNIPROT ID | Gene Symbol | Description | p-value | -Log Ratio (+/- HA) |
| --- | --- | --- | --- | --- |
| Q8N357 | SLC35F6 | Solute carrier family 35 member F6 | 1.7E-07 | 7.76 |
| P14209 | CD99 | CD99 molecule (Xg blood group) | 1.4E-07 | 7.02 |
| A0A087WWM0 | TRAPPC3 | Trafficking protein particle complex subunit 3 | 4.1E-07 | 6.82 |
| Q9NRX5 | SERINC1 | Serine incorporator 1 | 3.6E-07 | 6.71 |
| A0A024QYW3 | PLP2 | Proteolipid protein 2 | 2.1E-06 | 6.67 |
| Q9Y287 | ITM2B | Integral membrane protein 2B | 2.9E-07 | 6.63 |
| E9PJK1 | CD81 | CD81 molecule | 3.2E-07 | 6.49 |
| A0A024RCB3 | CD151 | CD151 molecule (Raph blood group) | 1.2E-06 | 6.41 |
| A0A024R3I5 | MCAM | Melanoma cell adhesion molecule | 5.6E-07 | 6.27 |
| P62745 | RHOB | Ras homolog family member B | 1.6E-06 | 6.24 |
